# Lineage tracing reveals asynchronous fate restriction in mouse embryos and stem cell-derived embryo models

**DOI:** 10.64898/2026.10.06.756974

**Authors:** Yunzheng Liu, Jihye Park, Laura Amaya, Sergi Junyent, Xiangkai Sun, Jiabao Sean Xiao, Yuqi Wang, Hao Wang, Yunrui Lu, Shoma Nakagawa, Sung-Eun Park, Wenqi Hu, Shichen Liu, Jiangbo Wei, Hyongbum Henry Kim, Matthew Thomson, Cheng-Wei Ju, Marianne E. Bronner, Junhong Choi, Magdalena Zernicka-Goetz

**Affiliations:** Division of Biology and Biological Engineering, California Institute of Technology, Pasadena, CA 91125, USA; Division of Chemistry and Chemical Engineering, California Institute of Technology, Pasadena, CA, 91125, USA; Developmental Biology Program, Memorial Sloan Kettering Cancer Center, New York City, NY 10065, USA; Department of Pharmacology, Yonsei University College of Medicine, Seoul, Republic of Korea; Division of Physics, Mathematics and Astronomy, California Institute of Technology, Pasadena, CA 91125, USA; Division of Engineering and Applied Science, California Institute of Technology, Pasadena, CA 91125, USA; Department of Chemistry, Department of Biological Sciences, and Cancer Science Institute, National University of Singapore, Singapore 117544, Singapore

## Abstract

How a flexible mammalian embryo gives rise to a reproducible body plan remains unclear, partly because cell lineages cannot be followed continuously in vivo. We combined DNA Typewriter lineage recording with tetraploid complementation to reconstruct high-resolution single-cell genealogies across mouse development and built FateVec, an analysis framework that infers fate-bias dynamics and lineage-associated transcriptional programs from reconstructed phylogenies. Loss of developmental potential proved progressive and asynchronous, with multiple lineages acquiring detectable fate bias before morphological diversification at gastrulation. Within the neural crest, most cells remained multipotent over an extended window, while independent founder clades differed markedly in restriction timing. Extending lineage recording to a stem-cell-derived embryo model revealed that cells with shared transcriptional identities can arise from distinct lineage relationships. We further found embryonic-lineage contributions to extraembryonic-like states, exposing an unexplored lineage plasticity. This lineage-resolved view of embryogenesis reveals that cell identity alone predicts neither developmental ancestry nor the timing of fate restriction.

## Introduction

Mouse development is remarkably robust and flexible. Unlike eutelic organisms such as certain nematodes that develop with a fixed number of cells and invariant lineages (*1*), mammalian embryos can compensate for variation in early cell numbers (*2, 3*), and mouse embryos regulate tissue size after experimental increases in blastocyst cell number (*4*). This developmental plasticity enables powerful experimental paradigms such as embryonic stem cell-tetraploid complementation, in which a small number of genetically engineered mouse embryonic stem cells (mESCs) are injected into tetraploid blastocysts (*5, 6*) that can generate the entire embryo and adult (*7*).

Understanding how a variable early embryo gives rise to a stereotyped organism requires knowledge of the genealogical relationships among cells. These relationships can be represented as cell lineage trees, which define how each cell arises and adopts its fate within the larger context of tissue and organ formation. In the mouse, however, reconstructing the embryonic lineage tree has remained difficult because development proceeds in utero, limiting continuous observation of living embryos and making direct lineage tracking by microscopy challenging.

To overcome this challenge, a variety of powerful fate-mapping and lineage reconstruction methods have been developed. These include recombinase-based systems that generate static genomic barcodes in progenitor cells (*8*), as well as CRISPR-based and newer synthetic recording approaches that continuously edit defined genomic loci to create evolving barcodes over time (*9–12*). Such systems offer the possibility of reconstructing lineage relationships at high resolution, analogous to the way naturally occurring genetic and epigenetic variation can record developmental history (*13–15*). Among these approaches, Prime Editing–based recorders (*16–18*) extend this concept by enabling time-resolved lineage recording at near cell-division resolution, providing a dynamic view of lineage diversification during embryonic development.

Lineage information is also needed to determine whether similar cell identities require comparable developmental histories or alternative differentiation trajectories exist. Stem cell-derived embryo models generate many recognizable embryonic and extraembryonic-like cell types (*19–22*), but whether these identities arise through lineage relationships comparable to those in embryos remains unclear. Moreover, cell identity need not uniquely specify embryonic or extraembryonic ancestry, for example, cells from both origins contribute to the mouse gut endoderm and converge on similar transcriptional identities (*23, 24*). Together, these observations raise two related questions: whether shared cell identities reflect common ancestry, and how readily cells can cross the embryonic–extraembryonic lineage boundary.

Here, we combine Prime Editing-based lineage recording with tetraploid complementation to reconstruct mouse embryonic cell lineages in vivo at high resolution. Using mESCs engineered with the DNA Typewriter lineage recording (*16*), we generated viable embryos and profiled development across seven developmental stages from E7.5 to E14.5, spanning gastrulation, germ layer diversification, and organogenesis. By integrating recorded genealogies with single-cell transcriptomic states, we quantified how developmental potential becomes progressively restricted across embryogenesis. We demonstrate the power of this approach by reconstructing neural crest diversification during migration, revealing inferred progenitors that retain multipotent potential across an extended developmental window. Applying the framework to a stem cell-derived embryo model showed that similar transcriptional identities can arise through distinct lineage histories. We also found evidence that embryonic lineages contribute to extraembryonic-like states in both embryo models and natural mouse embryos. Collectively, the results we present here provide a high-resolution map of mouse embryogenesis and establish a strategy for resolving the ancestry, fate potential, and plasticity of dynamic cell populations.

## Results

### Generation of a DNA Typewriter lineage-recording mESC line

We established a monoclonal mESC line with a DNA Typewriter-based lineage recording system (*16*) including three main components (**fig. S1A**): 1) expression cassettes for doxycycline-inducible PEmax-P2A-mCherry, where PEmax (*28, 29*) expression can be controlled using addition of doxycycline and monitored through mCherry fluorescence, 2) expression cassettes for prime editing guide RNA (pegRNA) that inserts one of 64 possible NNNGGA sequences (InsBC) into a synthetic target site, and 3) expression cassettes for eGFP transcript that contains DNA Tape sequence in the 3’-untranslated region, including 6 sequential editing target sites (6xTape) and a unique 12-bp barcode (TapeBC).

To achieve high-resolution lineage recording, we combined pegRNA- and 6×Tape-cassettes into a single construct for high-copy-number integration (**fig. S1A**), where multiple InsBC and 6×Tape increases the diversity of possible editing patterns and the number of recoverable recording sites, respectively. Amplicon sequencing identified 50 distinct TapeBCs in the final mESC line, each associated with a 6×Tape array. Editing frequencies at all six sites increased following 2 days of doxycycline treatment relative to untreated cells (**fig. S1, B to D**). To characterize baseline editing and subsequent doxycycline-dependent recording, we performed a 12-day bulk genomic DNA time course at three doxycycline concentrations (0, 0.25 μg, and 1 μg ml^−1^). A substantial fraction of Tapes was already edited before induction, but editing depth remained largely stable in untreated cultures. In contrast, doxycycline treatment promoted continued sequential editing, with later Tape positions progressively accumulating edits over time (**fig. S1, E and F**). These results demonstrate continued doxycycline-dependent sequential recording across the Tape array.

### Generation of lineage-recording mouse embryos

We used the engineered Lineage Recording mESC line to generate mouse embryos by tetraploid complementation (*5, 6*). To this end, we recovered embryos at the 2-cell stage, electrofused the two blastomeres to generate tetraploid (4N) cells, which contribute predominantly to extraembryonic tissues, and cultured the resulting embryos to the blastocyst stage. In each experiment, 12 engineered mESCs were injected to 4N blastocysts (**Fig. 1A**). Approximately 20 reconstructed E3.5 embryos were transferred into each E2.5 pseudopregnant surrogate. At the time of embryo transfer, each surrogate was injected with doxycycline to initiate the PEmax expression for recording in utero. After the initial injection, doxycycline level was sustained via feeding throughout the experiment, timing the recording window from around E4.0 to the time of embryo collection and dissection.

**Fig. 1.**
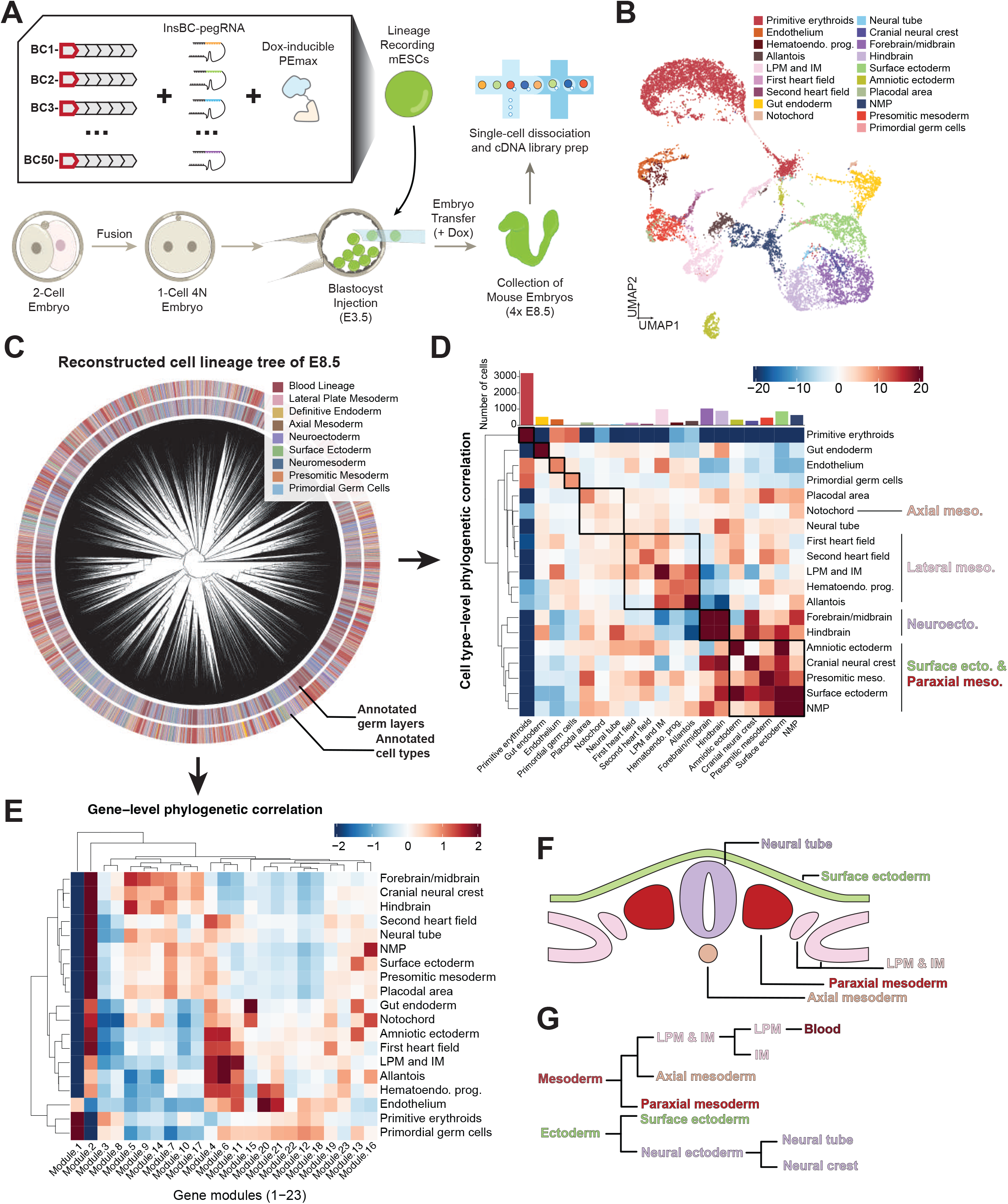
Single-cell lineage reconstruction for mouse E8.5 embryos. **(A)** Schematic of generating lineage recording mouse embryo through tetraploid complementation assay. Mouse embryonic stem cells (mESCs) engineered with dox-inducible lineage recording components (an inducible prime editor, insertion-barcode guides, and 50 six-site DNA Tapes) were injected to the blastocyst of E3.5 4N mouse embryo, contributing to the embryo proper. Upon injection of lineage tracing mESCs, doxycycline is added to promote additional lineage recording. Four resulting mouse embryos were pooled at E8.5 stage, dissociated, and processed for single-cell RNA sequencing with simultaneous recovery of DNA Tape sequences. **(B)** Uniform manifold approximation and projection (UMAP) of single cell transcriptomic data from E8.5 embryos (n = 11,710 cells), annotated with major germ layers and 19 cell types, excluding extraembryonic visceral endoderm (ExVE). **(C)** The single-cell lineage tree reconstructed for E8.5 mouse embryos (n = 10,579 cells with ≥8 recovered edited DNA Tapes). The inner ring is colored as the broad germ layer (colored as in the legend), while the outer ring is colored as the 19 cell types (consistent with the color in **(B)**). **(D)** Cell-type-level phylogenetic cross-correlation matrix computed from the reconstructed lineage tree using the PATH framework. Pairwise correlations reflect the degree of shared lineage history between annotated cell types. Color scale indicates the phylogenetic correlation coefficient (z-scored). Bar chart above the matrix shows the number of cells per cell type. **(E)** Gene-level phylogenetic correlation analysis. Twenty-three co-expression modules identified by Hotspot analysis using lineage-derived cell–cell distances are shown as columns; rows represent annotated cell types. Color indicates mean module score (z-scored across modules within each cell type). Hierarchical clustering of cell types reflects similarity in module-score profiles. **(F)** Cross-sectional cartoon schematic of the non-limb somitic region of the embryo, illustrating the spatial relationships among major ectodermal and mesodermal lineages including neural tube, surface ectoderm, paraxial mesoderm, lateral plate mesoderm and intermediate mesoderm (LPM&IM), and axial mesoderm. **(G)** Developmental hierarchy of mesodermal and ectodermal lineages inferred from phylogenetic correlation analysis, showing the branching relationships among major germ-layer programs from mesoderm and ectoderm to their downstream derivatives.

Embryo development was robust throughout embryogenesis. We collected embryos at seven developmental stages (two at E7.5, four at E8.5, two at E9.5, three at E10.5, two at E11.5, one at E13.5, and three at E14.5), capturing late head fold formation, the embryo turning, neuropore closure, limb and tail bud formation, somitogenesis, and digit and facial development. At each stage, embryos displayed the expected morphology (**fig. S2A**). Tetraploid complementation consistently yielded stage-appropriate embryos, demonstrating the reproducibility of this approach (**fig. S2A**). Green and red reporter fluorescence, together with the recovery of Tape transcripts, confirmed expression of the recorder components in embryos (**fig. S2, B to D**).

At each time point, embryos were dissected according to their developmental stage: E7.5 and E8.5 embryos were processed whole, while embryos from E9.5 to E13.5 were separated into head and body, E14.5 embryos were divided into head, body, and vertebral regions (**Fig. 1A**). For the E8.5 and later collections, whole embryos or stage-matched dissected regions were collected from the same pregnant dam and pooled for single-cell dissociation and droplet-based single-cell RNA sequencing (scRNA-seq). Transcriptomic libraries were prepared using a standard protocol with one modification in the cDNA PCR step to generate a separate amplicon library for DNA Tape sequences (**Methods**).

After filtering out low-quality cells and consolidating technical duplicate records, we recovered 140,592 single-cell transcriptomes across seven developmental stages (**fig. S2, C to E,** and **table S1**). We confirmed continued Tape editing across stages, with increased occupancy at later Tape positions (**fig. S2F**). We first established our analytical pipeline using the E8.5 dataset, then assessed reproducibility in E7.5 embryos, whose smaller cell numbers enabled us to capture a substantial fraction of cells from each embryo, before extending the analysis to later developmental stages.

### Reconstruction of an E8.5 single-cell lineage tree

Using the standard scRNA-seq analysis pipeline, we recovered 13,832 high-quality single-cell transcriptomes from four E8.5 embryos pooled at the time of collection before additional lineage-analysis filters (**Methods**). Extraembryonic visceral endoderm cells were excluded from the primary embryonic lineage analysis and examined separately below. Embryonic cells were annotated into 19 major cell types based on established marker genes (*30*) (**Fig. 1B**). Lineage-recording quality was assessed by quantifying the number of DNA Tape sequences recovered per cell. Because all cells were derived from a monoclonal mESC line, each was expected to carry the same number of DNA Tape copies in the genome, while their capture from transcriptome may vary across cells as previously observed (*16*). The number of recovered DNA Tape molecules was associated with the number of reverse-transcription unique molecular identifiers (RT-UMIs; 16 nucleotides; Pearson r = 0.39; Spearman ρ = 0.31) per cell. The total number of edits correlated more strongly with the number of recovered DNA Tapes (Pearson r = 0.94; Spearman ρ = 0.89), consistent with a contribution of capture depth to variation in recovered lineage information (**fig. S3, A and B**).

Including cells with few recovered DNA Tape molecules risks introducing low-confidence lineage information that could compromise tree reconstruction. To minimize these effects, we excluded cells below a defined recovery threshold. Raising this recovery threshold from 1 to 20 edited DNA Tape molecules progressively decreased the number of retained cells, while minimally affecting cell-type composition (**fig. S3, C and D**). We chose a recovery threshold of at least eight edited DNA Tape molecules, retaining 10,579 cells for lineage-tree reconstruction (**Fig. 1C and fig. S3E**).

### Cell type–level phylogenetic correlation analysis

The reconstructed lineage tree and the matching single-cell transcriptomes contain multiple layers of information for resolving relationships among cell types, the timing of fate restriction, and the regulatory histories associated with differentiation. To interrogate this structure, we first applied the PATH framework to calculate phylogenetic correlation among annotated cell types (*31*). Although originally developed for cancer lineage tracing data, PATH allowed us to calculate phylogenetic cross-correlations and group cell types into broad clusters that matched their developmental origin (**Fig. 1D**). Consistent with the sequential nature of the recorder, incorporating additional editing sites into tree reconstruction progressively increased cell-type clustering, indicating that edits at later sites improved lineage resolution **(fig. S3F)**.

Primitive erythroids were the outlier, exhibiting the highest autocorrelation values, consistent with rapid proliferation and self-renewal within the lineage. This was also evident from their high prevalence and strong clustering within the lineage tree (**Fig. 1C**). Primitive erythroids formed a distinct group, with high autocorrelation followed by cross-correlation to endothelium cell types. On the other hand, ectoderm-derived cell types, such as neural crest, surface ectoderm, and other neuroectodermal populations were largely grouped together. Neuromesodermal progenitors (NMPs) and their paraxial mesoderm derivatives were in this group, reflecting their close relationship to neural lineages. Endoderm-derived gut cells, by contrast, mostly had low cross-correlation values, consistent with a distinct germ layer identity in the sampled population.

### Gene-level phylogenetic correlation analysis

Although PATH analysis recapitulates expected relationships among annotated cell types, its resolution is inherently constrained by the annotations themselves: transcriptionally similar cells with distinct differentiation trajectories may be grouped under the same label. To examine more directly how the lineage tree is reflected at the transcriptome level, we extended the phylogenetic correlation analysis from cell types to individual genes using Hotspot (*32*). Hotspot is a general framework that operates on an arbitrary cell-cell similarity metric. In our case, we defined similarity using lineage-derived distances between cells. Conceptually, using a lineage-based neighborhood graph in Hotspot is closely related to computing phylogenetic correlations for quantitative traits (*e.g.*, gene expression) on a tree, as demonstrated for cell-type traits with PATH (*31*). We chose Hotspot because it offers an efficient, integrated workflow to identify genes showing structured variation across the lineage metric and to group co-varying genes into modules.

We applied Hotspot module discovery to the expressed gene set and identified 23 gene modules (**fig. S4, A and B**, **table S2**). We then calculated module scores across 19 annotated cell types (**Fig. 1E**). This module-level view revealed two complementary patterns. First, a small subset of modules was highly cell-type specific. For example, module 1 showed strong enrichment in primitive erythroid cells, consistent with the strong lineage autocorrelation observed for this cell type in PATH. More generally, when a cell type occupies a tight region of the tree, its marker genes are expected to exhibit elevated within-module concordance. A similar specificity was observed for module 15 in gut cells.

Second, many modules displayed coordinated enrichment across related cell types. Hierarchical clustering of cell types by their module score profiles (**Fig. 1E**) highlighted a major split: excluding a few outliers (*e.g.*, primitive erythroid and gut/endoderm-derived cells), cell types largely partitioned into (i) mesodermal derivatives (including lateral plate mesoderm, intermediate mesoderm, first heart field, second heart field, and axial mesoderm/notochord) and (ii) ectodermal and neuromesodermal derivatives. This division closely mirrors the relationships inferred from lineage-based cell-type phylogenetic correlations alone (**Fig. 1D**). Multiple modules contributed to this separation. For instance, modules 4, 6, and 11 tended to score higher in lateral plate mesoderm derivatives, whereas modules 2, 5, and 9 tended to score higher in neuroectodermal and neural crest lineages. Amniotic ectoderm clustered with mesodermal populations in the module-score heatmap and shared elevated scores for modules 4 and 6 (**Fig. 1E**). This grouping reflects similarity in lineage-associated gene-module profiles, whereas PATH summarizes associations between annotated cell types on the reconstructed lineage tree (**Fig. 1D**). Together, these lineage-associated gene modules support the idea that early lineage origin imposes a strong organizing principle on transcriptional programs, which can manifest as structured relationships among major lineages even without strict dependence on spatial proximity (**Fig. 1, F and G** and **fig. S4C**).

### Lineage-resolved dynamics of fate restriction

Because edits accumulate on DNA Tapes before and during the induced in utero recording window, the reconstructed phylogeny supports clonal tracing and provides a relative lineage-depth coordinate that can be related to developmental progression. We therefore asked whether changes in descendant fate composition along this coordinate could reveal when cell ancestry first becomes preferentially associated with particular terminal fates.

Existing lineage-tree approaches for analyzing fate decisions on single-cell lineage trees provide complementary descriptions of progenitor organization (*33*). First, TarCA estimates effective progenitor numbers from tissue-specific coalescence (*34*). Second, quantitative fate mapping (QFM) infers progenitor-state hierarchies and commitment parameters from lineage barcodes and terminal cell types (*35*). Third, Carta reconstructs discrete potency states (*36*). When applied to our E8.5 reconstructed tree, however, TarCA and Carta did not yield readily interpretable estimates of fate-restriction timings (results from QFM analysis are discussed in the next section). TarCA-derived progenitor numbers showed limited correspondence with established developmental ordering, whereas Carta identified broad candidate progenitor sets that could not be mapped unambiguously to fate-specific restriction depths (**fig. S5, A to C**). This may reflect the noise and reconstruction uncertainty inherent in the experimental lineage tree, rather than limitations of these methods in their intended applications. We therefore developed a complementary framework tailored to extracting continuous fate-bias dynamics from a fixed reconstructed phylogeny.

To extract fate restriction dynamics from our reconstructed phylogeny, we therefore developed FateVec, a lineage-grounded framework that represents each node as a quantitative fate state vector and estimates fate-bias timing from its evolution along lineage tree depth (**Fig. 2A** and **fig. S6, A to H**). In the FateVec formulation, let *K* denote the number of annotated terminal fates (19 annotated cell types at E8.5). We assign each terminal cell (tip of the tree) a one-hot fate vector v∈{0,1}*^K^* according to its cell-type annotation. For each internal node *p*, a fate-specific vector *V_p_ = (V_p_^1^,…, V_p_^K^)* is computed by recursively aggregating across children: for a binary split into children *m* and *n*, *V_p_* and the descendant count vector *N_p_* are updated as shown in **Fig. 2A**:

**Fig. 2.**
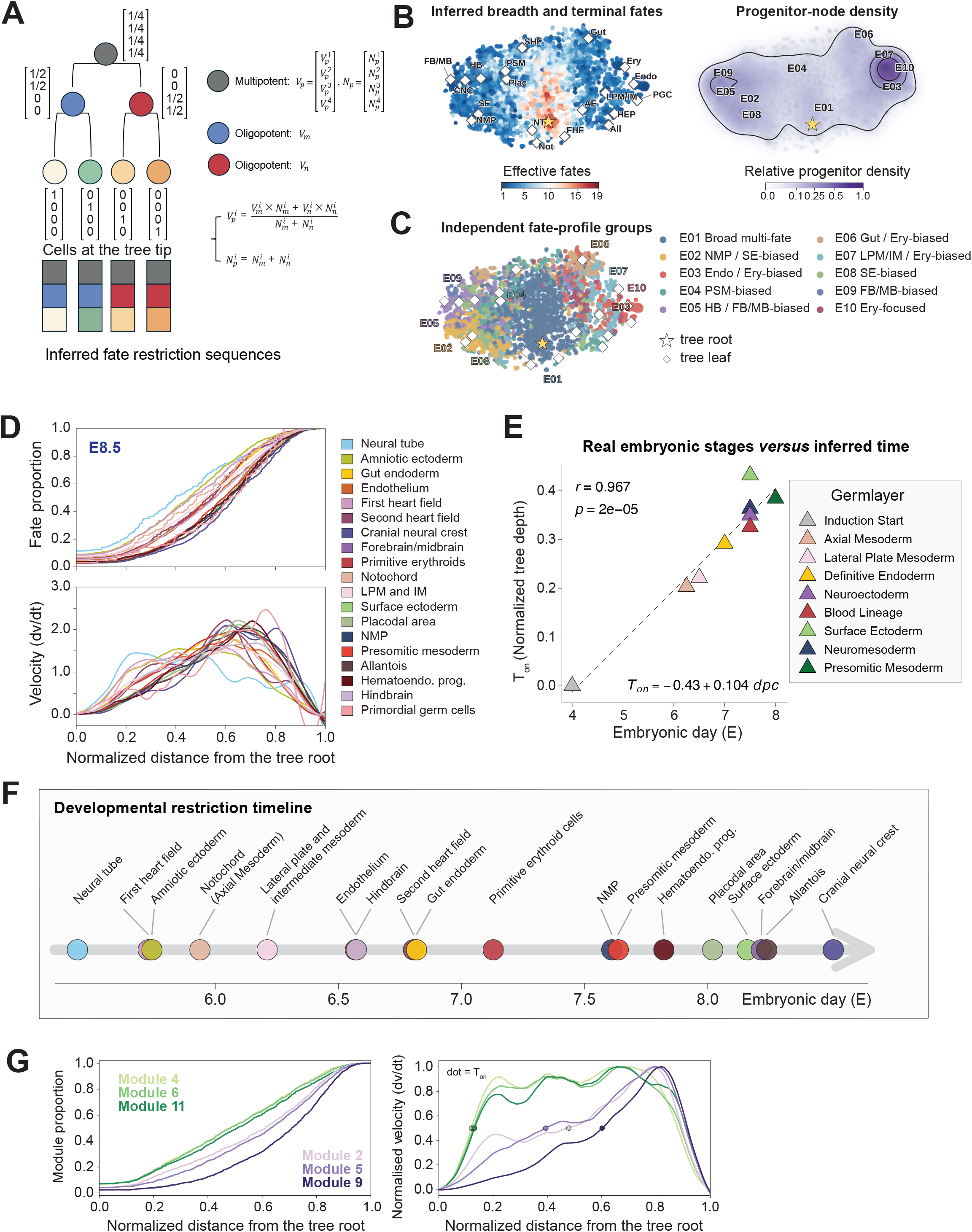
Inferring the gene module activation and fate restriction timings. (A) Schematic illustrating the FateVec framework applied to a reconstructed lineage tree. Terminal cells (tips) are encoded as one-hot fate vectors reflecting their annotated cell-type identity. Internal nodes represent ancestral progenitors whose fate profiles are inferred by bottom-up aggregation from their descendants (**Methods**). At each ancestral node, each fate component is averaged across child nodes, weighted by the number of descendants of that fate, and the resulting vector is normalized across fates. Node colors and stacked bars illustrate ancestral fate profiles along root-to-tip paths, tracing the emergence of fate-specific lineage organization. (B) t-SNE embedding of lineage-tree nodes based on their 19-component FateVec profiles. Left, effective fate breadth, defined as the exponential Shannon entropy of the normalized FateVec profile; higher values indicate broader fate associations. The star marks the root; diamonds indicate terminal cell-type profiles. Right, relative density of internal nodes in the same embedding. (C) Ten groups of internal-node fate profiles (E01–E10), identified by clustering their FateVec profiles and displayed in the same embedding as in **(B)**. Colors distinguish groups with similar ancestral fate associations. NMP: Neuromesodermal progenitors; SE: Surface Ectoderm; Endo: Endothelium; Ery: Primitive Erythroids; PSM: Presomitic Mesoderm; HB/FB/MB: Hind-/Fore-/Mid-brain; LPM and IM: Lateral Plate Mesoderm and Intermediate Mesoderm. (D) Fate restriction dynamics for all 19 annotated cell types in E8.5 embryos. Upper panel: mean FateVec trajectories *V̄ᵏ(t)* as a function of normalized distance from the tree root *t*, with each fate component averaged at each depth along root-to-tip paths ending in cells of that type. Lower panel: derivatives *dV̄ᵏ(t)/dt* for each cell type, highlighting intervals of increasing fate specificity. Tree depth is normalized from 0 (root) to 1 (tips). (E) Calibration of FateVec restriction-onset depth to developmental chronology. *T_on_* is defined as the first depth at which the smoothed derivative of the baseline-normalized FateVec trajectory reaches half its maximum. For eight broad lineages, *T_on_* is plotted against assigned developmental reference ages, with an additional reference anchor assigned to E4.0 at normalized depth 0. The dashed line shows the linear calibration across all nine points, used to convert *T_on_* estimates to approximate embryonic ages. (F) Calibrated restriction-onset timeline for E8.5 cell types, projected onto calibrated embryonic time using the linear model in **(E)**. Each circle represents a cell type, colored as in **Fig. 1B**, and positioned at its inferred restriction-onset time (*T_on_*) along the embryonic day axis. The primordial germ-cell estimate falls outside the plotted range, and the remaining 18 cell types are shown. (G) Gene module activation dynamics for six lineage-associated co-expression modules inferred from the E8.5 lineage tree: mesodermal modules M4, M6, and M11 (green) and neural modules M2, M5, and M9 (purple). Left: mean FateVec module trajectories as a function of distance from the tree root. Right: peak-normalized derivatives; dots mark *T_on_*.

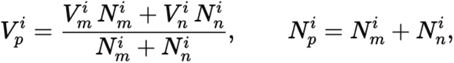

Under this construction, ancestral nodes with multiple observed descendant fates showed broadly distributed fate-bias vectors *V_p_*, whereas more restricted nodes showed weights increasingly concentrated toward a single fate component. To visualize these patterns in the E8.5 dataset, we embedded the 19-dimensional fate-bias vectors of internal nodes using t-SNE. We summarized effective fate breadth as the exponential Shannon entropy of each node’s normalized FateVec profile, with higher values indicating broader inferred fate associations (**Fig. 2B**; **Methods**). Neuroectodermal and neural crest fates, including fore/mid/hindbrain (FB/MB/HB) and cranial neural crest (CNC), occupied neighboring regions of the embedding together with their associated ancestral nodes (**Fig. 2B**). We then clustered ancestral nodes by their fate-bias profiles and traced backward from terminal fates to identify the putative progenitor groups associated with each lineage. Primitive erythroid–associated nodes included a distinct erythroid-focused group whereas neural crest cells traced back to a more heterogeneous set of progenitor states (**Fig. 2C**; **fig. S6I**).

We next quantified how fate bias toward each terminal fate emerged along reconstructed lineage history. For each annotated fate *k*, we traced the corresponding fate-bias component *V^k^(t)* along root-to-tip trajectories terminating in that fate. We averaged these trajectories to obtain a mean fate-bias profile and calculated its derivative, *dV^k^(t)/dt*, as the fate-bias velocity. We defined *T_on_* as the leading half-maximum crossing on the rising flank of this velocity profile, before its peak (**Methods**). This operational onset metric marks the beginning of fate-bias acquisition in each cell type category.

### Benchmarking FateVec using lineage reconstruction simulations

Before applying FateVec to the reconstructed embryonic phylogeny, we benchmarked it using simulated developmental programs with known fate-restriction depths. We simulated an eight-fate tree with highly unequal terminal abundances, generated DNA Typewriter recording outcomes, reconstructed the lineage tree, and applied FateVec without providing the programmed depths. FateVec recovered the correct restriction order despite unequal fate abundance (**fig. S7, A to C**).

We then modeled three major sources of recorder noise: (1) nonlinear changes in editing activity over time (**fig. S7D**); (2) heritable heterogeneity in editing activity among descendant lineages (**fig. S7E**); (3) biased rather than uniform barcode insertion outcomes (**fig. S7F**). We also modeled Tape dropout across a range comparable to that observed experimentally (**fig. S7G**). To reduce these effects, we calculated a relative lineage distance by normalizing pairwise differences between cells to their total editing load (**fig. S7H**). Nonlinear editing rates and skewed insertion outcomes had little effect on fate ordering, whereas heterogeneous inheritance of editing activity was the main source of uncertainty (**fig. S7I**). Relative normalization substantially improved performance under this condition, and fate ordering remained robust to Tape dropout (*ρ* > 0.90). Subsampling further preserved fate ordering after relative normalization (**fig. S7J**). Although recorder noise could shift inferred absolute depths, relative fate ordering remained stable and inferred depths remained approximately linear with programmed depths, supporting empirical calibration of relative lineage depth using developmental landmarks (**fig. S8**).

Finally, we examined cases in which terminal fates did not map cleanly onto lineage topology. We compared FateVec with QFM because the two methods provide complementary information. QFM infers progenitor-state hierarchies (*35*), whereas FateVec estimates the relative timing of individual fate restrictions. When one cell type arose from two independent progenitor populations, FateVec retained the programmed restriction order, while QFM recovered six of seven programmed progenitor states (**fig. S9A**). Relabeling 10% of cells into or out of this cell type yielded FateVec rank correlations of 0.98 ± 0.02 and 0.97 ± 0.03, respectively; QFM recovered an average of 5.6 and 4.8 of seven programmed states (**fig. S9, B and C**). Together, these simulations show that FateVec robustly recovers relative fate-restriction order despite simulated recorder noise, incomplete sampling, lineage convergence, and moderate annotation error.

### Defining fate-restriction timing from vector dynamics

Applying FateVec to the E8.5 phylogeny, we traced the fate-bias component *V^k^(t)* for each annotated terminal fate *k* along normalized distance *t* from the lineage tree root. For each cell type *k*, we averaged *V^k^(t)* across root-to-tip paths ending in cells of that type (**Fig. 2D**, top). These averaged trajectories showed a gradual increase in fate bias, consistent with progressive narrowing of the observed descendant-fate composition. Although the trajectories overlapped extensively, they differed in the timing and rate of fate-bias acquisition.

We next differentiated the averaged trajectories to obtain a commitment velocity, *dV_k_(t)/dt*, which captures the rate of fate-bias acquisition (**Fig. 2D**, bottom). The resulting velocity profiles differed in onset position and width, revealing variation in how rapidly and over what interval fate bias increased across lineages. We summarized these dynamics using *T_on_*, defined as the leading half-maximum crossing of each velocity profile (**Methods**). This provided a common metric for comparing the relative onset of fate-bias acquisition across lineages (**table S3**).

To coordinate these lineage-based onset estimates to embryonic development, we compared FateVec-inferred onset *T_on_* with canonical embryological timelines for major germ-layer programs and representative lineages with well-established onset/commitment windows (*30, 37, 38*). Specifically, we focused on axial and lateral plate mesoderm, definitive endoderm, neuroectoderm, blood lineages, surface ectoderm, neuromesoderm, and presomitic mesoderm (**table S4**). FateVec timings closely tracked the embryonic stage, exhibiting a strong linear relationship between embryonic day and the inferred onset estimates *T_on_* (Pearson *r* = 0.967, P = 2 × 10^−5^; **Fig. 2E**). To assess the robustness of this correlation, we applied label-permutation and cell-subsampling analyses. The observed agreement exceeded the null expectation from 1,000 permutations of terminal cell-type labels that preserved cell-type abundance, both with and without the induction-start anchor (observed r = 0.967 and 0.903; empirical P = 0.002 and 0.001, respectively; **fig. S10, A and B**). We also reran FateVec on 50 subsampled trees, each reconstructed from a random 80% subset of cells sampled without replacement. The corresponding correlations were 0.950 ± 0.033 and 0.858 ± 0.099 (mean ± SD), and the mean timing estimates remained strongly correlated with the developmental reference (**fig. S10, C to E**).

We note that *T_on_* marks the onset of rapid descendant-fate bias along the reconstructed tree, while the PATH analysis summarizes lineage relationships among terminal cell types. This lead to high autocorrelation with relatively weak cross-correlation with most other cell types, as in the case of gut endoderm (**Fig. 1D**), with moderate *T_on_* value (9th out of 19 cell types; **Fig. 2, E and F**). On the other hand, cell types can have early *T_on_* values (e.g., axial mesoderm and lateral plate mesoderm with 4th and 5th lowest *T_on_* values, respectively) with high cross-correlations with other cell types. Strong terminal lineage isolation inferred from PATH analysis therefore need not coincide with the earliest onset of fate bias measured through FateVec.

Together, these analyses established that (i) FateVec yields lineage-specific commitment trajectories whose overlapping fate-bias trajectories differ in the timing and rate of fate-bias acquisition (**Fig. 2, D and E**), and (ii) the inferred restriction times recapitulate the expected developmental ordering across germ-layer programs (**Fig. 2F**), supporting the use of the reconstructed phylogeny as a relative lineage coordinate that can be empirically calibrated to approximate developmental chronology.

### Sequential gene module activation accompanies early fate bias

Calibrating lineage tree depth to embryonic time (**Fig. 2F** and **table S4**) enabled us to reinterpret the fate-bias vector dynamics on the reconstructed phylogeny in a developmental framework. When we projected the inferred restriction times back onto the developmental timeline, we observed that many fate trajectories reached their fate bias timing calibrated by *T_on_* well before E7.0, indicating that lineage-specific fate bias is established prior to, rather than only during, overt morphological diversification. This inference rests on the strong linear correspondence between calibrated tree depth and known developmental landmarks (**Fig. 2E and fig. S10**), though direct profiling at pre-gastrulation stages would provide further validation. Nonetheless, this finding extends classical fate maps and molecular descriptions of gastrulation (*30, 37–41*), and supports recent evidence that transcriptional and regulatory heterogeneity can precede clear anatomical landmarks (*8, 42*). Together, these results suggest that fate restriction begins earlier than generally appreciated and unfolds as a gradual, asynchronous process rather than as a single coordinated transition.

To interpret these early restriction events, we turned to lineage-specific gene modules. Hotspot analysis on the lineage metric identified gene modules whose activities segregate canonical lineages beyond what is captured by transcription-only annotation (**Fig. 1E** and **fig. S4**). For example, neuroectoderm-associated populations such as forebrain/midbrain showed preferential activity of modules 2, 5 and 9, whereas lateral plate mesoderm derivatives were enriched for modules 4, 6, and 11.

To understand the temporal sequence of this combinatorial structure, we applied the same vector-propagation framework used for fate bias to these gene modules, inferring internal-node module bias by bottom-up aggregation (**Methods**). Along normalized tree depth, the peak-normalized derivatives of mesoderm-associated modules 4, 6, and 11 reached their leading half-maximum crossings earlier than those of neuroectoderm-associated modules 2, 5, and 9 (**Fig. 2G**). These module-bias trajectories describe lineage-associated gene programs, whereas the cell-type trajectories summarize bias toward individual terminal identities (**Fig. 2, D and F**). The early onset of neural-tube fate bias therefore need not coincide with the onset of every neuroectoderm-associated module; the two analyses summarize different features of the reconstructed lineage tree.

These distinct module enrichments suggest that the successive waves of fate commitment are driven by sequential activation of specific regulatory programs. Early-activated modules may encompass regulators of pluripotency exit and germ layer allocation, whereas later modules likely reflect transcriptional networks driving lineage-specific differentiation and morphogenesis (**fig. S11, A and B**). To identify candidate regulators driving this sequence, we examined transcription factors enriched in these modules. *Sp1*, which is required for early embryonic development (*43, 44*), emerged as a candidate regulator for early-primed populations in modules 4, 6, and 11 (**fig. S11C**), whereas Fox-family regulators were preferentially associated with late-activating modules 2, 5, and 9 (**fig. S11C**). To validate *Sp1* as an early regulator of fate decision in epiblasts, we analyzed single cell transcriptomic data from E5.25 embryos (**fig. S11D**) (*42*). Consistent with a role for *Sp1* in early fate decisions (*45*), we observed a high level of heterogeneity in *Sp1* expression across epiblasts, potentially underlying preferential activation of its downstream targets (**fig. S11, E to G**). In contrast, other well-known transcription factors did not exhibit comparable levels of heterogeneity at this stage (**fig. S11, H and I**).

Collectively, these findings support an emerging model of asynchronous development, characterized by a progressive rather than abrupt loss of pluripotency (*46, 47*). Our data suggest that upstream transcriptional heterogeneity, or regulatory symmetry breaking, is established as early as E5.25, initiating molecular diversification well before the onset of gastrulation.

### Reproducibility across embryos and clades

To assess embryo-to-embryo reproducibility, we analyzed 6,107 and 11,083 transcriptomes from the retained populations of two E7.5 embryos, of which 5,873 and 8,922 cells, respectively, were included in lineage-tree reconstruction (**fig. S12A**). After matching cell numbers by cell type, between-type PATH associations and within-type autocorrelations were highly concordant between embryos (Spearman *ρ* = 0.815 and 0.976, respectively; **fig. S12, B to D**). Both embryos showed preferential coupling of notochord with definitive endoderm relative to paraxial mesoderm, resolving lineage relationships beyond broad germ-layer assignments (**fig. S12C**). The estimated onset of fate-bias acquisition (*T_on_*) also showed concordant ordering across the seven cell types represented by more than 200 cells in each embryo (Spearman *ρ* = 0.893; **fig. S12, E and F**). These comparisons support reproducibility of both lineage relationships and the relative ordering of fate-bias acquisition.

Because DNA Tape recordings retain both clonal information and relative lineage depth, we next asked whether fate-bias ordering was shared across inferred clades. We pooled the two E7.5 datasets and grouped cells using first-site insertion patterns without supplying embryo labels (**fig. S13, A and B**). Revealing embryo identities afterward showed that all ten inferred clusters were nearly embryo-specific, with purities of 99.66–100%. Thus, marker-based grouping largely preserved embryo-of-origin separation even when embryo identity was withheld from the analysis.

We then compared clade-specific *T_on_* rankings for shared cell types. Pairwise correlations were generally positive both within and between embryos, although agreement was lower between embryos (median Spearman ρ = 0.39 and 0.57 within embryos 1 and 2, respectively, versus 0.31 between embryos, P = 0.024; **fig. S13, C to E**). These results indicate that fate-bias ordering recurs across multiple clades, while also revealing variation in the degree of agreement among clade pairs.

We extended this analysis to the pooled E8.5 dataset, identifying 14 inferred clades from first-site insertion patterns (**fig. S14, A to D**). Despite differences in clade size and lineage composition, the relative ordering of fate-bias onset was generally concordant for shared fates across the ten clades included in the timing comparisons (**fig. S14, E to G**). Across 45 clade-pair comparisons, the observed correlations exceeded the within-clade label-permutation null (P < 0.005; **fig. S14H**), supporting nonrandom concordance in fate-bias ordering across clades. Together, these analyses support reproducibility at two levels of individual embryos and inferred clades.

### Reconstructing lineage trees through mouse organogenesis

We extended our lineage tree analysis to the later-staged mouse embryos, collected across E9.5 to E14.5 during mouse organogenesis (**Fig. 3A**). These later-stage (E9.5 to E14.5) collections yielded 103,757 single-cell transcriptomes, which we integrated with the E7.5 and E8.5 datasets to generate a cross-stage atlas (**Fig. 3, B and C**). We assigned the later-stage cells to 17 major cell types (**Fig. 3B**) and retained those with at least eight edited DNA Tapes for lineage reconstruction (**Fig. 3C**). These cells capture the differentiation of injected mESCs as epiblasts into diverse tissues, including the maturing brain, beating heart, skin, somite, limb, cartilage, and other organs (**Fig. 3D**). From the 53,881 cells that passed this filter, we reconstructed five cell lineage trees from E9.5 to E14.5 (**Fig. 3E**).

**Fig. 3.**
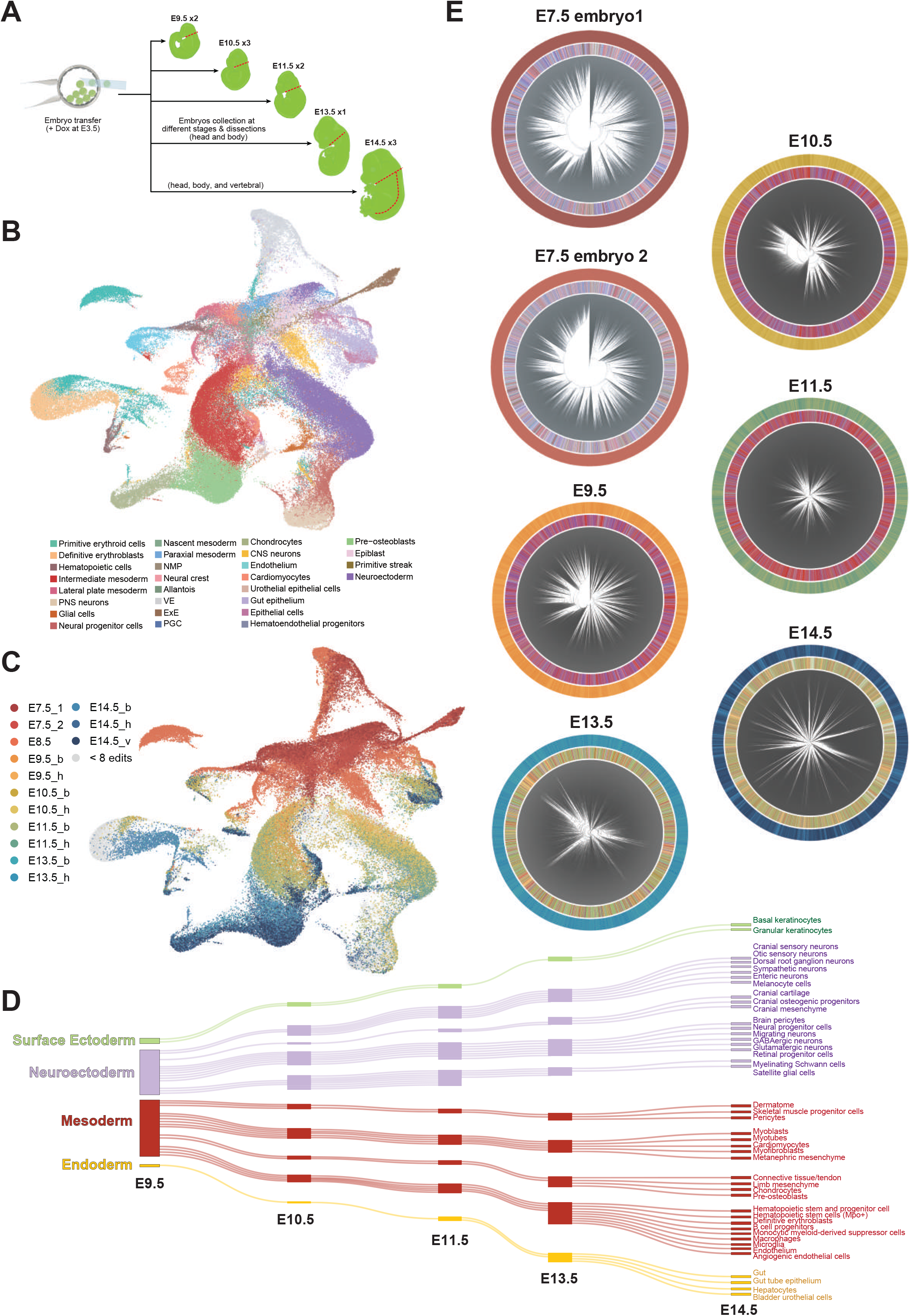
Single-cell lineage reconstruction of mouse organogenesis. **(A)** Schematic of embryo dissection strategies and collection time points for lineage-recording embryos from E9.5 to E14.5 after Dox induction at E3.5. Embryos at E9.5–E13.5 were separated into head and body fractions; E14.5 embryos were further divided into head, body, and vertebral fractions prior to single-cell dissociation and scRNA-seq. **(B)** UMAP of transcriptomes from all collected embryos (E7.5–E14.5), colored by broad cell-type annotation. **(C)** UMAP as in **(B)**, colored by sample of origin. Sample labels indicate embryonic stage and dissection region (e.g., E14.5_v: E14.5 vertebral fraction; E11.5_h: E11.5 head fraction; E10.5_b: E10.5 body fraction), with E7.5_1 and E7.5_2 denoting the two independently processed embryos. Cells with fewer than 8 recovered edited DNA Tapes are shown in gray and excluded from lineage analysis. **(D)** Directed tree graph visualized as a Sankey diagram illustrating cell-type relationships from E9.5 to E14.5. Each node corresponds to a cell type at each developmental stage, arranged by time points along the horizontal axis, and links connect nodes sharing lineage annotations across successive stages. Link and node thickness is proportional to log₂-transformed predicted cell numbers, obtained by scaling estimated total embryonic cell numbers by the observed cell-type proportions (**Methods**). **(E)** Reconstructed single-cell lineage trees for the two independently processed E7.5 embryos and the E9.5, E10.5, E11.5, E13.5, and E14.5 collections, displayed in circular layout. Each leaf represents a single cell. The inner ring is colored by cell-type annotation as in **(B)**; the outer ring is colored by sample of origin as in **(C)**.

We next integrated the reconstructed trees with gene-expression profiles to examine developmental relationships among cell types and identify lineage-associated transcriptional programs (**fig. S15**). Cell-type pairs within the same germ layer showed stronger phylogenetic correlations than pairs from different germ layers, with significant germ-layer separation at every stage from E8.5 to E14.5 (**fig. S16A**). From E9.5 to E11.5, cell types clustered largely according to germ-layer origin, with ectoderm separating into neuroectodermal and surface ectodermal populations, and mesoderm separating into blood progenitors and other mesodermal derivatives (**fig. S15, B to D**). At later stages, E13.5 and E14.5, broad phylogenetic relationships became harder to resolve, likely reflecting both rapid embryonic growth and increasing cell-type complexity (**fig. S15, E and F**). Embryonic cell numbers are estimated to increase approximately 13-fold from E9.5 to E11.5 and a further 8-fold from E11.5 to E14.5 (*41*), while broad populations such as neurons diversify into finer subtypes distinguished by anatomical position and maturation state.

We then compared lineage-associated gene modules across stages to identify recurrent and stage-dependent transcriptional programs. Modules associated with ribonucleoprotein biogenesis were detected throughout E7.5–E14.5, whereas enrichment for translation and posterior Hox patterning was concentrated from E7.5 to E11.5. Extracellular matrix organization and epithelial development were also represented in the later-stage modules (**fig. S16B**). Comparing module gene membership revealed overlapping gene sets across successive stages, indicating cross-stage recurrence of lineage-associated programs despite changes in module size and composition (**fig. S16C**). Thus, the module analysis complemented cell-type-level relationships by revealing both shared and stage-dependent features of lineage-associated gene expression. We also compared cell-type composition between cells with at least one recovered Tape and those with at least 8 edited Tape recovered for lineage reconstruction (**fig. S16D**). We confirmed composition remained similar at E8.5–E10.5 but also observed shifted at later stages, may partly reflecting variation in Tape recovery across cell types.

Together, these analyses establish a lineage-resolved atlas (www.mouse-it.com) that integrates single-cell transcriptomic profiles with lineage reconstructions across mouse development, from gastrulation-associated lineage segregation to organogenesis-stage tissue diversification. This resource enables both global comparisons of developmental relationships and focused analyses of specific multipotent lineages.

### Early neural crest diversification

The neural crest is a migratory cell population that has a remarkably broad developmental potential, with the ability to form cell types as diverse as neurons and glia of the peripheral nervous system, cartilage and bone of the face, pigment cells of the skin, and cells of the cardiac outflow tract (**Fig. 4A)** (*25, 26, 48–50*). The neural crest originates within the dorsal neural tube and represents a transient, multipotent stem cell population within the embryo. Premigratory neural crest cells then undergo an epithelial-to-mesenchymal transition (EMT) to leave the neural tube and migrate throughout the periphery prior to differentiating into numerous cell types in diverse locations (*25, 50*).

**Fig. 4.**
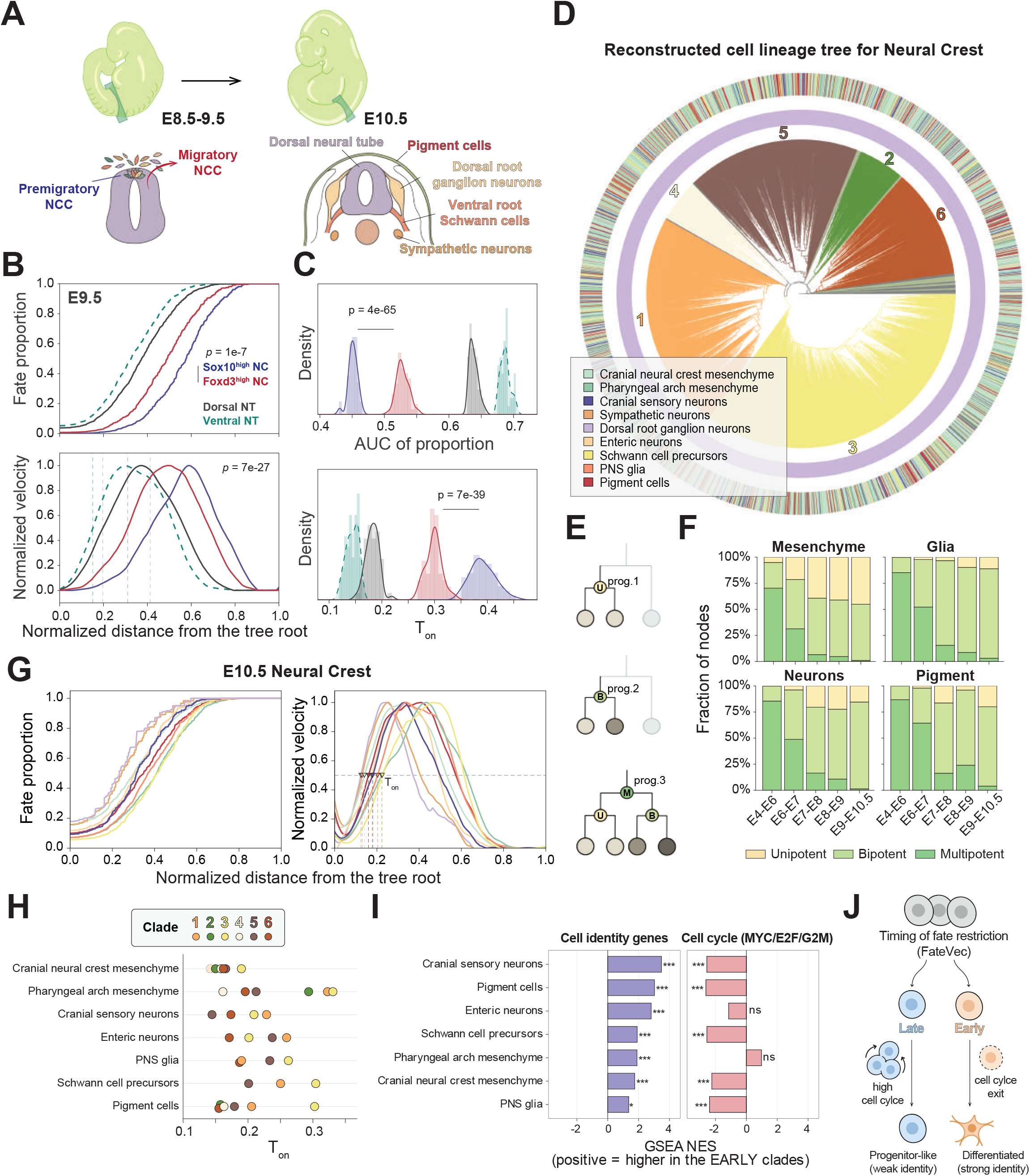
Regulatory kinetics of neural crest diversification. (A) Cartoon schematic of neural crest development between E8.5 and E10.5. Neural crest cells delaminate from the dorsal neural tube and migrate throughout the embryo. By E10.5, they give rise to diverse derivatives in the periphery, including dorsal root ganglion neurons, sympathetic neurons, Schwann cell precursors, and pigment cells. (B) FateVec trajectories (Upper panel) and peak-normalized derivatives (Lower panel) for E9.5 Foxd3^high^ and Sox10^high^ neural crest cells (n = 155 and 130 tree tips), with dorsal and ventral neural tube populations as references. P values compare per-tip trajectory areas for the two neural crest populations by two-sided Mann–Whitney tests. (C) Distributions of FateVec trajectory area (Upper) and *T_on_* (Lower) across 50 tree reconstructions, each using an 80% cell subsample without replacement. Colors correspond to the populations in **(B)**. P values compare Foxd3^high^ and Sox10^high^ neural crest populations by Welch’s tests on resampled estimates. (D) Reconstructed single-cell lineage tree for neural crest-associated cells from E10.5 embryos. Six major clades (numbered 1–6) are highlighted in distinct branch colors. The inner ring indicates broad lineage identity; the outer ring indicates the annotated neural crest-derived cell types. (E) Schematics describing progenitor nodes with different observed fate breadth (U, unipotent; B, bipotent; M, multipotent). Nodes in reconstructed trees are classified according to whether their observed descendants represent one, two, or at least three of the four major derivative classes: mesenchyme, neurons, glia, and pigment cells. (F) The proportion of unipotent, bipotent, and multipotent progenitor nodes, as defined in **(E)**, across four major neural crest-derived cell classes over five developmental intervals. Mesenchyme: Cranial neural crest mesenchyme and Pharyngeal arch mesenchyme. Neurons: Cranial sensory, Dorsal root ganglion, Sympathetic, and Enteric neurons. Glia: PNS glia and Schwann cell precursors. Pigment: Pigment cells. Node dates were estimated by least-squares dating (LSD), with root and collection-stage anchors and a cell-count ceiling (**Methods**). (G) Fate restriction dynamics calculated for all neural crest-derived cell types inferred from the E10.5 lineage tree. Left: mean FateVec trajectories V̄ᵏ(t) as a function of lineage tree depth *t*. Right: peak-normalized derivatives; markers indicate *T_on_*. Cell types are colored as in **(D)**. (H) Restriction-onset timing (*T_on_*) for annotated neural crest-derived cell types, shown separately for each of the six major clades. (I) GSEA compares cells from early- and late-restricting clades within each indicated neural crest-derived cell type, with early and late referring to restriction-onset timing (*T_on_*). Left, cell-type identity signatures. Right, MYC-, E2F-, and G2/M-associated proliferation programs. Bars show normalized enrichment scores (NES); positive values indicate enrichment in early-restricting clades. (J) Proposed relationship between restriction-onset timing, differentiated identity, and proliferation.

Given this incredible diversity, a long-standing question in the neural crest field is the timing of fate restrictions in neural crest cells and when they are committing to a particular cell type. This has been controversial with some authors suggesting early restriction to a particular fate (*51–55*), whereas others have argued for prolonged multipotency (*27, 56–60*). Resolving this has been difficult because neural crest cells disperse widely and intermingle during migration, obscuring clonal relationships and confounding attempts to align early states with later fates. By applying our DNA Typewriter-based lineage tracing *in vivo*, we set out to address this question by resolving clonal relationships and fate-restriction timings for focused populations of neural crest-derived cells.

We first analysed the E9.5 single cell data set with matched phylogeny, in which two neural crest cell types, premigratory (Foxd3^high^) neural crest cells and migratory (Sox10^high^) neural crest cells, emerge from the dorsal neural tube. We include dorsal and ventral neural tube as reference as well. FateVec analysis of annotated cell types revealed a transition from neural tube to premigratory neural crest cells with a following transition from premigratory to migratory neural crest (**Fig. 4, B and C**), consistent with previously observed migration of neural crest cells *in vivo* (*26*).

We next investigated how neural crest cell proliferation and differentiation give rise to neuronal and glial derivatives by reconstructing a neural crest-specific lineage tree at E10.5 (**Fig. 4D** and **fig. S17, A to C**). We defined each node within the tree (representing neural crest progenitors) as either unipotent, bipotent, or multipotent. Unipotent cells were defined as forming only a single derivative, such as mesenchyme, glia, neuron, or pigment (**Fig. 4E**). Bipotent cells gave rise to two of those categories, whereas multipotent progenitors formed three or more. We then used least-squares dating (LSD) to estimate ancestral node times and place these fate contributions on a developmental timescale (*61, 62*). Bipotent and multipotent nodes were detected in calibrated time bins extending to E10.5, with distributions that varied among derivative classes (**Fig. 4F** and **fig. S17D**). Quantification of fate-restriction timing across neural crest derivatives revealed that neuronal lineages undergo fate restriction earlier than glial precursors, such as Schwann cell precursors (SCPs) (**Fig. 4G**). This result suggests that SCPs retain a prolonged multipotent window, consistent with previous observations that SCPs can contribute to multiple neural crest derivatives beyond Schwann cells (*63, 64*). We further quantified the specific multipotent and bipotent combinations represented among inferred nodes (**fig. S17E**).

### Heterogeneity across founder epiblasts couples fate restriction to cell state

We next examined whether fate restriction occurs synchronously across major clades, defined operationally from lineage-recording patterns (**Fig. 4D**). These clades contributed to overlapping neural crest derivatives but differed in their relative contributions (**fig. S18, A to D**). We then compared the relative timing of fate restriction, summarized by *T_on_*, across clades contributing to the same annotated derivative. Despite sharing the E10.5 collection stage and derivative identity, these clades differed in their lineage-inferred onsets of rapid fate-bias acquisition, revealing asynchronous fate restriction within individual derivative classes (**Fig. 4H**).

We next examined transcriptional profiles for additional evidence of this lineage-inferred asynchrony. We first defined a set of marker genes for each cell type (**fig. S18E**). We found that earlier-restricted clades were enriched for cell-type identity genes across the derivatives examined, whereas later-restricted clades generally showed stronger enrichment of proliferation-associated programs (**Fig. 4I** and **fig. S18F**). Across clade–derivative pairs, we observed that earlier *T_on_* was strongly associated with higher cell-identity scores (ρ = −0.66, P = 3.3 × 10⁻⁵) and lower proliferation-program scores (ρ = 0.52, P = 0.002; **fig. S18G**). Thus, the transcriptional differences were concordant with the lineage-inferred temporal ordering: at the same collection stage, earlier-restricted clades displayed more pronounced differentiation-associated programs, whereas later-restricted clades retained stronger proliferative signatures (**Fig. 4J**). These results support the interpretation that the inferred restriction-time differences reflect asynchronous progression through neural crest differentiation.

Together, these analyses support prolonged neural crest multipotency as most cells remain bipotent or multipotent from neural tube emigration through late migration, whereas only a minority generate a single cell type (*27, 57, 59*). Clade-specific differences in restriction timing within the same derivative class identify clonal bias as previously underappreciated contributor to neural crest differentiation in vivo.

### Lineage-inferred timing resolves regional maturation in skeletal derivatives

In addition to early diversification, we next set out to resolve later differentiation programs and regional differences in skeletal derivatives, comparing cranial neural crest-derived and trunk mesoderm-derived lineages. Focusing on E14.5 skeletal mesenchymal lineages, we identified five coherent lineage-specific gene modules that stratified major skeletal cell types and differentiation programs (**fig. S19, A to D**). Pathway enrichment of module genes highlighted expected biological themes, including extracellular matrix organization and morphogenesis, osteogenic/bone formation programs, and cell-cycle–associated activity. In parallel, we infer developmental-time progression from lineage tree for each skeletal derivative and quantified maturation kinetics, revealing distinct differentiation dynamics across lineages (**fig. S19E**).

Comparisons between cranial and trunk origins revealed region-dependent differences in both module usage and lineage-inferred timing. Osteogenic progenitors displayed a pronounced temporal shift, with trunk-derived cells biased toward later inferred developmental time and enriched for morphogenesis-associated programs (module 2) together with elevated proliferation signatures (modules 3 and 5), consistent with a less mature transcriptional state (**fig. S19, F and H**). By contrast, although chondrocytes exhibited regional differences in proliferative gene expression, they showed uniformly high activation of the lineage-associated cartilage program (module 1) and broadly similar inferred timing across regions (**fig. S19, G and I**). These findings demonstrate that lineage information can resolve anatomically patterned maturation dynamics at later developmental stages.

### Embryo models reproduce cell identities but differ in lineage organization

How developmental history relates to present cell identity and subsequent fate is a longstanding question in developmental biology. In the mouse gut endoderm, for example, cells of embryonic and extraembryonic origin converge on similar transcriptional identities despite their distinct lineage histories (*23, 24*). This distinction is particularly relevant to stem cell-derived embryo models, which reproduce many embryonic transcriptional identities (*19, 20*), yet such resemblance alone does not establish whether those identities arise through comparable developmental routes. We therefore asked whether shared cell identities in embryos and embryo models arise within comparable lineage relationships.

We used EiTiX embryoids, in which mESCs, iCdx2 mESCs, and iGata4 mESCs assemble embryonic, extraembryonic ectoderm-like, and visceral endoderm-like compartments, respectively (*19, 65*). Incorporating DNA Typewriter-engineered mESCs into the embryonic compartment and maintaining doxycycline throughout culture enabled paired transcriptomic and lineage analysis at day 8 (EiTiX8; **Fig. 5A**). EiTiX8 generated recognizable embryonic and extraembryonic-like populations. Harmonized annotation with E8.5 embryos identified shared cell identities supported by marker expression and cross-system transcriptomic similarity, despite differences in cell-type proportions (**Fig. 5B** and **fig. S20, A to C**). We reconstructed the EiTiX8 lineage tree using the same Tape-processing framework applied to embryos (**Fig. 5C**). Hotspot further identified coherent lineage-associated gene modules involving extracellular matrix organization, adhesion, motility, and epithelial differentiation (**fig. S20, D and E**).

**Fig. 5.**
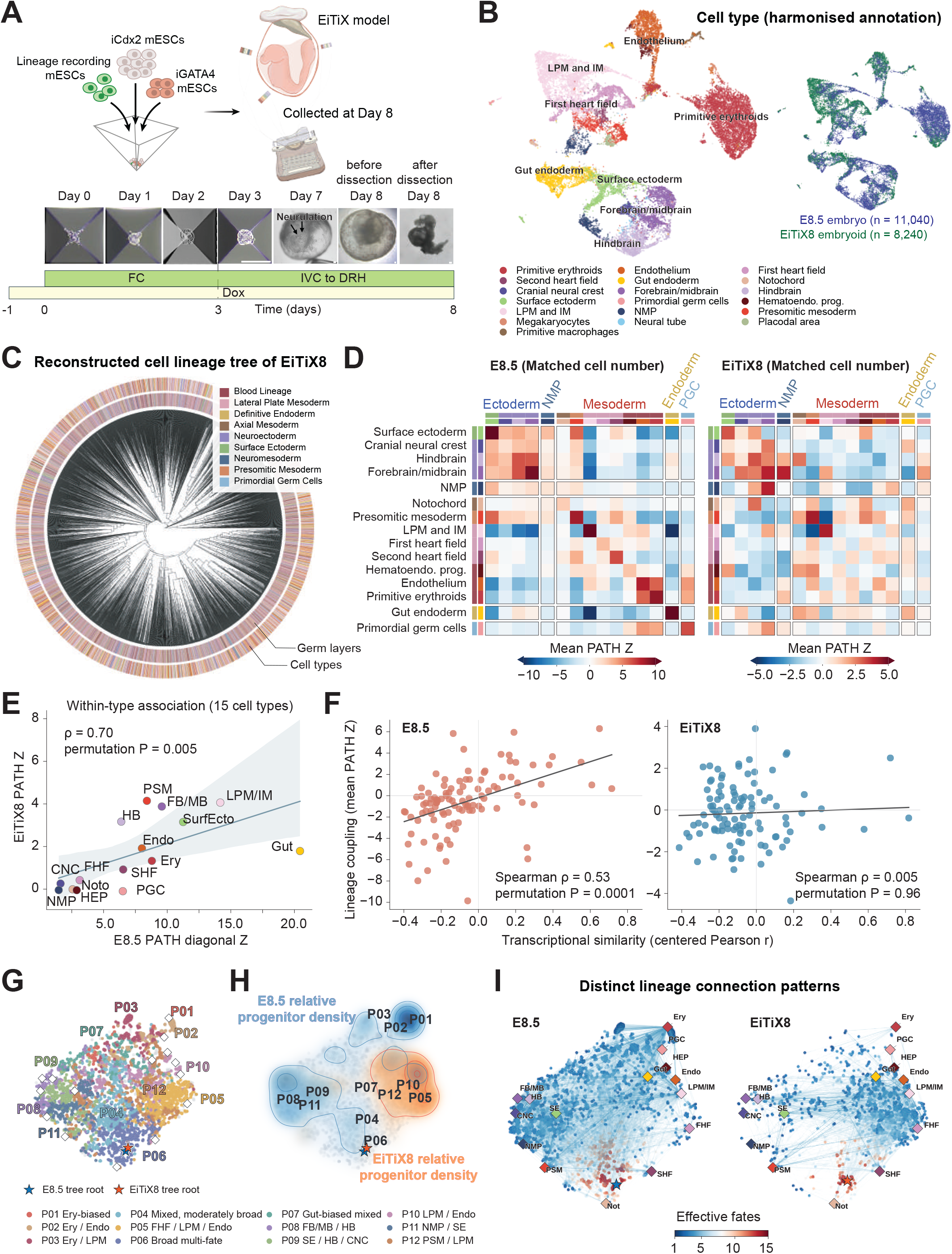
Shared cell identities arise within distinct lineage histories in embryos and EiTiX embryoids. **(A)** EiTiX-embryoids assembly, recording, and representative culture images. **(B)** Joint UMAP of E8.5 mouse embryo cells and day-8 EiTiX embryoid cell after exclusion of extraembryonic populations and reciprocal-PCA integration. The same embedding is colored by harmonized cell-type annotation (Left) or source (Right). **(C)** Reconstructed lineage tree of 3,606 day-8 EiTiX embryoid (EiTiX8) cells, excluding extraembryonic tissues. Inner and outer rings indicate nine broad lineage groups (colored as in the legend) and 18 cell types (colored as in **(B)**), respectively. **(D)** Mean PATH Z scores for 15 shared cell types over 200 rounds of cell-type-matched subsampling, each containing 2,894 cells per system with matched type counts. Color limits differ between systems. **(E)** Cross-system agreement of within-type lineage associations. Each point represents one shared cell type, comparing its mean diagonal PATH Z score in **(D)** between E8.5 and EiTiX8. The line indicates a linear fit; shading shows the 95% confidence band from cell-type bootstrap resampling. **(F)** Transcriptional similarity versus lineage coupling within each system. Each point represents one of 105 distinct pairs among the 15 shared cell types. Transcriptional similarity is the Pearson correlation of gene-centered pseudobulk profiles; lineage coupling is the symmetrized mean PATH Z score from **(D)**. Lines indicate linear fits. **(G)** Joint t-SNE of FateVec profiles from both E8.5 and EiTiX8 lineage trees, defined over 15 shared cell types. Colors identify 12 groups of similar ancestral fate profiles (P01–P12); labels summarize their dominant fate associations; Internal nodes outside the grouped subset are gray. **(H)** Kernel-density estimates of internal nodes in the shared embedding for E8.5 (blue) and EiTiX (orange). Each internal node contributes equal weight; leaves are excluded. Both systems use the same kernel bandwidth and density normalization, without matching node counts. **(I)** Terminal identity connections from each lineage tree projected onto the shared embedding. Node colors indicate effective fate breadth, defined as the exponential Shannon entropy of the normalized FateVec profile across the 15 shared types; higher values indicate broader fate associations. Diamonds mark observed terminal cell-type profiles, and stars mark tree roots.

This correspondence in cell identity did not extend uniformly to lineage associations. After matching cell numbers between systems for each of the 15 shared cell types, within-type PATH autocorrelation values were positively correlated between systems (Spearman ρ = 0.70, permutation P = 0.005; **Fig. 5, D and E**), indicating partial conservation of within-type lineage associations. By contrast, between-type lineage associations across the 105 distinct cell-type pairs showed much weaker concordance (ρ = 0.09, permutation P = 0.34, **fig. S21A**). Repeated matched-size sampling and permutation controls supported this distinction (**fig. S21, B and C**).

Remarkably, some lineage associations reversed direction. For example, gut endoderm was negatively associated with presomitic mesoderm in E8.5 embryos but positively associated in EiTiX8 (matched-cell-number PATH Z scores of −4.69 and 1.38, respectively). Its associations with hindbrain and forebrain/midbrain populations shifted in the opposite direction, from positive to negative (**fig. S21, D to F**). PGCs also showed a different pattern of lineage associations. In natural embryos, PGCs showed strong lineage associations within their own population and with mesoderm population, including lateral plate mesoderm and blood-related populations (**Fig. 5D** for E8.5; **fig. S12C** for E7.5). This pattern is consistent with the proximal epiblast origin of PGC precursors and their early association with presumptive extraembryonic mesoderm (*66*). In EiTiX8, however, PGCs showed lineage associations with NMPs and ectodermal populations (**Fig. 5D**).

Thus, EiTiX partially preserved within-type lineage associations, whereas the between-type lineage associations differed substantially from those in embryos.

### Shared cell identities are embedded in distinct ancestral contexts

In the absence of direct lineage records, transcriptional similarity can help infer developmental relationships, but the extent to which it reflects shared ancestry may differ between systems. We next examined this relationship in E8.5 embryos, where expression similarity across cell-type pairs correlated moderately with lineage coupling (Spearman ρ = 0.53, permutation P = 0.0001). By contrast, this association was nearly absent in EiTiX8 (ρ = 0.005, permutation P = 0.96; **Fig. 5F**). Thus, transcriptional similarity showed little correspondence with recorded lineage coupling in EiTiX8.

To examine the progenitor contexts underlying these differences, we applied FateVec to characterize inferred ancestors by the cell-type composition of their descendants. Joint embedding of internal-node fate bias vectors across the 15 shared endpoint identities resolved 12 ancestry-profile groups, spanning broad multi-fate and lineage-biased compositions (**Fig. 5G** and **fig. S21G**). The inferred ancestral fate profiles differed markedly between the two systems (**Fig. 5H**). For example, P05, an ancestry-profile group associated with first heart field, lateral plate mesoderm, and endothelium, accounted for 1.2% of analyzed E8.5 internal nodes but 42.1% of EiTiX8 nodes (**fig. S21G**). The ancestry profiles associated with individual endpoint identities also differed (**Fig. 5I and fig. S21H**). Forebrain/midbrain and hindbrain populations were prominently linked to the neuroectoderm-associated P08 group in embryos, whereas their EiTiX8 counterparts were more closely linked to mixed profiles such as P05 and P10 (**fig. S21H**).

Together, these analyses indicate that shared cell identities in EiTiX and embryos are embedded in distinct reconstructed lineage histories. Transcriptomic resemblance therefore does not establish equivalence in the ancestral relationships that give rise to those identities.

### Lineage recording supports embryonic lineage contributions to extraembryonic-like states

We next asked whether the distinction between cell identity and lineage history extended across embryonic and extraembryonic compartments. In EiTiX8, DNA Tape marks the engineered mESCs representing the epiblast compartment, whereas extraembryonic visceral endoderm (ExVE)-like cells are expected to arise from the iGata4-induced population (*19*). Nevertheless, 6.3% of annotated ExVE-like cells were Tape-positive (136 cells; **Fig. 6A**). This observation was not restricted to EiTiX: applying the same quality control and doublet filtering to tetraploid-complementation embryos identified Tape-positive cells in E8.5 ExVE (7.1%, 150 cells) and E7.5 extraembryonic ectoderm and visceral endoderm (16.4%, 383 cells, and 16.1%, 558 cells, respectively; **Fig. 6A**).

**Fig. 6.**
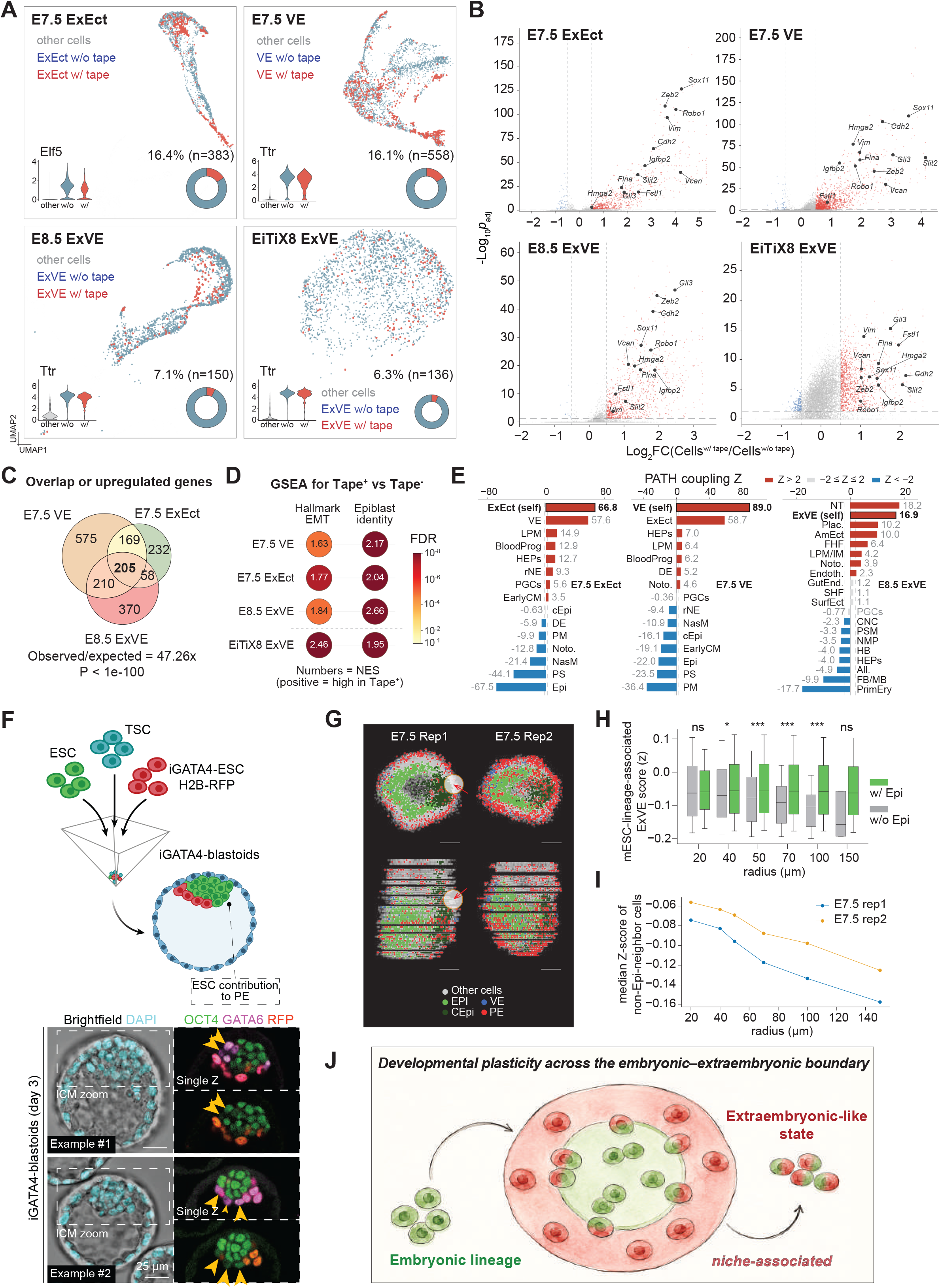
Evidence for lineage plasticity across the embryonic–extraembryonic boundary. **(A)** Transcriptomic locations, marker expression, and Tape-positive fractions in E7.5 extraembryonic ectoderm (ExEct), E7.5 visceral endoderm (VE), E8.5 extraembryonic visceral endoderm (ExVE), and EiTiX ExVE-like cells. Tape positivity requires recovery at least eight edited Tapes per cell in embryos and at least one edited Tapes per cell in EiTiX. **(B)** Differential expression between Tape-positive and Tape-negative cells. Highlighted genes have |log₂ fold change| >0.5 and Bonferroni-adjusted P <0.05. **(C)** Upregulated-gene overlap among the three embryo compartments: 205 shared genes, 47.26-fold above the independent-set expectation; P <10⁻¹⁰⁰, exact three-set overlap test. **(D)** GSEA of epithelial–mesenchymal transition and an independent E5.25 epiblast signature in Tape-positive versus Tape-negative extraembryonic cells. Numbers indicate normalized enrichment scores (NES), with positive NES indicating enrichment in Tape-positive cells; colors indicate false discovery rates (FDR). **(E)** Lineage associations of E7.5 ExEct, E7.5 VE, and E8.5 ExVE across annotated cell types. Bars show PATH Z scores from each target population’s matrix column, including self-association (bold). Positive and negative values indicate stronger and weaker lineage association than expected under the PATH null, respectively. Red, gray, and blue bars indicate Z > 2, −2 ≤ Z ≤ 2, and Z < −2, respectively. LPM: Lateral plate mesoderm; HEPs: Hematoendothelial progenitors; rNE: Rostral neuroectoderm; EarlyCM: Early cardiogenic mesoderm; cEpi: Caudal epiblast; DE: definitive endoderm; PM: Paraxial mesoderm; Noto: Notochord; NasM: Nascent mesoderm; PS: primitive streak **(F)** iGata4-blastoid assay using unlabeled mESCs and mTSCs together with H2B-RFP-labeled iGata4-mESCs (Upper), and representative day-3 images (Lower). Single z-plane images show OCT4, GATA6, and RFP; yellow arrowheads mark GATA6-positive, H2B-RFP-negative cells. Scale bar, 25 μm. **(G)** Cell-state maps from two E7.5 spatial replicates based on the public SEU-3D dataset (GSE278603), shown in top and side views. EPI, epiblast; CEpi, caudal epiblast; VE, visceral endoderm; PE, parietal endoderm. **(H)** mESC-lineage-associated ExVE z scores in pooled VE/PE cells with or without EPI/CEpi neighbors within the indicated radii. Green and gray boxes indicate cells with and without epiblast neighbors, respectively. Boxes, median and interquartile range; whiskers, 10th–90th percentiles. Two-sided Mann–Whitney tests, unadjusted; ns, P >0.05; one, two, or three asterisks, P ≤0.05, ≤0.01, or ≤0.001. **(I)** Median mESC-lineage-associated ExVE z scores of cells lacking EPI/CEpi neighbors, by radius and E7.5 spatial replicate. **(J)** Proposed relationship between local niche and ExVE-like state acquisition.

We next assessed the extraembryonic identity and potential mESC origin of these Tape-positive populations. Tape-positive cells mapped to the same broad cell-type clusters as their Tape-negative (defined as below-threshold) counterparts and expressed canonical extraembryonic markers at comparable levels (**Fig. 6A** and **fig. S22A**). Separate reclustering of the ExVE, VE and ExEct populations nevertheless revealed preferential clustering among Tape-positive cells, suggesting transcriptional differences within these shared identities. Donor-associated SNPs provided additional support for an mESC contribution in embryos (**fig. S22B**), and transcriptional differences between Tape-positive and Tape-negative cells persisted under doublet filtering (scores < 0.1; **fig. S22C**).

To determine whether Tape-positive extraembryonic populations shared transcriptional features across developmental stages and systems, we compared Tape-positive and Tape-negative cells of the same annotation. We identified 205 commonly upregulated genes across the three embryo comparisons, a 47.26-fold enrichment over chance (P < 10^-100^; **Fig. 6, B and C**). Overlap extending to EiTiX8 supported a recurring expression pattern across all four comparisons (**fig. S22, D and E**). Epithelial-to-mesenchymal transition (EMT) was among the most prominently enriched programs (**Fig. 6D** and **fig. S22F**). We further reasoned that, if these cells arose from mESC-derived embryonic lineages, they might retain epiblast-associated transcriptional features despite acquiring extraembryonic identity. We therefore defined an epiblast-associated signature comprising 500 genes highly expressed in E5.25 epiblast cells (*42*). This signature was enriched in Tape-positive cells relative to their Tape-negative counterparts (**Fig. 6D** and **fig. S22G**), consistent with retention of embryonic-lineage features alongside canonical extraembryonic identity. Although uptake of donor-derived material remained an alternative explanation for the lineage and transcriptional signals (*67*), we found no corresponding enrichment of uptake/phagocytosis signatures (**fig. S22H**).

Together, these findings support ESC contribution to extraembryonic-like states. Within reconstructed lineage trees, Tape-positive extraembryonic populations showed within-type clustering and associations with selected embryonic cell types (**Fig. 6E**). This organization suggests a contribution from particular clones and is compatible with a model in which their descendants locate into extraembryonic compartments and expand.

### Evidence for embryonic–extraembryonic plasticity across developmental contexts

Focusing on the endodermal populations, we next asked whether related states could be identified in an independent embryo model (*68–70*). We examined blastoids assembled from unlabeled mESCs and mouse trophoblast stem cells (mTSCs) together with RFP-labeled iGata4 mESCs, distinguishing the directly induced population from the other input cells (*71*). Previous work established that GATA6-positive primitive endoderm-like cells do not form without iGata4 induction in this system (*71*). Alongside the expected GATA6-positive, RFP-positive cells, however, we detected GATA6-positive, RFP-negative cells that retained higher OCT4 expression than their RFP-positive counterparts (**Fig. 6F**). Thus, endodermal marker expression extended beyond the directly induced population and coexisted with an epiblast-associated marker.

We then asked whether a related transcriptional state was present in natural embryos and whether its distribution was associated with proximity to the epiblast. Using shared upregulated genes from the Tape-positive versus Tape-negative comparisons, we defined a transcriptional signature (**table S5**) and applied it to three-dimensional spatial transcriptomic data (*72*). This signature assessed similarity to the Tape-associated expression state rather than assigning lineage origin. After annotating epiblast and VE populations (**Fig. 6G** and **fig. S23A**), we found that VE cells near the epiblast had higher signature scores across multiple neighborhood radii, with consistent associations in both replicates (**Fig. 6, H and I**, and **fig. S23, B and C**). Scores correlated positively with the local epiblast fraction (r = 0.401) and showed a weaker negative association with distance to the nearest epiblast cell (ρ = −0.09, P = 0.001; **fig. S23, D and E**). These results placed a related ExVE expression state near the epiblast in natural embryos.

Previous work established that visceral endoderm of extraembryonic origin contributes to the embryonic gut, demonstrating that extraembryonic ancestry does not restrict cells to extraembryonic tissues (*23, 24*). Our lineage-recording results support plasticity in the complementary direction, providing evidence for an embryonic lineage contribution to extraembryonic-like identities, including ExVE. The blastoid and spatial analyses of unperturbed embryo further identify related endodermal states beyond the tetraploid systems. Together, these findings highlight plasticity across the embryonic–extraembryonic boundary and suggest that extraembryonic cell identity need not imply exclusively extraembryonic ancestry (**Fig. 6J**).

## Discussion

Here, we integrated DNA Typewriter-based lineage recording with tetraploid complementation to generate embryo-wide single-cell transcriptomic profiles with high-resolution genealogical relationships across multiple stages of mouse embryonic development. To analyze this rich dataset with high resolution in lineage relations, we developed FateVec, an analytical framework to infer the dynamics of cell fate decisions and transcriptional regulations from single-cell lineage tree data. This approach allowed us to quantify changes in developmental dynamics as biological phenotype, complementing conventional analyses based on cell-type composition alone. Extending this analysis to embryo models and extraembryonic-like populations further reveals that similar cell identities can be associated with distinct lineage histories.

Using the FateVec framework, we find that the loss of developmental potential is much more progressive and asynchronous than was expected. Multiple lineages display detectable fate bias before overt morphological diversification associated with gastrulation, consistent with emerging evidence that molecular symmetry breaking and lineage priming can precede classical anatomical landmarks (*8, 42*). In this view, the pluripotency exit is not a single transition coordinated across the embryo but an asynchronous process in which distinct lineages undergo commitment at different rates and in distinct time frames. Thus, our results suggest that the blueprint for cell fate bias is laid down significantly earlier than previously appreciated, foreshadowing differentiation trajectories long before the embryo’s morphological landmarks reveal them. While this manuscript was in preparation, an independent preprint also identified early lineage biases and reported their high reproducibility across embryos (*73*). Consistent with these findings, our analyses of independently profiled E7.5 embryos and distinct clades in the pooled E8.5 dataset recovered concordant fate-bias orderings, providing complementary support for reproducibility at the embryo and clade levels.

Lineage-specific transcriptional modules provide a mechanistic link between ancestry and commitment dynamics. Using reconstructed cell lineage trees with high resolution, we identify gene modules whose activities are coherent on the phylogeny and reveal wave-like deployment of programs throughout development. Sequential module activation, together with stage-calibrated restriction timings, supports a multi-phasic architecture in which distinct regulatory programs are engaged in their combinatorial orders, linking morphological staging with underlying gene regulatory kinetics. This framework enhances our understanding of the temporal logic of cell fate decisions. More broadly, this framework provides a quantitative platform for comparing developmental timing across lineages and, in the future, across perturbations*(74)*.

Our focused analysis of neural crest derivatives illustrates the power of lineage-resolved timing to address long-standing debates regarding the multipotency of neural crest progenitors (*26, 27, 53*). These results support multipotency among inferred neural crest progenitors, while also exhibiting substantial interclonal heterogeneity in commitment kinetics. Within nominally identical derivative classes, independent founder epiblasts differed systematically in restriction timing. These "early-restricted" versus "late-restricted" clades correlate with distinct transcriptional states: early clades show stronger differentiation-associated programs, while late clades show stronger proliferation-associated programs. Thus, developmental tempo emerges as a key dimension of heterogeneity, suggesting that the embryo achieves robust organogenesis not through rigid synchronization, but through a diverse repertoire of clonal trajectories that converge on the same functional identity at different speeds.

Comparisons with EiTiX broaden this analysis from variation in developmental tempo to variation in lineage organization. Recognizable cell identities in EiTiX were embedded in lineage relationships that differed from those in embryos, suggesting that transcriptional convergence can occur through different patterns of fate diversification. Ancestral relationships and fate-bias dynamics therefore provide complementary measures of model fidelity, capturing how cell types arise in relation to one another. Together with the clonal heterogeneity observed in embryos, these findings raise a broader question about developmental robustness: which aspects of developmental history can vary while preserving developmental outcomes, and which are required for subsequent developmental competence? Resolving this question would help identify the features of lineage organization that matter for responses to inductive signals and coordinated tissue maturation.

The extraembryonic findings raise a complementary question about how lineage origin constrains the identities cells can acquire. The established contribution of visceral endoderm to embryonic gut illustrates that early lineage segregation does not impose a simple correspondence between ancestry and later tissue identity (*23, 24*). Our observations suggest that cells of embryonic origin may acquire extraembryonic-like states in experimental systems, raising the possibility of flexibility in the complementary direction. Related expression states in blastoids and unmanipulated embryos motivate examining whether this flexibility has physiological relevance. These findings direct attention toward how embryonic and extraembryonic identities become stabilized, and under what conditions cells retain access to programs associated with other lineages (*75*). Establishing the cellular origins of these populations will be essential for defining the extent and developmental significance of this plasticity.

Several limitations remain, including incomplete cellular sampling and restricted developmental coverage. Cell-type-dependent retention during filtering cells with sufficient Tape recoveries for lineage reconstruction may distort population representation, particularly at later stages, so cell-type proportions in the trees may differ from those in the embryo. Targeted profiling of rare populations, such as PGCs, together with improved recording efficiency and capacity, should enable more comprehensive analysis across tissues and developmental stages. For extraembryonic populations, complementary genomic lineage tracing could help establish cellular ancestry by distinguishing donor-cell contributions from possible transfer of donor material.

A central next step is to establish how developmental history interacts with molecular state and tissue context to shape fate acquisition. Integration with additional modalities, including chromatin state, transcription-factor activity, and spatial transcriptomics, should further enrich our understanding of the role of cell history in cell fate. Combined with genetic or chemical perturbations, lineage-based frameworks such as FateVec should help define how specific signalling pathways alter commitment kinetics *in vivo*. Our current analyses focus on recorder-bearing postimplantation lineages in tetraploid-complementation embryos. Establishing mouse lines carrying the recorder could extend tracing to preimplantation development and broaden coverage of both embryonic and extraembryonic lineages, supporting a more comprehensive view of embryogenesis. This, in turn, may provide a deeper understanding of the logic of robust developmental programs and inform efforts to engineer functional tissues from stem cells.

## Acknowledgments

We thank the members of the Zernicka-Goetz, Choi, Bronner, Kim, Thomson, and Wei labs for helpful discussions, and Itai Yanai, Sarkis Mazmanian, Pietro Perona, and Jake Cornwall-Scoones for feedback and advice. Funding: This work was supported by the National Institutes of Health (R00HG012973 to J.C. and Director’s Pioneer Award DP1HD104575A to M.Z.-G.). J.C. also received support from the Searle Scholars Award, the Damon Runyon Cancer Research Foundation (DFS-64-24), and the National Cancer Institute Cancer Center Support Grant (P30CA008748). M.Z.-G. also received support from the Allen Discovery Center for Cell Lineage Tracing (UWSC13405).

## Data and materials availability

The raw and processed sequencing data have been deposited into the NCBI Gene Expression Omnibus (GEO) database under accession number GSE324482. The custom analysis code used here is available at: https://github.com/choi-junhong/mouse-lineage-recording. The Mouse IT resource (www.mouse-it.com) provides interactive exploration of the reconstructed lineage trees. An accompanying video tutorial explaining FateVec is available on YouTub (https://www.youtube.com/watch?v=srKANcVEVYo).

## Author contributions

Conceptualization: Y.L., J.C., M.Z.-G;

Experiments: Y.L., with assistance from J.P., L.A., S.J., Y.W., S.N., S.E.P., S.L., H.W.;

Formal analysis: Y.L., J.C., with assistance from C.W.J., J.P., L.A., X.S., Y.Lu, S.J.;

Methodology: Y.L., J.C.;

Investigation: Y.L., J.C.;

Visualization: Y.L., J.C.;

Funding acquisition: J.C., M.Z.-G;

Supervision: J.W., H.H.K., M.T., C.W.J., M.E.B., J.C., M.Z.-G;

Writing: Y.L., J.C., M.Z.-G.

## Materials and Methods

### Tetraploid embryo generation, microinjection, embryo transfer and dissociation

All experiments were approved by the Institutional Review Board and the Institutional Animal Care and Use Committee (IACUC) at the California Institute of Technology. Two-cell stage embryos were collected from 6- to 12-week-old B6D2F1 females after superovulation by priming of 5 IU of pregnant mare’s serum gonadotropin (PMSG, Prospec tany technogene) and 5 IU of human chorionic gonadotropin (hCG, Patterson Veterinary) 48 hours later. BDF1 female mice were mated with BDF1 males (Jackson Laboratory). Two-cell stage embryos were collected 44 hours after hCG injection in M2 medium (Embryo Max, Sigma Millipore) and incubated in Advanced KSOM medium (Embryo Max, Sigma Millipore) until the electrofusion.

For electrofusion, embryos were washed three to five times with 300 mM Mannitol-based solution (prepared in-house) and placed in the fusion chamber (BTX, 1.0 mm gap). A single DC pulse (BTX 2001, 150V, 60 µs) was applied to generate the tetraploid embryos, as described previously (*76*). After the electrofusion pulse, embryos were washed three times with M2 medium and three times with KSOM medium, respectively. The electrofusion was assessed 30 minutes later. The fused (tetraploid) embryos were further cultured in KSOM medium at 5% CO2 and 37 °C for two days (until E3.5).

DNA Typewriter–engineered cells were briefly trypsinized and resuspended in cold M2 medium before microinjection. 12 cells were microinjected into the cavity of E3.5 blastocysts. Injected embryos were cultured at 37 °C, 5% CO2 in KSOM medium until uterine embryo transfer two to three hours later. 10 to 20 blastocysts were transferred into CD1 (Charles River) pseudo-pregnant females (2.5 days after copulation with vasectomized males). 100 µl of 10 mg/ml Dox was injected intraperitoneally immediately after embryo transfer. The Dox level was maintained with Dox chow (200 mg/kg) and Dox water (0.4g Dox and 10g sucrose in 200 mL water), both of which were replenished every two days to prevent Dox degradation.

Implanted embryos were collected at E7.5, E8.5, E9.5, E10.5, E11.5, E13.5 and E14.5. E7.5 and E8.5 embryos were processed whole. The two E7.5 embryos were processed separately. Embryos from E9.5 to E13.5 were dissected into head and body, and E14.5 embryos were dissected into head, body and vertebral regions. For each collection from E8.5 onward, we collected embryos from one recipient female mouse and pooled for downstream processing, including four embryos at E8.5, two at E9.5, three at E10.5, two at E11.5, one at E13.5 and three at E14.5. Tissues were minced into small pieces, transferred into 5 ml tubes and dissociated with TrypLE Express Enzyme for 20 min. Cells were pelleted by centrifugation at 200 rcf for 5 min and washed twice with 1× PBS containing 0.2% BSA. The cell numbers were counted manually using 2-Chip Hemocytometer and automatically using Countess II Cell Counter. The viability was evaluated (>90%) and the single-cell suspension was then ready to follow the 10x Genomics experimental protocol for scRNA-seq library preparation.

### Cell lines

We cultured the mESC lines (CD-1 outbred strain) on 0.2% (w/v) gelatin-coated plate and maintained them in N2B27 2iLIF medium in 5% CO_2_ at 37 °C. mTSCs were cultured with irradiated C57bl/6 Mouse Embryonic Fibroblasts (MEFs) in TSF4H medium at 37°C in hypoxic conditions of 5% O_2_ and 5% CO_2_, as previously described (*71*). The DNA Typewriter–engineered mESC line was transfected by mixing PB-Tape, PB-iPEmax-BlastR, and Dox-inducible PEmax-P2A-mCherry expression with Lipofectamine 3000 reagent. Those single clones with high GFP expression were manually picked under microscope to minimize the passage number of the cells. These single clones were grown in 96-well plates first and expanded to 6-well plates. All cell lines were tested every two weeks for mycoplasma contamination. For the recorder induction time course, the clonal DNA Typewriter mESC line was cultured with 0, 0.25, or 1 μg ml^−1^ doxycycline and sampled over 0–12 days. Bulk genomic DNA amplicon sequencing quantified editing frequency at each of the six sequential Tape sites and the fraction of tapes reaching each cumulative editing depth.

### Generation of EiTiX embryoids

EiTiX embryoids were generated using a modified version of a previously described aggregation protocol (*19, 65*). The day of cell aggregation was defined as Day 0. DNA Typewriter–engineered mESCs were treated with doxycycline beginning 24 h before aggregation at a final concentration of 100 ng/mL. Doxycycline was maintained at 100 ng/mL^−1^ in subsequent culture media and replenished daily until the experimental endpoint. The mESC line used to generate EiTiX embryoids carried 8 Tapes, whereas the line used for tetraploid complementation carried 50 Tapes. Accordingly, a minimum of 4 Tapes per cell was required for inclusion in the EiTiX lineage analysis.

For EiTiX embryoid assembly, iCdx2 mESCs and iGata4 mESCs were induced with doxycycline for 6 h before aggregation. AggreWell400 plates were prepared by adding 500 μL anti-adherence rinsing solution to each well, followed by centrifugation at 2,000g for 5 min to remove trapped bubbles. After a 30 min incubation, the rinsing solution was removed, and each well was washed twice with PBS. The dissociated iCdx2 ESCs, iGata4 ESCs and doxycycline-pretreated DNA Typewriter–engineered mESCs were manually counted using a 2-Chip Hemocytometer and combined at 38,400 iCdx2 ESCs, 19,200 iGata4 ESCs and 19,200 DNA Typewriter–engineered mESCs per AggreWell. The pooled cells were pelleted at 1,000 rpm for 4 min and resuspended in 1 mL FC medium supplemented with 1 μg mL^−1^ heparin, 25 ng mL^−1^ FGF4 and 7.5 nM Y27632. The cell suspension was added dropwise into AggreWells preloaded with 500 μL FC medium containing heparin and FGF4, and the plate was centrifuged at 100g for 3 min to distribute cells into the microwells.

On day 1, 1 mL medium was replaced with fresh FC medium, and the medium exchange was repeated once. On day 2, 1 mL medium was replaced with fresh FC medium. On day 3, 1.3 mL medium was removed from each well and 1.5 mL peri-implantation medium containing 20% iFBS was added. Peri-implantation medium consisted of Advanced DMEM/F12 supplemented with, 2 mM GlutaMAX, 1% penicillin-streptomycin, 1× insulin-transferrin-selenium-ethanolamine, 8 nM β-estradiol, 200 ng mL−1 progesterone and 25 mM N-acetyl-L-cysteine. On day 4, morphologically appropriate structures were selected under a dissection microscope and transferred, up to 20 structures per well, into non-adherent 6-well plates containing 5 mL peri-implantation medium with 30% FBS. On day 5, structures were transferred into peri-implantation medium containing 30% knockout serum replacement.

For post-implantation culture, selected Day 5 EiTiX embryoids were transferred individually into wells of a 48-well suspension culture plate containing 250 μL post-implantation culture medium. Post-implantation medium was prepared from DMEM supplemented with 1× GlutaMAX, 100 units/ml penicillin-streptomycin and 11 mM HEPES, together with 50% heat-inactivated rat serum and 25% human cord serum. The medium was filter-sterilized and equilibrated at 37°C before use. On day 6, 100 μL medium was removed and 250 μL fresh post-implantation medium was added to each well. On day 7, EiTiX embryoids were transferred to custom-made ex vivo culture system, hereafter referred to as the roller culture system, with up to three embryoids cultured per bottle in 2 mL post-implantation medium supplemented with 3.0 mg mL−1 D-glucose with 20% O_2_ and 5% CO_2_.

### Generation of blastocyst-like structures

The blastocyst-like structures, iG4-blastoids, were generated using a previously described aggregation protocol (*71*). On the day of aggregation, WT ESCs and tetO-Gata4 ESCs were dissociated into single-cell suspensions using 0.05% trypsin-EDTA at 37°C for 3 min. Cells were collected by centrifugation at 1,000 rpm for 5 min, resuspended in FCLY medium and counted using a 2-Chip Hemocytometer. For each AggreWell, 6,000 WT ESCs and 4,800 tetO-Gata4 ESCs were combined in FCLY medium and added dropwise to the microwells to a final volume of 1 mL per well. Unused wells were filled with PBS to maintain humidity and reduce evaporation. Plates were centrifuged at 100g for 3 min and then incubated at 37°C under normoxic conditions with 5% CO_2_ and 20% O_2_. FCLY medium consisted of feeder cell medium supplemented with 10 ng mL^−1^mouse recombinant LIF and 2 nM Y27632. Feeder cell medium was composed of DMEM supplemented with 15% inactivated FBS, 1% GlutaMAX, 0.1 mM 2-mercaptoethanol, 1% non-essential amino acids, 1% sodium pyruvate and 1% penicillin-streptomycin.

After ESCs had compacted into single aggregates, typically 6–8 h after seeding, we seeded mTSCs. mTSC colonies were dissociated into single-cells with 0.05% trypsin-EDTA for 10-12 min, pelleted by centrifugation and resuspended in TSF4H medium. To deplete MEFs, the cell suspension was plated onto 0.1% gelatin-coated dishes and incubated at 37°C for 1 hour. Non-adherent mTSCs were then collected, counted using a 2-Chip Hemocytometer and added to the preformed ESC aggregates at 14,400 mTSCs per well. Before mTSC addition, FCLY medium was removed and replaced with TX medium supplemented with LIF. mTSCs were added dropwise, and the AggreWell plate was returned to 37°C under hypoxic conditions with 5% O_2_.

TX medium supplemented with LIF consisted of DMEM/F12, 64 mg/L L-ascorbic acid-2-phosphate magnesium, 14 μg/L sodium selenite, 19.4 mg/L insulin, 543 mg/L NaHCO3, 10.7 mg/L holo-transferrin, 25 μg/L human recombinant FGF4, 2 ng/mL human recombinant TGF-β1, 1 mg/L heparin, 1% GlutaMAX, 1% penicillin-streptomycin, 20 μM Y27632, 3 μM CHIR99021, 1 mM 8-bromo-cAMP, 30 ng/mL IL-11 and 10 ng/mL mouse recombinant LIF. Twenty-four hours after mTSC seeding, tetO-Gata4 ESCs were induced by supplementing the culture medium with 1 μg/mL doxycycline and 100 μM 8-bromo-cAMP.

### Immunofluorescence staining and confocal imaging

Stem cell-derived embryo-like structures were fixed in 4% paraformaldehyde for 20 min at room temperature. Samples were then washed twice in PBST, consisting of PBS supplemented with 0.05% Tween-20, and permeabilized for 30 min at room temperature in PBS containing 0.3% Triton X-100 and 0.1% glycine. After permeabilization, samples were incubated overnight at 4°C with primary antibodies diluted in blocking buffer containing PBS, 10% fetal bovine serum and 1% Tween-20. The next day, samples were washed twice with PBT and incubated overnight at 4°C with species-appropriate secondary antibodies diluted 1:500 in blocking buffer. On the third day, samples were washed twice in PBST and transferred into PBST droplets in oil-filled optical plates for confocal imaging. We used the primary antibodies of OCT4 (Cell Signaling Technology, 75463S) and GATA6 (R&D Systems, AF1700). Images were acquired on an inverted Leica SP8 confocal microscope using either a Leica Fluotar VISIR 0.95 NA ×25 NA objective or a 1.2 NA x40 objective. DAPI was excited with a 405 nm diode laser, Alexa Fluor 488 or GFP with a 488 nm argon laser, Alexa Fluor 543/555 with a 543 nm HeNe laser, and Alexa Fluor 633/647 with a 633 nm HeNe laser. Z-stacks were collected with a 2.5 μm interval between optical sections.

### 10x Genomics scRNA-seq library preparation and sequencing

Libraries were prepared according to the Chromium Next GEM Single-Cell 3′ Reagent Kit v3.1 with Feature Barcoding Technology for CRISPR Screening. Briefly, we collected both cDNA from poly-adenylated mRNA in 2.3A and cDNA from Feature Barcode compatible sgRNA in 2.3B. The cDNA product from 2.3A was used for further 3’ Gene expression library construction, while both 2.3A and 2.3B were used for the enrichment of DNA Tape sequences through PCR, as previously described (*16*).

### 10x Genomics scRNA-seq processing and analysis

The scRNA-seq fastq files from mouse embryos spanning developmental stages E7.5 to E14.5 were pre-processed using Cell Ranger count pipeline (version 8.0.1) (*77*). This was followed by the processing and filtering by the Seurat package (version 5.3.0) in R (*78*). Cells were retained if they satisfied all the following thresholds: number of unique molecular identifiers (nUMI) ≥ 500, number of genes ≥ 250, and mitochondrial gene ratio < 0.10. The gene-level filtering was then applied by retaining genes detected in at least 10 cells. The filtered count matrix was then used to reconstruct Seurat objects which were saved and used as input for downstream analysis. Quality-control metrics for each library are summarized in **table S1** and were consistent with high-quality single-cell datasets. Technical duplicates across the three E8.5 10x reactions were identified using shared 16-nt cell barcodes and concordant DNA Tape insertion patterns. Within each group, we retained the Tape profile associated with the highest transcriptomic UMI count and merged transcriptomic records before annotation.

Gene expression counts were log-normalized, scaled, and subjected to principal component analysis. A k-nearest-neighbor graph was constructed using the first 15 principal components, followed by graph-based clustering at a resolution of 2. Uniform manifold approximation and projection (*79*) was used to visualize transcriptional relationships among cells. Cluster-enriched marker genes were identified by comparing each cluster with all remaining cells and retaining positively enriched genes. Cell identities were assigned based on developmental stage, canonical marker genes, and published references (*30, 41*), as summarized in **table S6**. The resulting annotations were used for downstream lineage visualization and fate-restriction analyses.

### Processing of DNA Tape sequencing reads

The sequenced DNA Tape reads were initially processed along with the cDNA reads using 10X Genomics’ Cell Ranger, which corrects cellBC in each read based on its own allowed list. From the resulting sequencing reads, we follow the 8 steps outlined in **fig. S3E** for data pre-processing. First, we tabulated the number of reads and associated RT-UMIs for each unique DNA Tape read in each cell, and retained reads associated with more than 1 distinct RT-UMI. Second, we extracted 12-bp TapeBC and retained sequences that match to the list of 50 allowed TapeBC (**table S7**), previously determined through bulk-gDNA amplicon sequencing of DNA Tape, with fewer than two nucleotide substitutions. Third, reads with cellBC matching the list that passed transcriptomic quality control were retained. Fourth, the DNA Tape reads were searched from their first to sixth insertion sites for sequential insertions in the form of NNNGGA’s. Fifth, in each CellBC-TapeBC combination, the most frequent insertion patterns were selected based on the number of associated RT-UMIs. In case there are two insertion patterns with the same RT-UMIs, the number of read counts were used to determine the most frequent insertion pattern. Sixth, data from multiple scRNA-seq reactions used for each embryo collection time point were combined and formatted tracking their sample origin. Seventh, edit distances based on the sequential insertions were calculated for each cell pair, resulting in the cell-by-cell distance matrix. Finally, we retained cells with at least eight recovered, edited DNA Tape molecules for the lineage tree reconstruction.

### Lineage tree reconstruction and phylogenetic correlation analysis for DNA Tape

From the resulting cell-by-cell edit distance matrix for cells with at least eight recovered, edited DNA Tape molecules, we use the UPGMA method for reconstructing the tree in each embryonic stage. Phylogenetic correlation across the annotated cell types were calculated using the PATH package (*31*), where the inverse distance across the entire tree was used. Lineage-specific gene modules are selected using the Hotspot package (*32*), using the recommended parameters for K-nearest-neighbors (30), minimum gene threshold (50), and FDR threshold of 0.05 for tetraploid embyro and 0.25 for EiTiX.

### Subsampling analysis for tree stability

To evaluate the stability of the inferred lineage tree and downstream lineage-informed analyses, we performed resampling of cells for reconstructing lineage trees. In each subsampling iteration, 80% of cells were randomly sampled from the full dataset without replacement, and the lineage tree was reconstructed using the same pipeline as in the original analysis. This procedure was repeated 50 times. For each subsampling tree, node depth was calculated as a proxy for developmental progression, and the fate-bias trajectories and *T_on_* were recomputed using the same procedure as for the full tree. Tree stability was quantified by evaluating the consistency of node depths and the preservation of pairwise rank relationships among cell types across all 50 replicates, providing a measure of the robustness of both tree topology and lineage-informed developmental inferences.

### Permutation-based empirical null analysis

To assess whether FateVec-inferred restriction timing could arise from cell-type abundance or random assignment of cell identities on the tree, terminal cell-type labels were shuffled across tree tips while preserving the number of cells in each cell type. Tree topology was kept fixed, and the FateVec pipeline was rerun for each permuted dataset. This procedure was repeated for 1,000 permutations using the reference E8.5 germ-layer tree. For each permutation, we calculated the Pearson correlation between FateVec-inferred restriction timing and the expected embryonic ordering. Empirical P values were defined as the fraction of permutations with correlation values equal to or greater than the observed value. As a more stringent control, the analysis was repeated using only the eight germ-layer anchor points after excluding the induction-start reference anchor. This permutation analysis tested whether the observed FateVec ordering exceeded that expected from random cell-type placement on the lineage tree.

### Theory of lineage fate vectorization

To quantify fate restriction timing for each annotated cell type, we developed an algorithm, FateVec, to assigns every node of a reconstructed lineage tree a fate vector, a population-weighted measure of how strongly each cell type’s descendants are confined to lineages restricted to that type and follows these vectors along the ancestry of each cell type. The tree was stored as a set of nodes, each representing a cell or progenitor at a specific developmental time point. Input consisted of two files: (i) a cell-type annotation file mapping each terminal cell barcode to its annotated identity, and (ii) a tree node summary file specifying, for each node: a unique node identifier, a Boolean tip flag, the cell barcode (for tip nodes), the parent node identifier (-1 for the root), the cumulative branch-length distance from the root, and the number of tip descendants in the subtree rooted at that node. The tree was reconstructed in memory using a custom Python node class storing, for each node, its parent, children, depth, tip status, and — for tip nodes — the assigned cell-type index.

Each node *p* carries two vectors of length *K* (the number of distinct annotated cell types): a descendant count vector *N*_*p*_ = (*N*_*p*_^1^, …, *N*_*p*_^*k*^), recording the number of descendants of each fate, and a fate vector *V*_*p*_ = (*V*_*p*_^1^, …, *V*_*p*_^*k*^), recording the normalized fate proportions. Tip nodes were initialized with one-hot vectors: *N*_*p*_^*k*^ = *V*_p_^*k*^ = 1 for the assigned cell-type index *k*, and 0 otherwise. Internal nodes were populated by post-order (bottom-up) traversal. After all children of node *p* had been updated, their vectors were aggregated as:

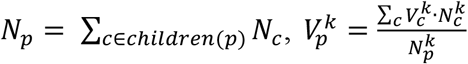

with *V_p_*^*k*^ set to zero wherever *N_p_*^*k*^ = 0. The resulting *V*_*p*_ was then renormalized to sum to unity, yielding a fate vector that reflects the weighted average contribution of each terminal fate to the node’s descendant population. It measures how exclusively the type-*k* descendants of progenitor *p* lie in lineages restricted to type *k*, and it is not the fraction of *p*’s descendants that are of type *k*.

### Temporal mapping of fate restriction timing

Node depth, the cumulative branch length from the root, served as a pseudotime coordinate that increases monotonically from root to tips, as a proxy for developmental progression. Because the tree is ultrametric, depths were normalized so that the root lies at 0 and all tips at 1. All trajectories were evaluated on a common grid of 10,001 evenly spaced points on [0, 1].

For each terminal cell type *k*, we traced the *k*-th component of fate vector ***V*** along the ancestral path of every tip of type *k*. The resulting single-tip trajectory is a step function of pseudotime: on each branch it takes the value of the node at the branch’s tip-ward end, so it begins at the value of the root’s child on that path and reaches 1 at the tip. Averaging over all *N_k_* tips of type *k* to yield the mean fate vector trajectory:

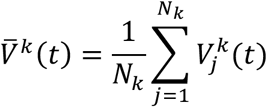

which represents the progressive enrichment of fate *k* along lineages leading to that terminal identity.

We computed the first derivative *dV^k^(t)/dt* using a Savitzky–Golay filter (*80*), which suppresses discretization noise while preserving transition shape. For each fate, *T_on_* was defined as the leading half-maximum crossing on the rising flank preceding the maximum of the derivative profile. *T_on_* therefore summarizes the onset of rapid fate-bias acquisition; peak height and the depth of the global maximum are distinct quantities and were not used as the onset metric. We compared *T_on_* across fates on normalized tree depth and mapped it to embryonic time by linear regression against the reference developmental landmarks (*30, 37, 38*). Bootstrap calibration was performed separately using the mean *T_on_* across resampled trees.

### Carta potency-set analysis

We applied a local implementation of Carta’s unweighted large-k objective *(36)* to the E8.5 non-ExVE tree. Candidate progenitor sets had to contain each node’s observed descendant states and nest within its parent’s assigned set. We minimized the total excess set size across 7,814 eligible nodes, ignoring branch lengths. Dynamic programming assigned nodes to candidate sets; simulated annealing and greedy refinement searched k = 2–8 with 16 restarts and 200,000 proposals per restart. We retained the lowest-cost feasible solution and displayed the six-set solution. This heuristic inferred potency sets without estimating developmental dates or guaranteeing a global optimum.

### Ancestral fate profiles and cross-system comparisons

We embedded the FateVec profiles of inferred ancestral nodes and visualized their effective fate breadth, density, and descendant outputs. Effective fate breadth was defined as the exponential of the Shannon entropy of each node’s normalized FateVec profile, using natural logarithms and taking zero-weight contributions as zero. Higher values indicate a broader distribution of inferred fate associations, rather than experimentally measured differentiation capacity. For comparisons between E8.5 embryos and EiTiX8, node profiles were represented in the same shared-fate space before joint embedding and group-level comparison.

For E8.5, we grouped all 10,578 internal nodes from the 10,579-tip non-ExVE tree by k-means on square-root FateVec profiles, weighting nodes equally. t-SNE embedded distinct profiles, including leaves, with perplexity 100, 2,000 iterations, PCA initialization, automatic learning rate and seed 42; identical profiles shared coordinates. Terminal-output summaries assigned each leaf to its immediate parent’s group.

For joint E8.5–EiTiX8 analysis, we set contributions from unmatched fates and ExVE to zero and recomputed FateVec over the 15 shared states. Joint k-means used internal nodes with at least five matched tips in at least two child subtrees, yielding 12 independently defined groups (P01–P12; 30 initializations; seed 42). Group frequencies used eligible grouped nodes within each system as denominators, without matching node counts. Joint t-SNE included all nodes with matched descendants. Gaussian density estimates weighted internal nodes equally, used bandwidth 5% of the maximum coordinate span and shared one maximum for scaling. Each leaf inherited its nearest P-labeled ancestor’s group; proportions used all leaves of that terminal state, without renormalizing the displayed top three groups.

### Simulation benchmarking and label perturbations

An eight-fate developmental program generated 8,192 terminal cells with unequal fate abundances and prescribed restriction windows. Simulated recordings varied editing schedules, heritable editing-rate heterogeneity, insertion-spectrum skew, Tape recovery, and terminal-cell sampling. We compared fixed recovery of 20 Tapes with 50-Tape recording followed by dropout, and evaluated both absolute and editing-load-normalized pairwise distances. UPGMA trees were reconstructed from recording information without terminal-fate labels, and FateVec was evaluated against the programmed restriction ordering. Separate perturbations introduced convergent fate labels or annotation errors. For the annotation-error analyses, we relabeled either 64 cells from other fates as G or 64 G cells as other fates, corresponding to 10% of the original G population. QFM (*35*) was used to assess progenitor-state recovery under these conditions.

### Reproducibility across embryos and clades

For biological-replicate comparisons, the two E7.5 embryos were analyzed separately. PATH comparisons matched the number of cells per cell type between embryos, and FateVec timing was compared for cell types with more than 200 cells in both embryos. For clade analyses, Ward clustering (*81*) used first-site insertion patterns without embryo labels; labels were revealed afterward for the E7.5 purity assessment. Within-clade trees used subsequent recording sites, and fate rankings were compared for shared sufficiently sampled cell types. E8.5 clade-rank agreement was evaluated against within-clade label permutations. Across clades in E8.5, clades without enough cell type numbers and editing have been excluded. Cell types required at least 20 cells per clade. We compared the median Spearman correlation across 45 clade pairs with 200 permutations that independently shuffled terminal labels within each retained clade and recomputed FateVec and timing.

### Lineage-tree dating and descendant-fate breadth

We dated the E10.5 tree using strict-clock least squares *(61)* and cell-count regularization *(62)*. For this dating analysis, we reconstructed the tree by neighbor joining of Ochiai-normalized insertion-prefix distances, followed by parsimony-based nearest-neighbor interchange and rooting with an unedited outgroup. After removing the outgroup, we assigned branch lengths from sequential insertion gains and collapsed zero-gain branches (14,313 tips). We jointly fitted internal dates and a shared editing rate to minimize squared differences between insertion-scaled branch lengths and rate multiplied by elapsed time, anchoring the root at E4 and tips at E10.5. A log-interpolated growth curve, scaled to three pooled embryos, approximately constrained ordered internal dates by the time needed to reach m + 1 cells for the m-th node. The later counts followed reference 62. We enforced parent–child ordering and collection-time bounds. Dates depended on the clock and growth assumptions.

For neural crest summaries, we retained ancestors with eligible descendants in at least two child subtrees across mesenchyme, neurons, glia and pigment cells. We classified nodes with descendants in one, two or three to four groups as unipotent, bipotent or multipotent, respectively. Fractions used all eligible ancestors of each focal group in E3.5–<E6, E6–<E7, E7–<E8, E8–<E9 and E9–E10.5 intervals. Shared ancestors could contribute to multiple groups. These categories measured observed descendant-fate breadth.

### Sankey diagram construction

To estimate the total cell number at each developmental stage, we applied the DNA-based quantification model as previously described (*41*), which fits a polynomial regression to experimentally measured genomic DNA yields across embryonic time points. For each annotated cell type, the predicted total cell number at a given stage was further scaled by its proportional representation in our scRNA-seq dataset. The resulting per-cell-type abundance estimates (log₂-transformed) were used to scale the link and node widths in a directed tree graph visualized as a Sankey diagram, where nodes represent anatomical regions at successive stages from E9.5 to E14.5 and edges connect regions sharing lineage annotations across adjacent time points.

### Clade-stratified transcriptional analysis

To relate lineage-inferred timing to transcriptional programs, we computed *T_on_* separately for each neural crest derivative represented within each clade. Differential gene expression and gene set enrichment compared earlier- and later-onset clades within the same derivative. Identity-associated expression and proliferation programs, including Hallmark MYC targets, Hallmark E2F targets, and Hallmark G2M checkpoint, were summarized at clade level. We then compared within-derivative timing ranks with transcriptional identity and proliferation ranks.

For E8.5–EiTiX8 comparisons, we harmonized 15 shared terminal-cell annotations and matched cell numbers per type before PATH analysis. Repeated matched-size sampling (200 iterations) and label-permutation controls evaluated the robustness of within-type and between-type comparisons. Transcriptomic similarity between each pair of shared types was compared with the corresponding phylogenetic association within each system; cross-system concordance was evaluated separately for diagonal and off-diagonal PATH entries.

### Extraembryonic expression and gene-set overlap

For extraembryonic analyses, DNA Tape-positive cells were compared with Tape-negative (below-threshold) cells of the same annotation in E7.5 extraembryonic ectoderm and visceral endoderm, E8.5 ExVE, and EiTiX8 ExVE-like populations. We assessed donor-associated SNP support, overlap of upregulated genes, and enrichment of EMT and epiblast-associated programs (*42*).

For these expression comparisons, embryo Tape positivity required the recovery of at least eight edited Tape; we classified other cells as Tape-negative. For EiTiX8, Tape positivity required at least one recovered Tape (given this permissive threshold, we additionally restricted analyses to cells with doublet scores <0.15 as a conservative quality-control measure). We reclustered each compartment independently in Seurat 5.3.0 using log-normalized RNA, 2,000 variable genes, 20 PCs and resolution 0.5. Two-sided Wilcoxon tests compared Tape groups; differential expression required Bonferroni-adjusted P <0.05 across all RNA-assay features and |avg_log2FC| >0.5. Fold-change correlations assessed sensitivity to the stricter doublet filter. These pooled-cell tests lacked embryo-level replication.

### Independent epiblast signature scoring

We used an independent 500-gene epiblast signature derived from E5.25 reference GSM8352336 (GSE270803) *(42)*. We used two-sided, tie-corrected Wilcoxon tests with Benjamini–Hochberg adjustment. Signature genes required epiblast-versus-rest FDR <0.01, log₂ fold change >1, detection in ≥25% of epiblast cells and a detection difference ≥15 percentage points. Each comparison with VE, ExE/trophoblast and parietal endoderm additionally required FDR <0.01 and log₂ fold change >0.5. We excluded mitochondrial, ribosomal and cell-cycle genes.

### Gene set enrichment analysis

Preranked enrichment used fgseaMultilevel (fgsea 1.28.0; R 4.3.3) *(82)*. Extraembryonic rankings included all genes detected in at least three baseline cells, ordered by log₂ fold change in mean counts per 10,000 with a pseudocount of 0.1. Enrichment required Benjamini–Hochberg FDR <0.05 within each compartment, filtering condition and collection. Uptake/phagocytosis terms retained full-collection correction.

### Donor-associated RNA SNP analysis

We selected five RNA SNPs using E7.5 ExVE with fewer than eight edited Tapes as the host reference and embryonic cells excluding VE and ExEct as the donor reference, with a deterministic 80% training/20% held-out split. Greedy selection maximized coverage across references. The panel comprised H2az2 chr11:6378306 C>T, Uqcr10 chr11:4652120 C>T, Bod1 chr11:31615374 T>C, Rars chr11:35708261 A>G and Actr2 chr11:20013048 T>C (one-based GRCm39). Reference alleles were host-associated at all five sites. Cellsnp-lite 1.2.3 *(83)* counted allele UMIs using CB/UB tags, mapping quality ≥20 and aligned read length ≥30. The donor-associated fraction equaled donor-allele UMIs divided by donor-plus host-allele UMIs across the panel, requiring ≥5 observations across ≥2 genes. Scores >0.75, <0.25 or between these thresholds defined donor-like, host-like or intermediate cells. We applied the panel to E7.5 VE, E7.5 ExEct and E8.5 ExVE; EiTiX8 had no SNP scores.

### Spatial mESC-lineage-associated ExVE scoring analysis

Whole-mouse-gastrula Stereo-seq sections with three-dimensional spatial coordinates and cell-type annotations (GSE278603; embryonic days E7.5; two replicate sections per stage) (*72*) were analyzed in Python (NumPy, pandas, AnnData). For each cell, a mESC-lineage-associated ExVE score was computed as the mean log-normalized expression (log1p of library-size–normalized counts) of a set of gene niche signature (**table S5**), defined as the intersection of a curated niche gene list with the genes detected in the dataset.

For each section, an anterior–posterior (AP) reference line was fitted in the x–y plane by scanning orientations over [0, 180°). At each orientation, the offset was set to the median projection of caudal epiblast cells, and the orientation minimizing the cell-count imbalance across the line for the anterior landmark was selected. Visceral and parietal endoderm cells were grouped into layers along the z axis, perpendicular to the AP reference line, and parallel to that line. Each axis spanned the full section and was divided into 3, 5, 10, 15 or 20 equal-width bins. Mean endoderm scores were calculated per bin. Complementary profiles were generated from within-section gene-wise z scores and averaged across sections on a common relative-position grid. Landmark definitions, line parameters, bin-level cell counts and mean scores are provided in **table S8**.

Neighborhood analyses used three-dimensional Euclidean distances. ExVE scores were compared between cells with and without epiblast neighbors across the plotted radii within each replicate. Local epiblast and caudal epiblast fractions within 100 μm and distance to the nearest epiblast cell were analyzed separately. Scores were interpreted as expression-state proxies.

## Supplementary figure legends

**Fig. S1.**
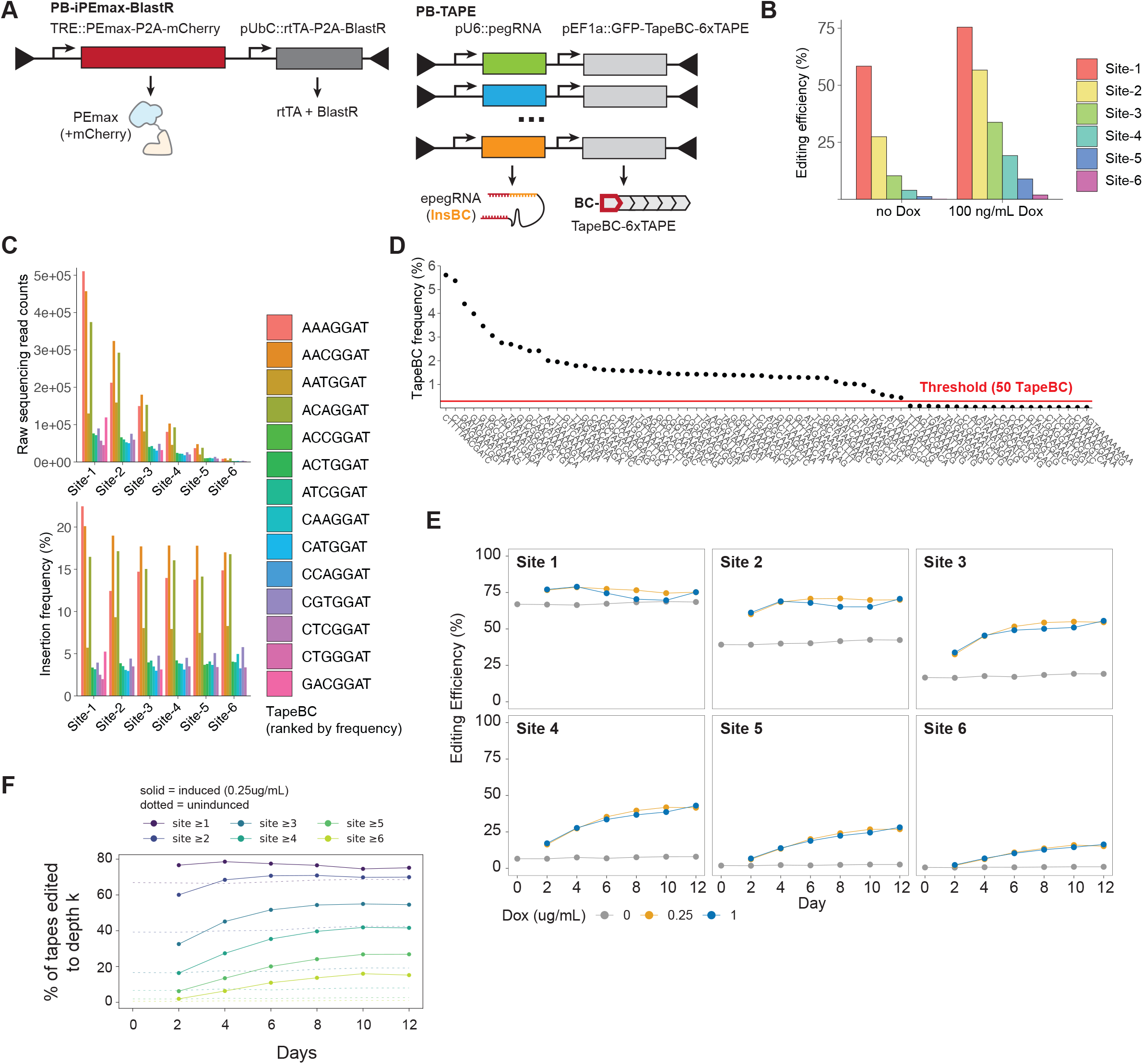
Characterizing DNA Typewriter-based Lineage Tracing mESCs. **(A)** Genetic components used to generate Lineage Tracing mESCs. PB-Tape constructs encode pegRNAs with NNNGGA insertion sequences (InsBCs) and GFP transcripts carrying NNNNNAANNNNN Tape barcodes (TapeBCs) and six sequential recording sites. PB-iPEmax-BlastR encodes doxycycline-inducible PEmax-P2A-mCherry and rtTA-P2A-BlastR. **(B)** Tape editing frequencies across 6 insertion sites on 6xTape for the clonal mESC used for tetraploid complementation after two days with or without 100 ng ml^−1^ doxycycline. **(C)** Raw read count (top) and frequencies (bottom) of insertion sequences (InsBCs) observed across 6 insertion sites. **(D)** Frequencies of detected TapeBC in the clonal mESC. TapeBC with frequencies above a set threshold (>0.4%) were used in subsequent analysis. **(E)** Site-specific editing frequencies during a 12-day mESC culture time course with 0, 0.25, or 1 μg ml^−1^ doxycycline, measured by bulk genomic DNA amplicon sequencing. **(F)** Fraction of Tapes reaching each cumulative editing depth during the time course. Solid lines indicate 0.25 μg ml^−1^ doxycycline; dotted lines indicate untreated cultures. Colors denote editing depth.

**Fig. S2.**
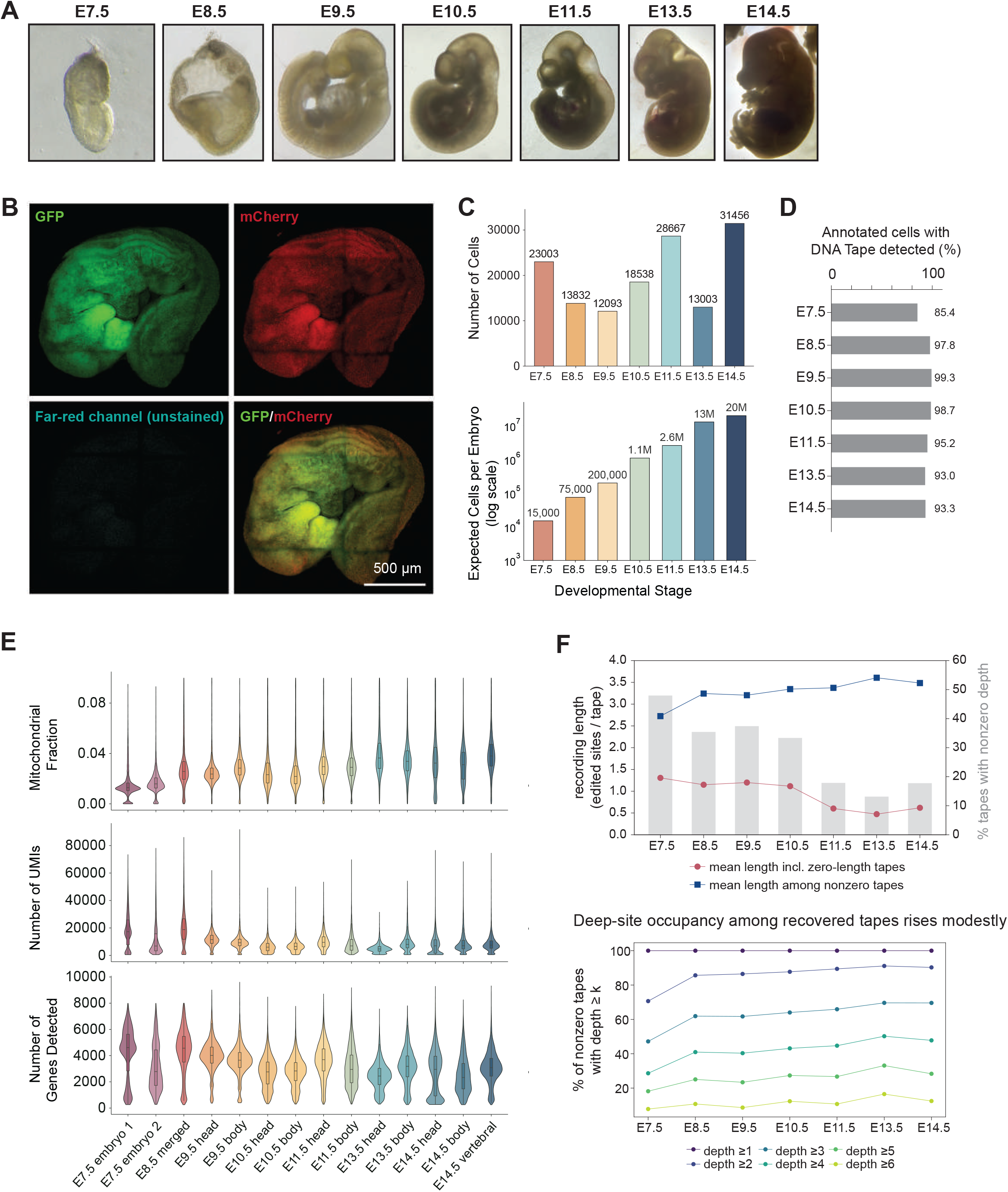
Quality control measures of transcriptome and genomic recordings in DNA Tape. **(A)** Representative images of embryos collected from E7.5 to E14.5. Images are shown not to the physical scale. **(B)** Reporter fluorescence in a lineage-recording embryo: GFP, mCherry, unstained far-red, and merged images. Scale bar, 500 μm. **(C)** (Upper) The number of single cells retained at each embryonic stage after quality control. (Lower) Estimated total cell numbers per embryo, shown on a logarithmic scale. **(D)** Fraction of transcriptome-annotated embyronic cells in which at least one DNA Tape was detected, per stage (extraembryonic is not included). **(E)** Mitochondrial fraction, number of UMIs (cDNA molecules) and genes detected across 14 sample groups. **(F)** Upper, mean sequential recording length across all Tape entries (red) or nonzero entries only (blue); gray bars indicate the fraction of entries with nonzero depth (right axis). Lower, fractions of nonzero entries reaching each editing depth. Zero-length entries can represent unedited or unrecovered Tapes.

**Fig. S3.**
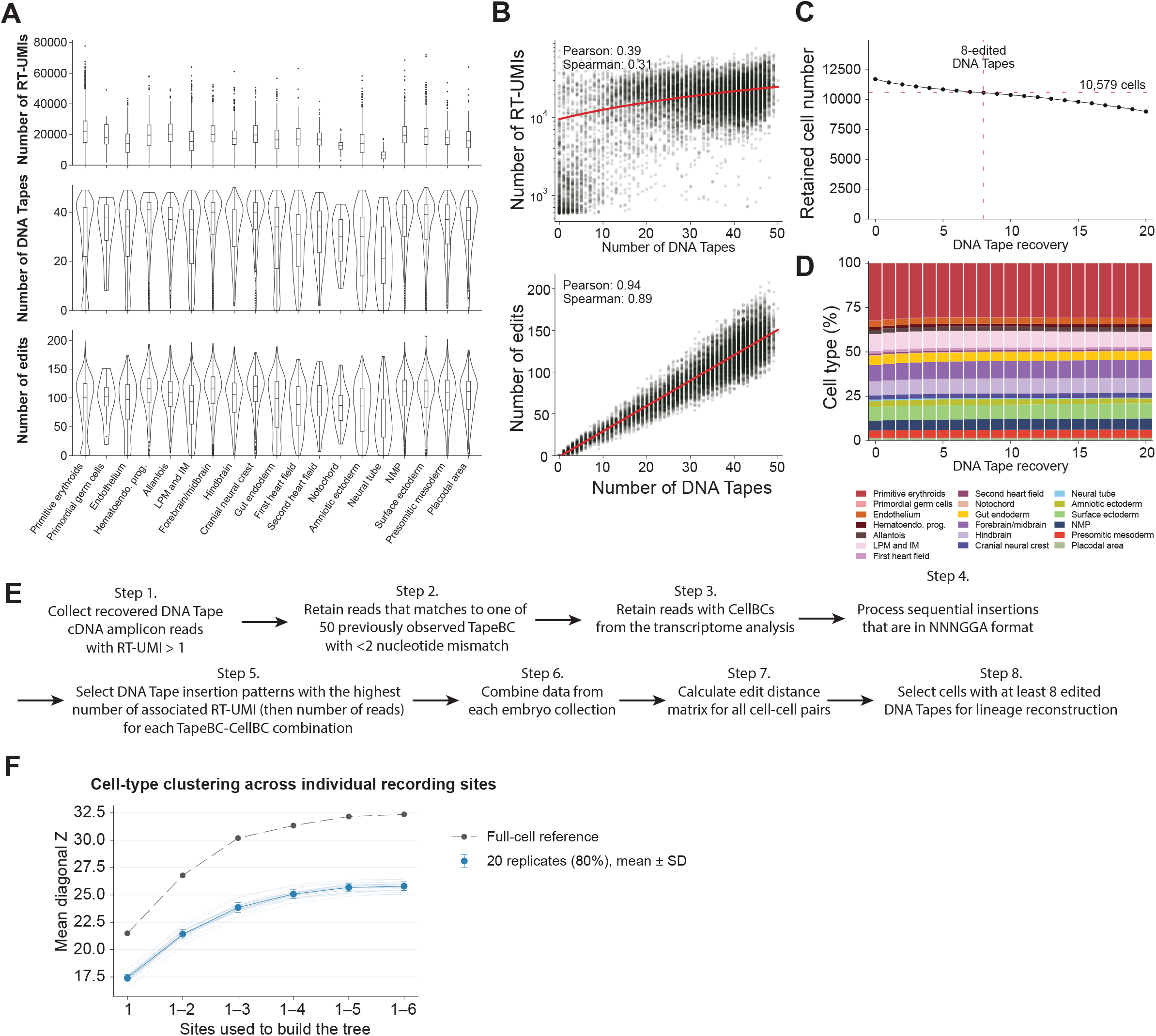
Processing genomic recordings recovered from single cell DNA Tapes. **(A)** Characterizing single cell recovery across different cell type annotations. (Top) Number of RT-UMIs, which are unique molecular identifier sequences attached to cDNA during reverse transcription, for counting the cDNA recovery efficiencies. (Center) Number of DNA Tapes recovered in each cell, out of the previously curated set of 50 DNA Tapes in the clonal mESC used. (Bottom) Number of recovered edits in DNA Tapes in each cell. **(B)** (Top) Correlation of number of recovered DNA Tapes and the number of RT-UMIs in each cell. (Bottom) Correlation of number of recovered DNA Tapes and the number of insertion barcode edits detected across all DNA Tapes per cell. Pearson and Spearman correlations are shown. **(C)** Number of retained cells across different thresholds of DNA Tape recovery. A threshold of ≥8 edited Tapes recovered per cell was used to select a set of 10,579 non-ExVE cells for the E8.5 embryo lineage reconstruction data set. **(D)** Cell type compositions across different thresholds of DNA Tape recovery. **(E)** Overview of the sequencing data processing scheme for recovered single-cell DNA Tapes. **(F)** Contribution of successive recording positions to mean within-type PATH Z scores, using cumulative Sites 1 through 1–6. Points and error bars show mean ± SD across 20 paired 80% cell subsamples without replacement; dashed values show the full cohort. Trees were rebuilt for each site set and subsample.

**Fig. S4.**
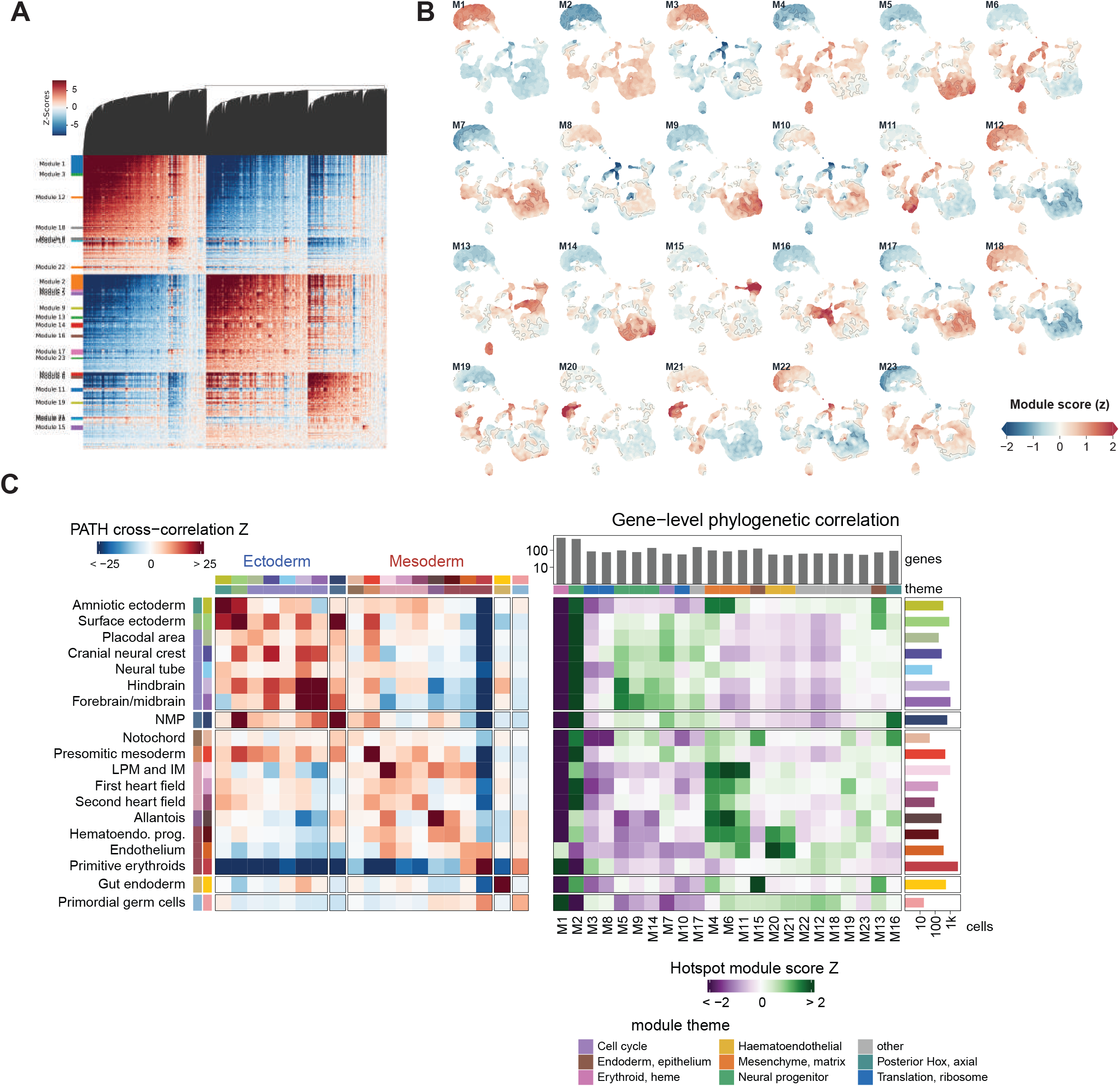
Lineage-associated transcriptional programs in the E8.5 embryo. **(A)** Genes with significant (FDR < 0.05) lineage autocorrelation are grouped into 23 gene modules based on pairwise lineage correlation. The heatmap shows Hotspot local-correlation Z scores. **(B)** Module summary scores plotted against the UMAP of the E8.5 embryo transcriptomic expression data. Scores are standardized across cells. **(C)** Cell-type PATH associations alongside module-score profiles. Module scores are standardized across modules within each cell type. Bars indicate cell and module-gene counts; module colors identify functional themes.

**Fig. S5.**
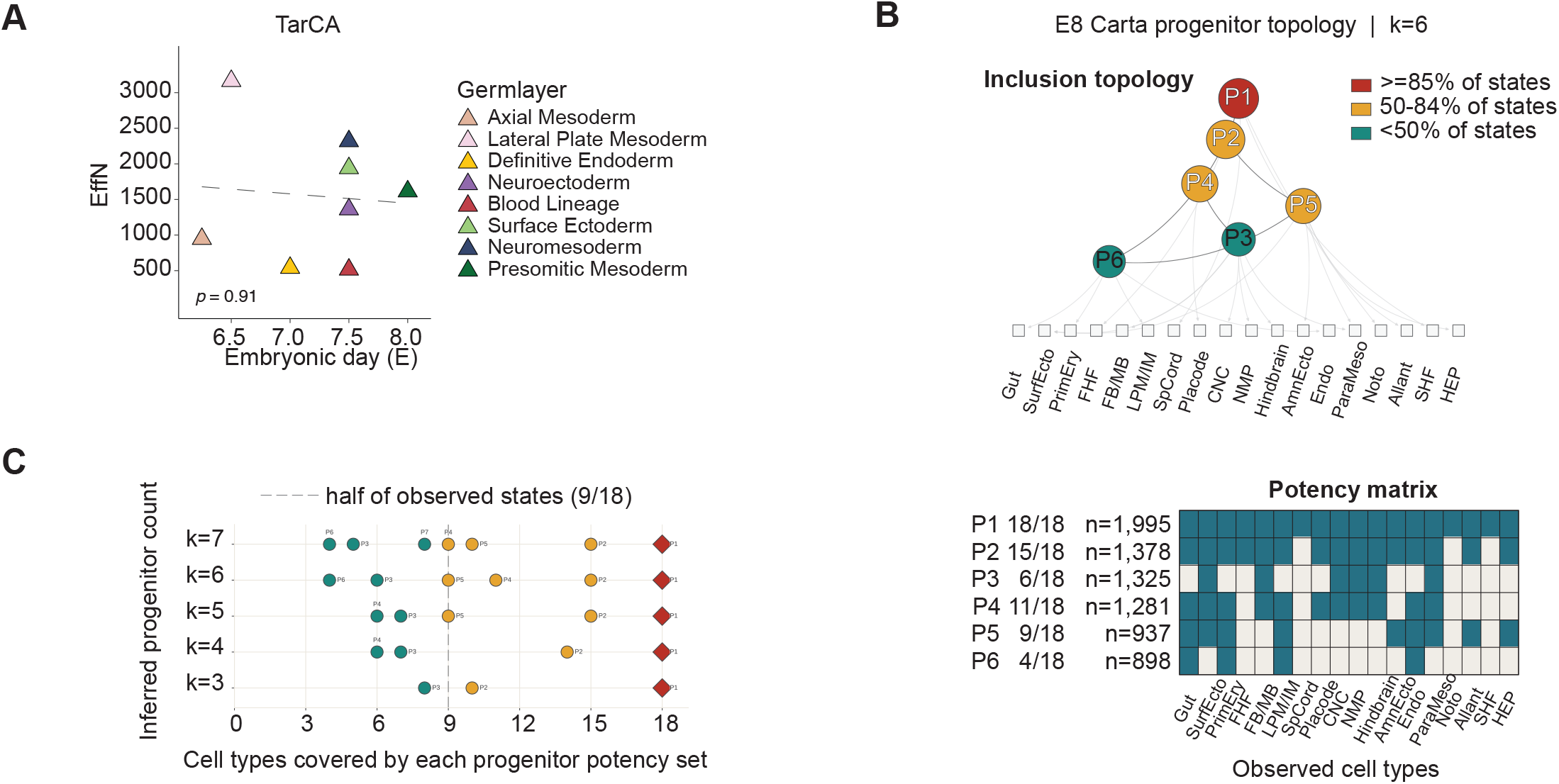
Progenitor diversity and fate hierarchies inferred from lineage trees. **(A)** Relationship between TarCA effective progenitor number and reference developmental time (Spearman ρ = 0.05, P = 0.91). Dashed line, linear fit. **(B)** CARTA potency-set analysis of 18 terminal E8.5 cell types with six selected sets (P1–P6). Upper, inclusion relationships among inferred sets and their associated terminal types. Node colors indicate the fraction of the 18 observed types covered by each set. Lower, binary membership matrix for the six sets. Cell-type abbreviations are as labeled in the panel: SurfEcto, surface ectoderm; PrimEry, primitive erythroid cells; FHF, first heart field; FB/MB, forebrain/midbrain; LPM/IM, lateral plate/intermediate mesoderm; SpCord, spinal cord; Placode, placodal area; CNC, cranial neural crest; NMP, neuromesodermal progenitors; AmnEcto, amniotic ectoderm; Endo, endothelium; ParaMeso, paraxial mesoderm; Noto, notochord; Allant, allantois; SHF, second heart field; HEP, hematoendothelial progenitors. **(C)** Dependence of inferred potency hierarchies on the number of selected sets, from three to seven. Each point represents one Carta-inferred progenitor. The dashed line marks half of all observed cell types, 9/18. Sets were optimized using the local CARTA-objective implementation (**Methods**). Most inferred progenitors cover many cell types, indicating that Carta mainly recovers broad potency sets in this dataset.

**Fig. S6.**
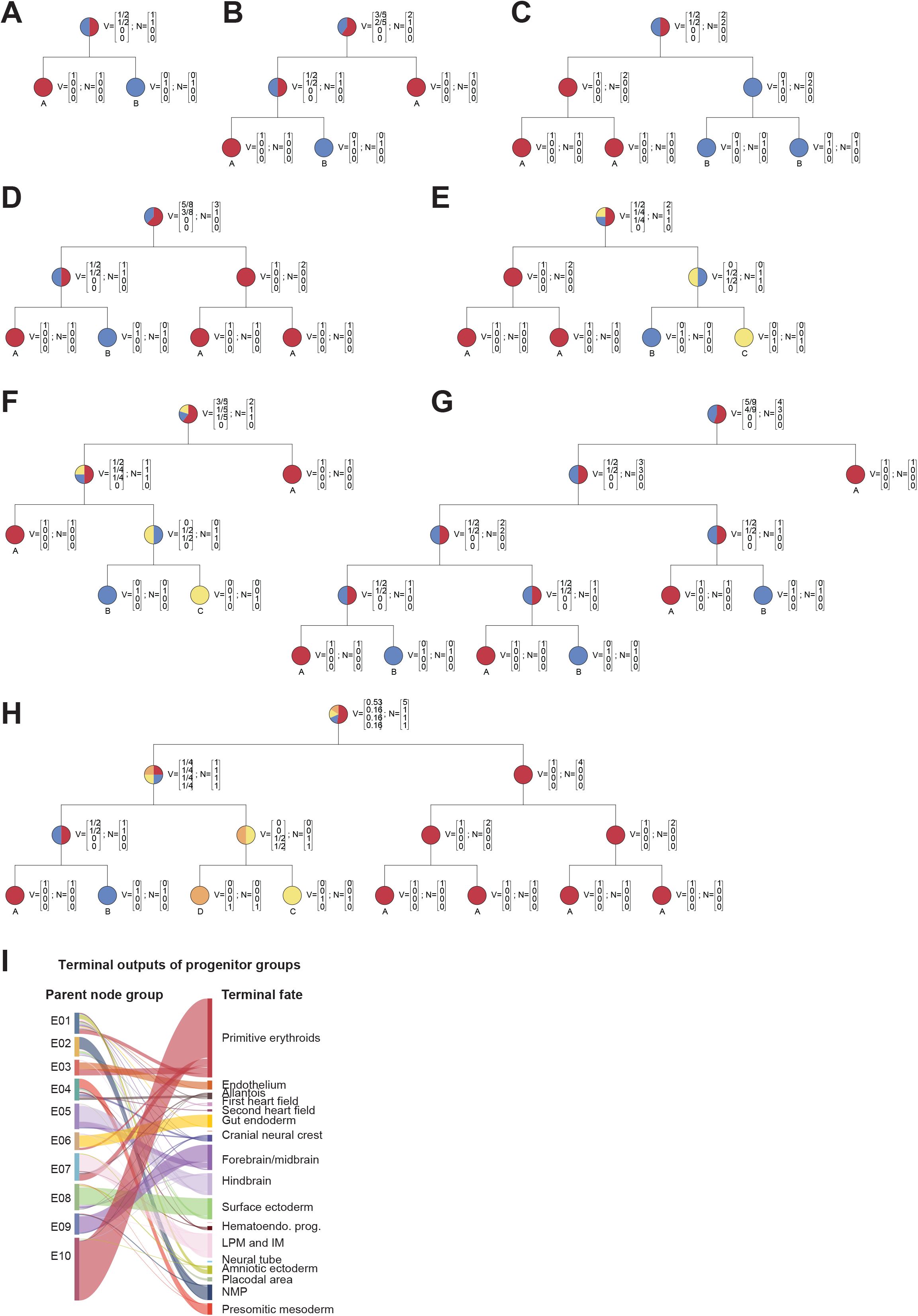
FateVec integrates lineage topology with descendant fate composition. Worked examples illustrate how the placement and expansion of fate-restricted branches influence ancestral fate profiles. **(A,C)** Balanced two-fate examples with equal terminal counts and equal root fate components, before **(A)** and after **(C)** expansion of fate-restricted branches. **(B,D,G)** Two-fate examples with unequal descendant counts and different branching structures, illustrating their effects on the root fate vector. **(E,F)** Identical terminal counts across three fates produce different root vectors when the branching topology differs. **(H)** Expansion of a fate-restricted branch attached to a four-fate lineage. Node labels show descendant counts and fate components. Internal vectors combine fate-specific descendant-count weighting with normalization across fates. **(I)** Correspondence between the E8.5 node-profile groups E01–E10 in **Fig. 2C** and terminal cell types. Each terminal cell is assigned to the group containing its immediate parent node. Ribbon widths indicate cell counts; colors denote terminal cell types. Links containing fewer than 20 cells are hidden, while bar totals and ribbon scaling retain all 10,579 cells.

**Fig. S7.**
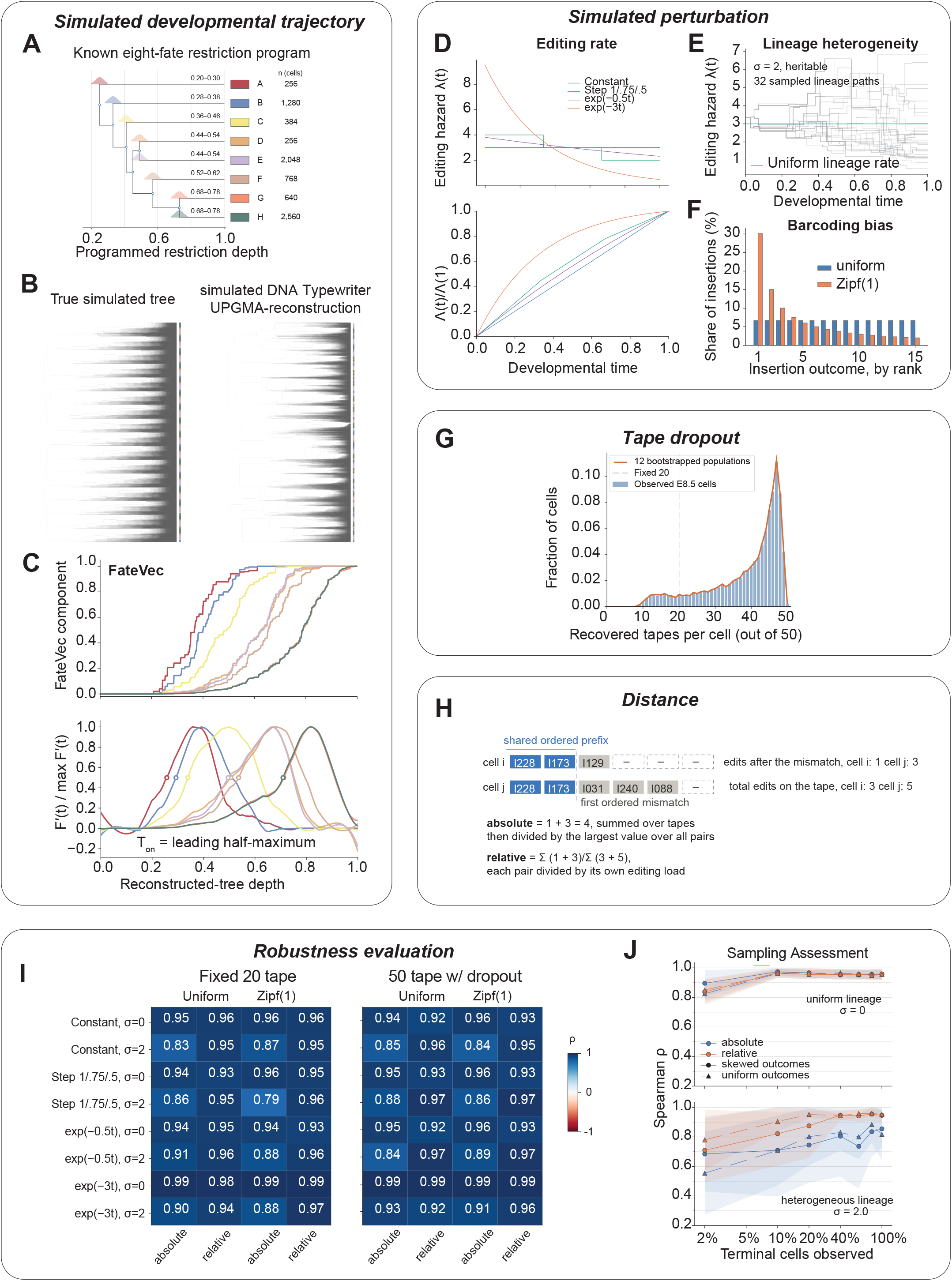
Recovering developmental timing from imperfect molecular records. A known developmental program provides a benchmark for testing how recording kinetics, lineage heterogeneity, Tape recovery, and cell sampling affect inferred restriction timing. **(A)** Programmed tree with eight fates, 16 progenitor modules, and 8,192 terminal cells; shading marks restriction windows. **(B)** Ground-truth lineage tree generated from the fate hierarchy and restriction windows in **(A)**, with stochastic branch durations (Left), and its reconstruction from simulated DNA Typewriter records by UPGMA using relative ordered-tape distances (Right). Records accumulated along the ground-truth tree under a constant editing schedule, heritable editing-rate variation (σ = 0.66), and Zipf(1) insertion outcomes. Tip colors denote fate; fate labels were overlaid after reconstruction. **(C)** FateVec trajectories inferred from the reconstructed tree in **(B)**. Upper panel: baseline-normalized mean fate components across reconstructed-tree depth. Lower panel: smoothed derivatives normalized to their respective maxima. Circles mark restriction onset (*T_on_*), defined as the first crossing of half the global derivative maximum for each fate. Colors correspond to **(A)**. **(D)** Temporal editing-rate profiles: constant, stepwise decline (relative rates of 1, 0.75, and 0.5 over three equal intervals), and exponential declines proportional to exp(−0.5t) or exp(−3t), where t is normalized developmental time. Profiles have the same integrated editing drive. Upper, instantaneous editing rates; lower, normalized cumulative editing drive. **(E)** Heritable editing-rate variation (σ = 2) across 32 sampled lineage paths; green line, uniform-rate reference. **(F)** Uniform versus nonuniform insertion probabilities across 15 possible sequences; the latter follow a Zipf distribution with exponent 1. **(G)** Empirical E8.5 Tape recovery and 12 bootstrapped recovery populations for the 50-Tape dropout model, compared with fixed 20-Tape recording. **(H)** Absolute and relative ordered-prefix distances. Starting at the first insertion mismatch on each Tape, the remaining edit counts in the two cells are summed across Tapes. Absolute distances are scaled by the largest pairwise value; relative distances are divided by the summed editing load of the corresponding cell pair. **(I)** Timing-order recovery across recording conditions, summarized by mean Spearman correlation over 12 replicates. **(J)** Recovery of fate-bias timing order at the plotted terminal-cell sampling fractions, spanning 2–100%. Upper, uniform lineage editing rates (σ = 0); lower, heritable rate variation (σ = 2), both with a constant baseline editing schedule. Spearman correlations compare inferred *T_on_* with programmed restriction centers across the eight fates. Colors distinguish absolute and relative distances; symbols distinguish uniform and Zipf(1) insertion outcomes. Lines and shading show mean ± SD across 12 replicates.

**Fig. S8.**
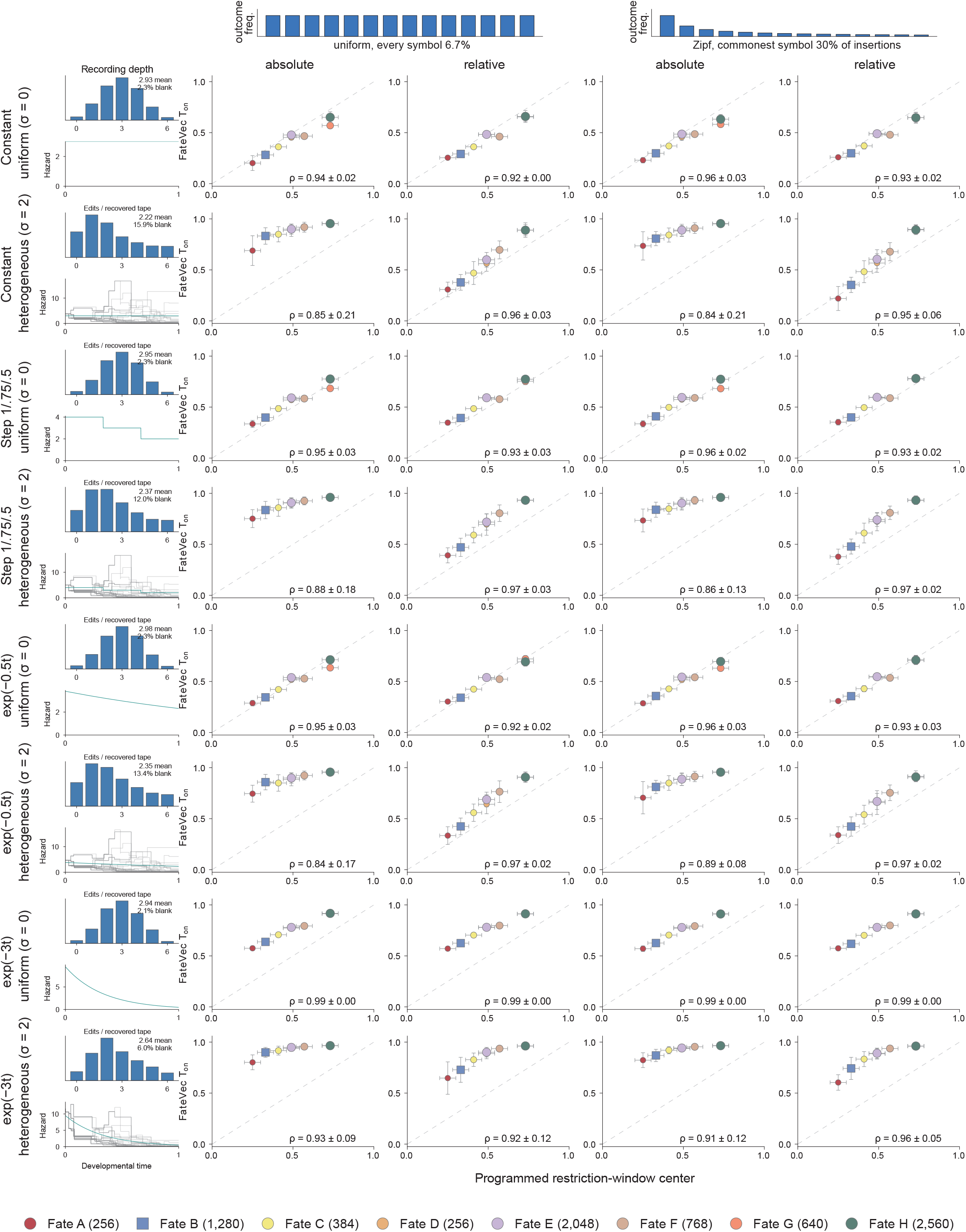
Recovery of developmental fate order across recording regimes. Fate-level comparisons resolve how recorder behavior affects inferred timing across lineages with different abundances and programmed restriction windows. Programmed restriction centers versus *T_on_* under four editing schedules and two lineage-heterogeneity levels (σ = 0 or 2). Columns combine absolute or relative distance with uniform or Zipf(1) insertion outcomes. Points show means across 12 replicates; vertical bars, SD; horizontal bars, programmed windows of ±0.05. Point size scales with log cell abundance. Recording distributions and editing-rate paths summarize the underlying conditions. Annotations give mean ± SD of within-replicate Spearman correlations across eight fates, retaining the shared timing centers of D/E and G/H.

**Fig. S9.**
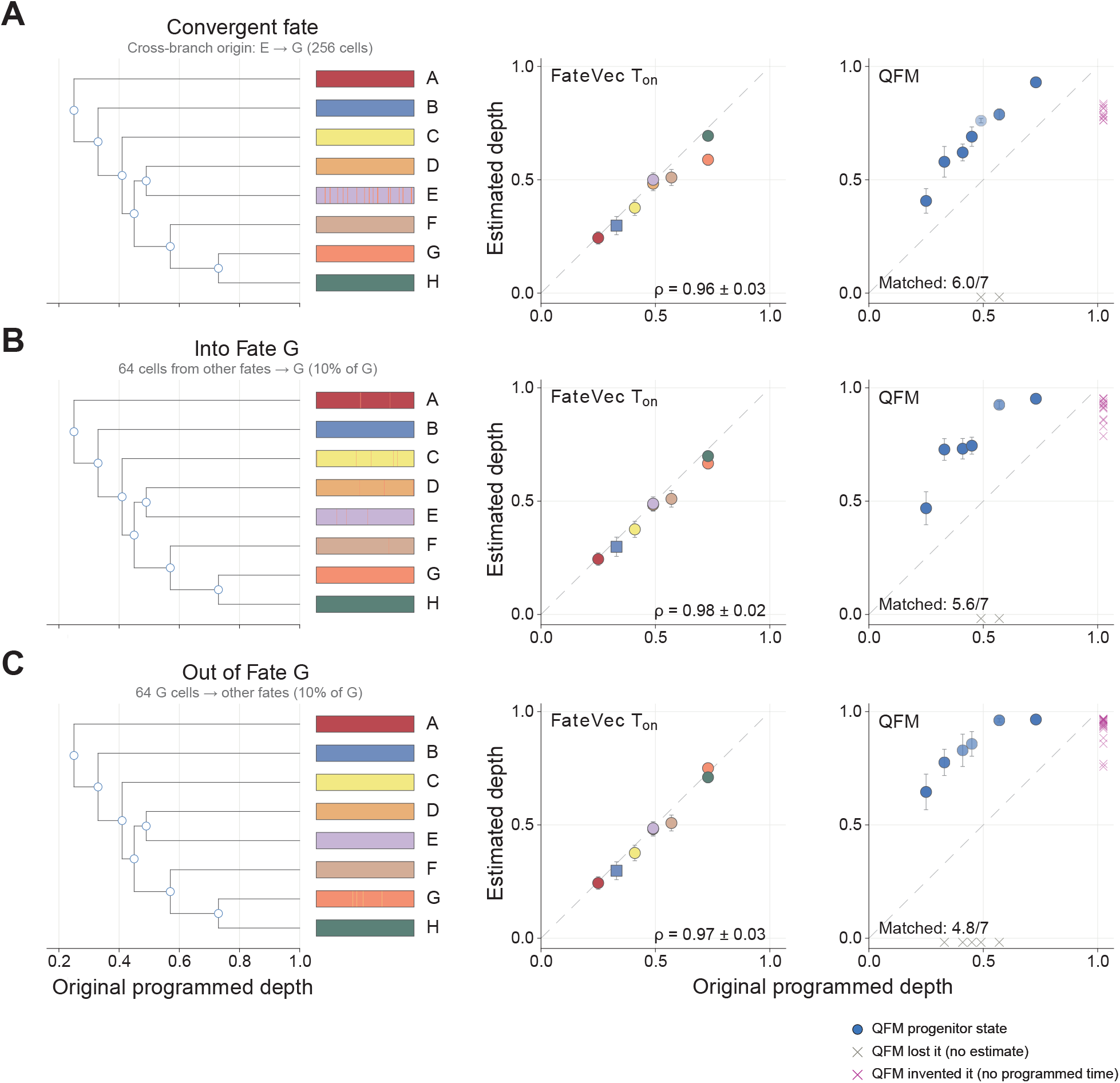
Fate-timing inference under convergent and perturbed cell identities. Holding the reconstructed tree fixed isolates the sensitivity of timing estimates to changes in terminal fate assignment. **(A)** Convergent identity, introduced by relabeling 256 E-subclade cells as G. **(B,C)** Dispersed label perturbations involving 64 non-G cells assigned to G **(B)** or 64 G cells assigned to other fates **(C)**. Left, fate program and terminal labels; middle, FateVec terminal-fate *T_on_*; right, QFM progenitor-state commitment times. Points and bars show mean ± SD across ten paired replicates, with the same tree used across methods and perturbations. QFM states were matched to seven programmed potency sets. Opacity indicates recovery frequency; means use replicates recovering each state. Gray crosses mark missing original states; magenta crosses mark unmatched states. Axes refer to the original program, before perturbation.

**Fig. S10.**
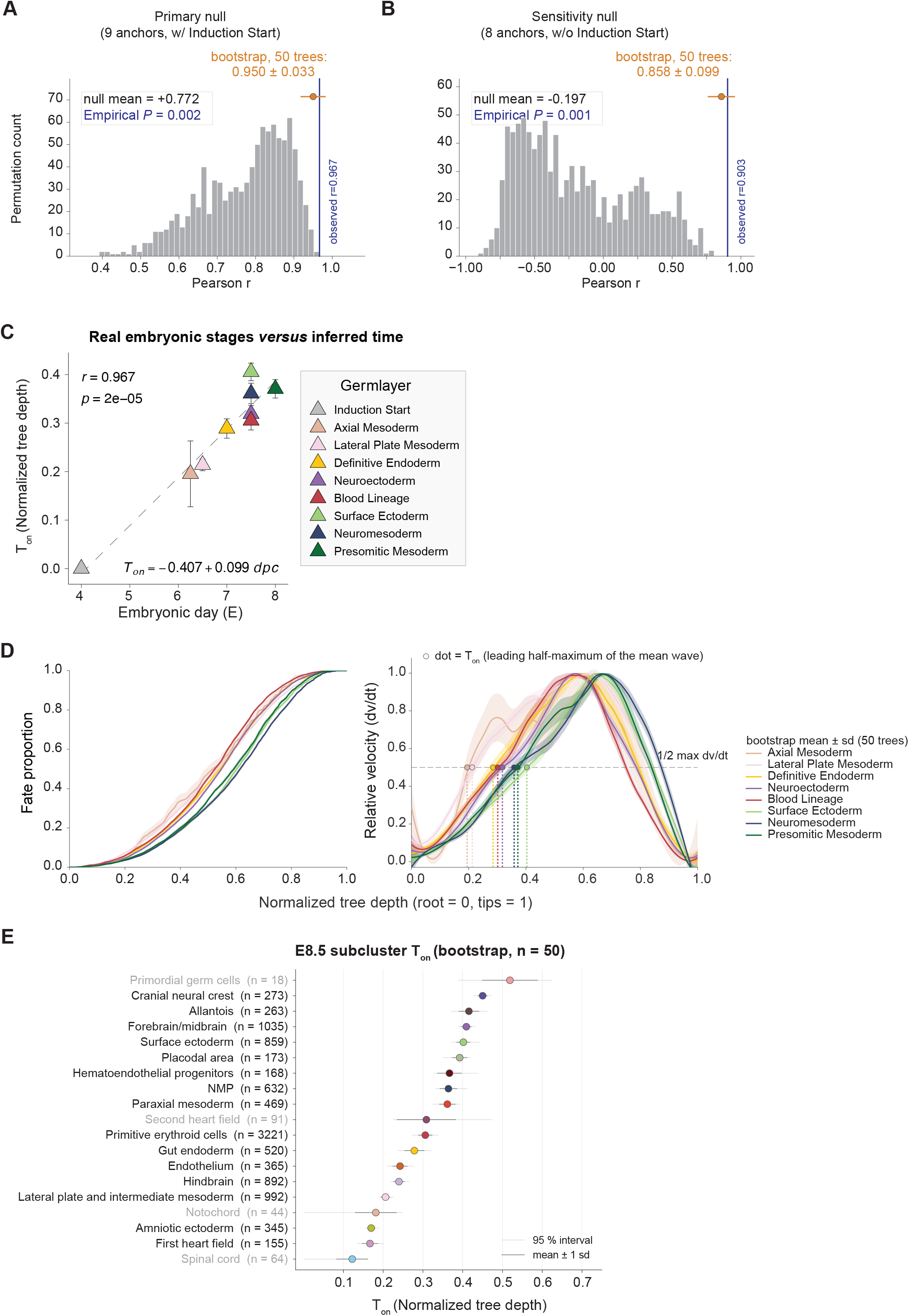
Calibrating lineage-derived fate timing to developmental chronology. Developmental references, label permutations, and cell subsampling assess the calibration and sampling sensitivity of inferred restriction onset. **(A, B)** Correlation of *T_on_* with reference time, including (**A**; nine points) or excluding (**B**; eight points) the E4.0 reference. Histograms show 1,000 count-preserving label permutations; observed r = 0.967 and 0.903, with empirical P = 0.002 and 0.001. Orange summaries show mean ± SD across 50 subsamples. **(C)** Calibration against the developmental reference times used in **Fig. 2E**. Points and error bars show mean *T_on_* ± SD across 50 trees reconstructed from 80% cell subsamples without replacement. Dashed line, linear fit including the reference anchor assigned to E4.0. **(D)** FateVec trajectories (left) and peak-normalized derivatives (right) for eight broad developmental programs across 50 tree reconstructions from 80% cell subsamples without replacement. Bands show SD; dots mark the leading half-maximum onset of the mean derivative. **(E)** Mean E8.5 cell-type onset estimates with SDs and empirical 2.5th–97.5th percentile intervals. Counts refer to the reference tree.

**Fig. S11.**
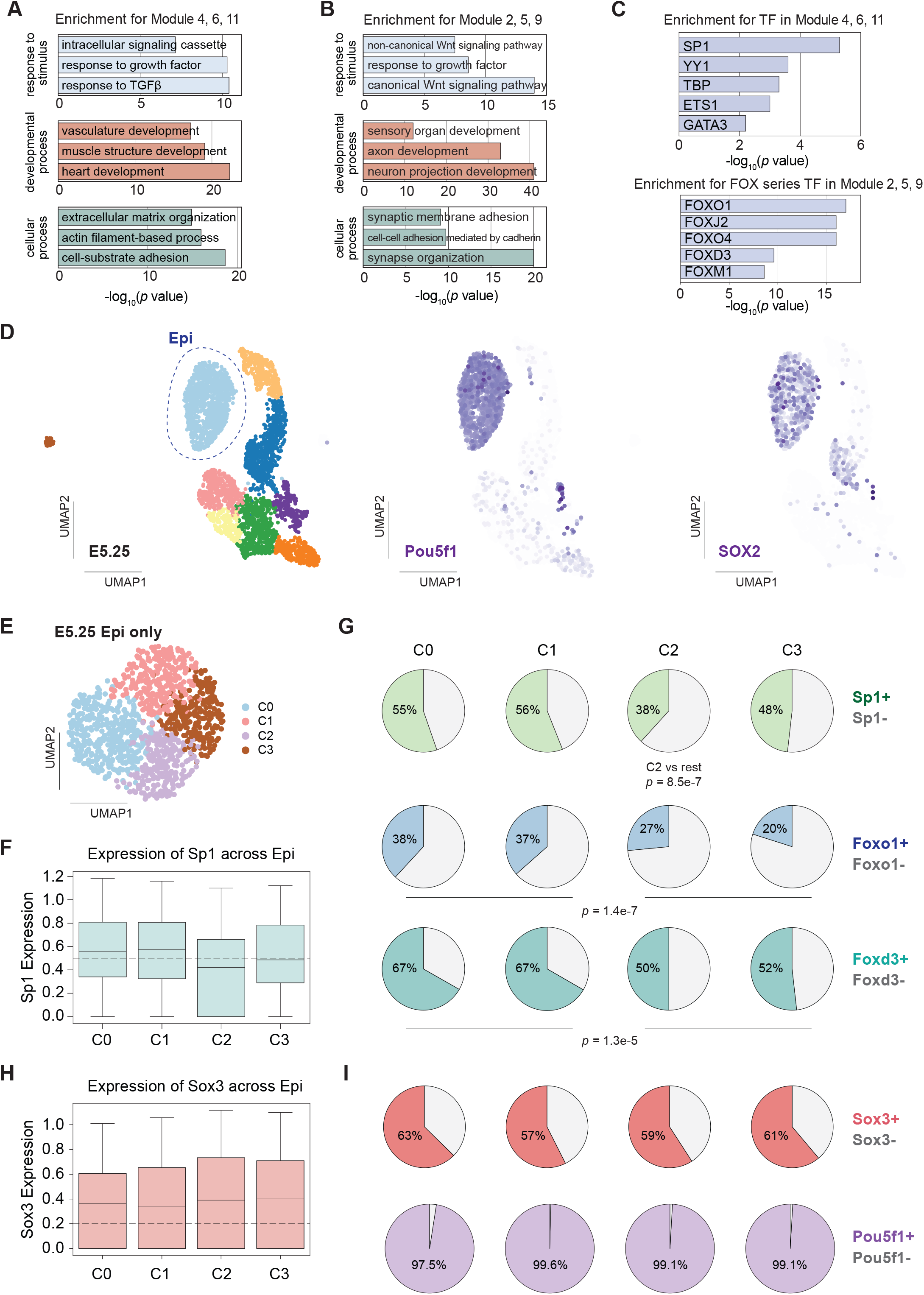
Early transcription factor heterogeneity within the E5.25 epiblast. **(A)** Gene Ontology (GO) Biological Process enrichment analysis of genes in co-expression Modules 4, 6, and 11. Bar length indicates enrichment significance; representative top terms are shown. **(B)** GO Biological Process enrichment analysis of genes in co-expression Modules 2, 5, and 9, displayed as in **A**. **(C)** Transcription factor (TF) target enrichment analysis for Modules 4/6/11 (top) and Modules 2/5/9 (bottom), highlighting distinct TF programs associated with each module group. **(D)** UMAP embedding of single-cell transcriptomes from E5.25. Epiblast (Epi) cells are circled based on canonical markers; feature plots of Pou5f1 and Sox2 are shown to the right. **(E)** Unbiased re-clustering of E5.25 epiblast cells, revealing four transcriptionally distinct subclusters (C0–C3) in UMAP space. **(F)** Sp1 expression across epiblast subclusters (C0–C3), shown as boxplots of expression (center line, median; box, interquartile range; whiskers, 10–90th percentiles). **(G)** Proportions of TF-positive and TF-negative cells across epiblast subclusters for Sp1, Foxo1, and Foxd3 (pie charts; percentages indicated). Statistical significance for differences in TF-positive fractions between clusters is annotated as shown. **(H)** Sox3 expression across epiblast subclusters (C0–C3), shown as boxplots of normalized expression (center line, median; box, interquartile range; whiskers, 10–90th percentiles). **(I)** Fractions of Sox3-positive and Pou5f1-positive cells in each epiblast subcluster (pie charts; percentages indicated).

**Fig. S12.**
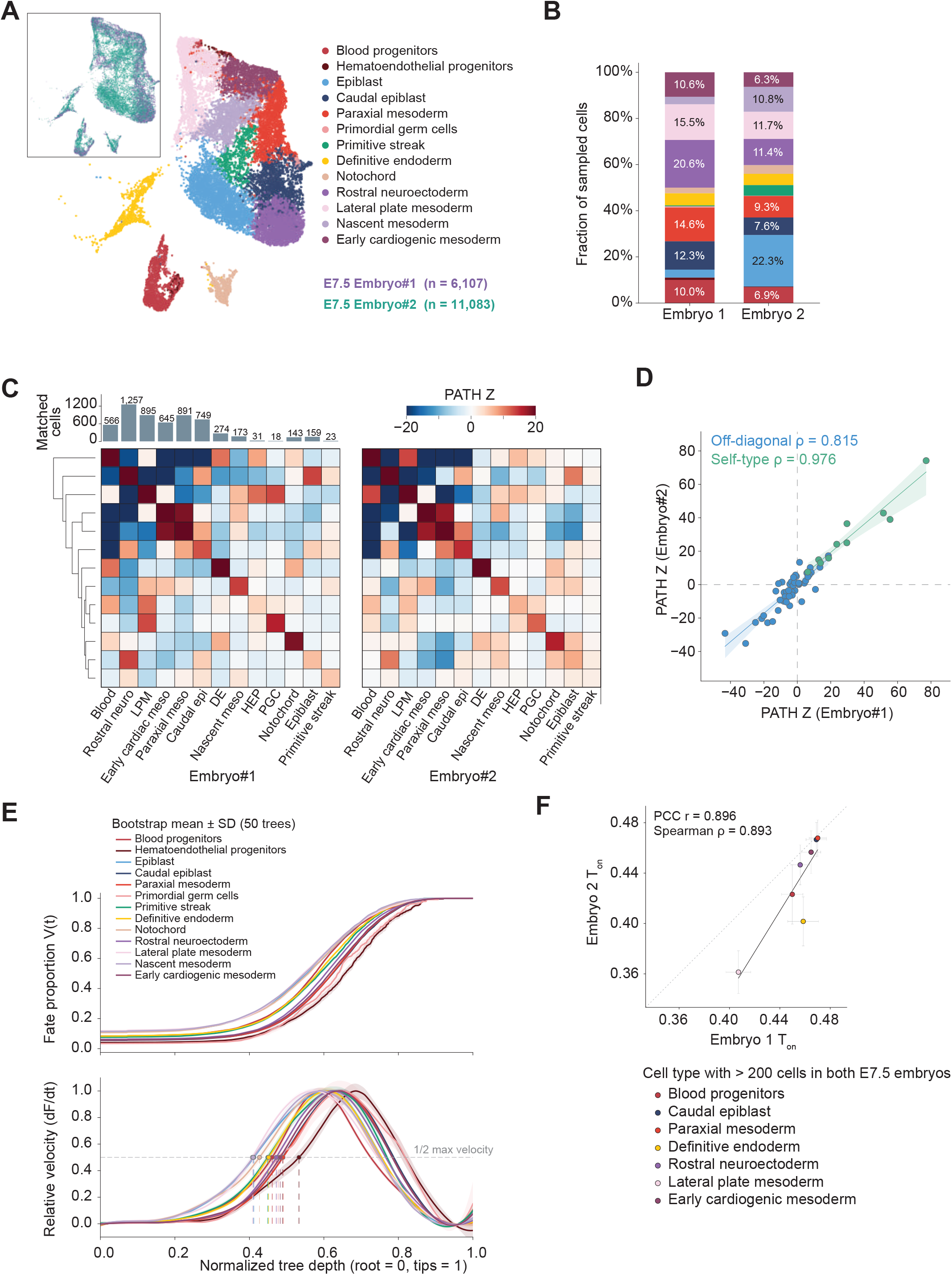
Reproducible lineage organization and fate timing across E7.5 embryos. **(A,B)** UMAP of single cell transcriptomic data **(A)** and cell-type composition **(B)** of 13 retained populations: 6,107 and 11,083 cells, with 5,873 and 8,922 lineage-tree cells in two E7.5 mouse embryos. **(C)** Mean symmetrized PATH Z scores across 100 subsampling iterations, with cell counts matched between embryos for each of the 13 cell types. Each iteration sampled the smaller available count per type from both embryos without replacement, yielding 5,824 cells per embryo. **(D)** Agreement of within-type and between-type associations for ten cell types with ≥100 cells per embryo (Spearman ρ = 0.976 and 0.815; 10 types and 45 pairs). Lines and bands indicate linear fits and bootstrap intervals. **(E)** Pooled FateVec trajectories and peak-normalized derivatives over 50 stratified 80% subsamples; shading, SD. **(F)** Mean *T_on_* ± SD for seven types with >200 lineage-tree cells per embryo (Pearson r = 0.896; Spearman ρ = 0.893).

**Fig. S13.**
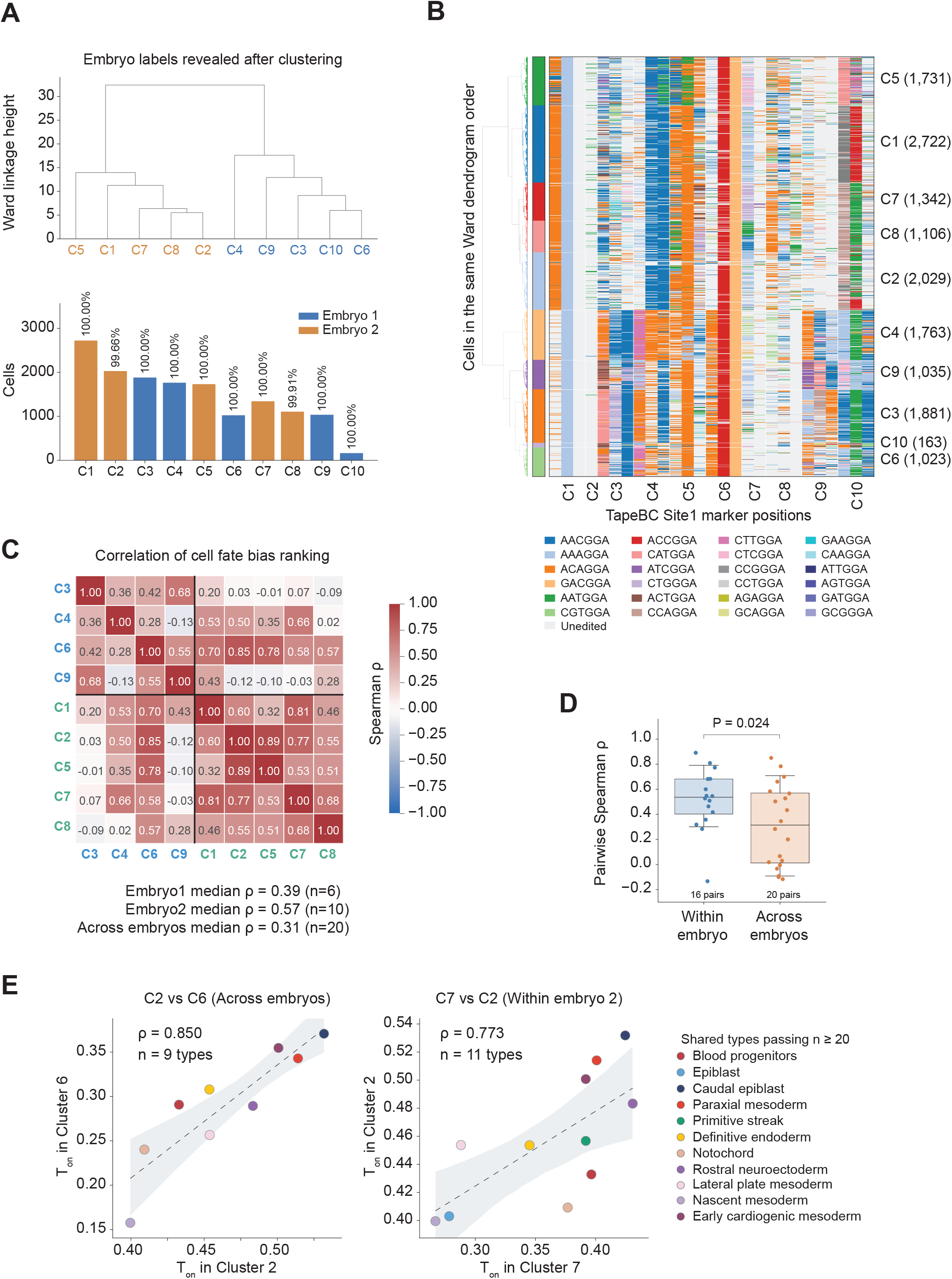
Clade-level variation in fate timing within and between E7.5 embryos. Clade-resolved comparisons examine whether embryo-level timing agreement extends to individual sampled lineage histories. **(A)** Ten clades (C1–C10) identified by Ward hierarchical clustering of Site-1 insertion profiles from pooled cells of two E7.5 embryos. Upper, dendrogram with clade labels colored by the predominant embryo, overlaid after clustering. Lower, cell counts by embryo within each clade; percentages indicate the contribution of the predominant embryo. **(B)** Clade-associated Site-1 insertion patterns. Up to three DNA Tapes per clade were selected for enrichment of a characteristic insertion sequence. Rows show cells in the dendrogram order of **(A)**; columns show selected Tapes, grouped by the clade for which they were selected. Colors denote insertion sequences; gray denotes the unedited category in the key. Labels on the right give clade identities and cell counts. **(C)** Pairwise Spearman correlations of mean *T_on_* across nine clades; the small C10 clade was excluded. Comparisons require ≥20 cells per type per clade and ≥5 shared types; estimates use 50 stratified 80% subsamples. **(D)** Distribution of the pairwise Spearman correlations in **(C)** for clades from the same or different embryos (16 and 20 pairs, respectively). Each correlation compares mean *T_on_* across shared cell types. Points represent clade pairs; boxes, median and interquartile range; whiskers, 10th–90th percentiles. P = 0.024 from a two-sided exact test comparing median correlations over 126 embryo-label permutations preserving the four-versus-five clade allocation. **(E)** Representative cross-embryo C2–C6 and within-embryo C7–C2 comparisons (ρ = 0.85 and 0.773, respectively).

**Fig. S14.**
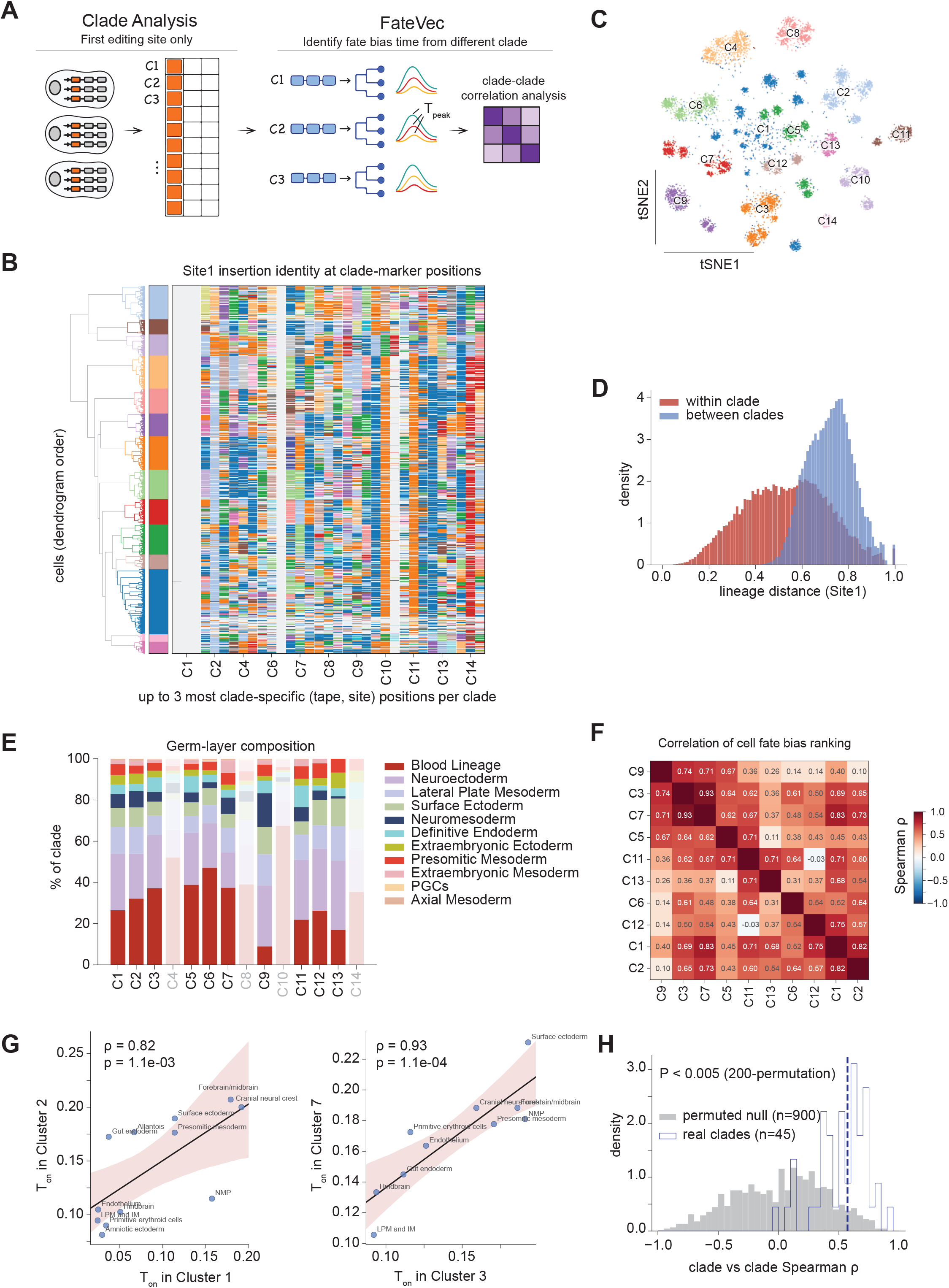
Concordant fate ordering across lineage clades in pooled E8.5 embryos. Site-1 records partition the pooled E8.5 cohort into clades for comparing the order of cell-type fate restriction across distinct lineage histories. **(A)** Clade-based analysis workflow. Site-1 insertion profiles define 14 clades by Ward hierarchical clustering. Lineage trees are reconstructed within each clade, and FateVec-derived *T_on_* values are compared across shared cell types. **(B)** Site-1 insertion patterns distinguishing the clades. Rows represent cells ordered by the Ward dendrogram; columns represent selected DNA Tapes, with up to three per clade chosen for enrichment of a characteristic insertion sequence. Heatmap colors indicate insertion identity; gray indicates missing records. The adjacent color strip marks clade membership. **(C)** t-SNE embedding of Site-1 insertion profiles, with each point representing a cell and colors indicating the 14 clades. **(D)** Distributions of pairwise Site-1 recording distances among 2,500 randomly sampled cells, comparing pairs within the same clade (red) or in different clades (blue). **(E)** Broad lineage composition of the 14 clades. Faded bars mark C4, C8, C10, and C14, excluded from the exploratory timing comparisons in **(F–H)** (**Methods**). **(F)** Pairwise Spearman correlations of *T_on_* across ten clades. **(G)** *T_on_* comparisons for C1 versus C2 (Left) and C3 versus C7 (Right). Lines and shading indicate linear fits and 95% bootstrap subsampling confidence intervals. Spearman ρ = 0.82 and 0.93, respectively; P values are two-sided. **(H)** Observed pairwise correlations from **(F)** (blue outline; 45 clade pairs) and a null distribution generated by shuffling cell-type labels within each clade while preserving tree topology and cell-type counts (gray; 900 correlations from 20 permutations). Dashed line, observed median ρ. P < 0.005 from a one-sided conditional permutation test of the median correlation using 200 permutations, with the ten-clade set held fixed.

**Fig. S15.**
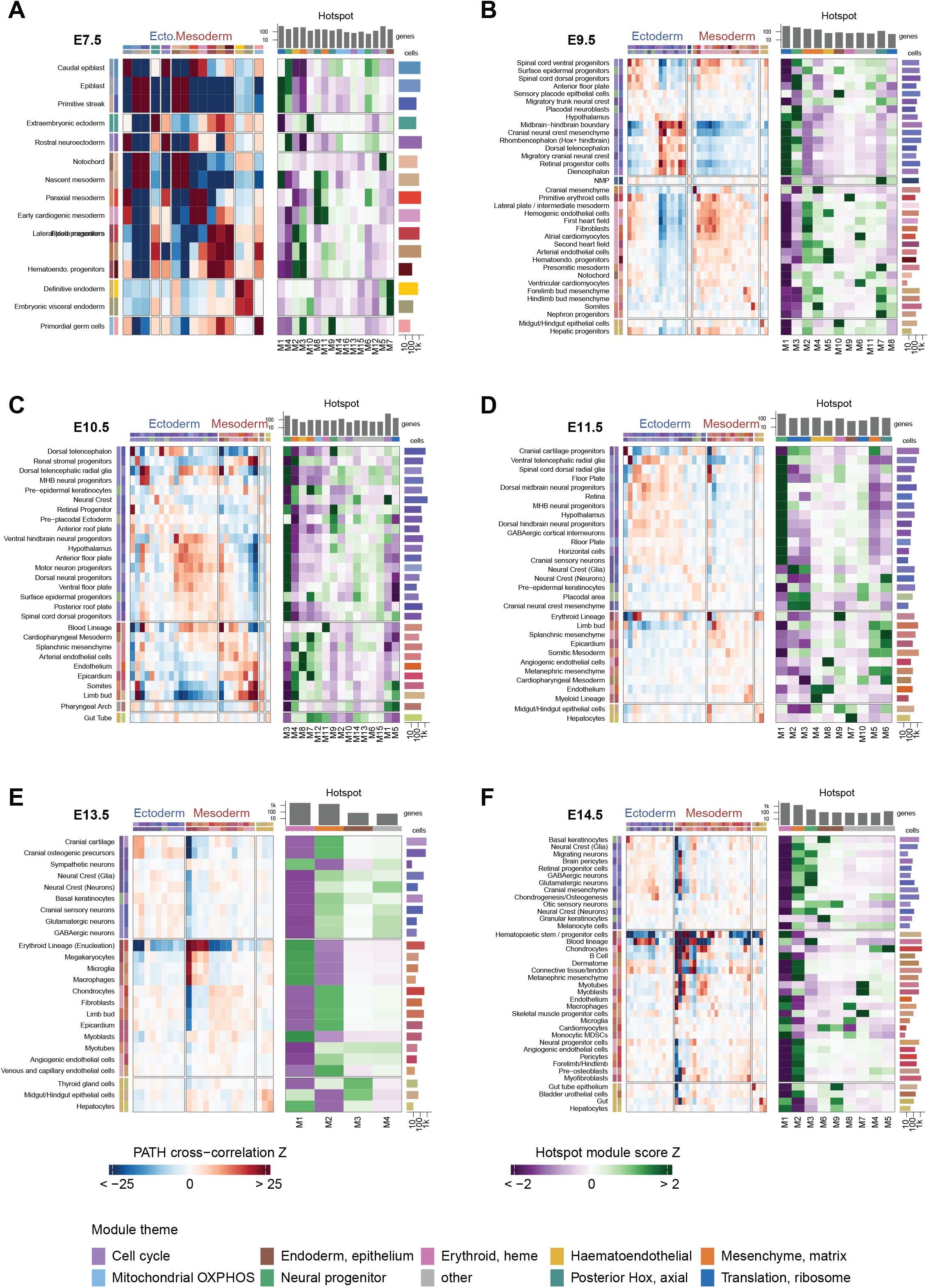
Summary of cell lineage trees across E7.5 to E14.5. **(A–F)** Phylogenetic correlation for cell types calculated from single-cell lineage trees (left) and gene modules grouped based on gene-level phylogenetic correlations and their scores across cell types (right) for E7.5 **(A)**, E9.5 **(B)**, E10.5 **(C)**, E11.5 **(D)**, E13.5 **(E)**, and E14.5 **(F)**. Heatmaps show PATH cross-correlation Z scores and standardized mean Hotspot module scores, respectively. Cell types and module IDs are specific to each stage. Bars show the number of cells per cell type and the number of genes assigned to each module. Module colors identify the functional themes in the key.

**Fig. S16.**
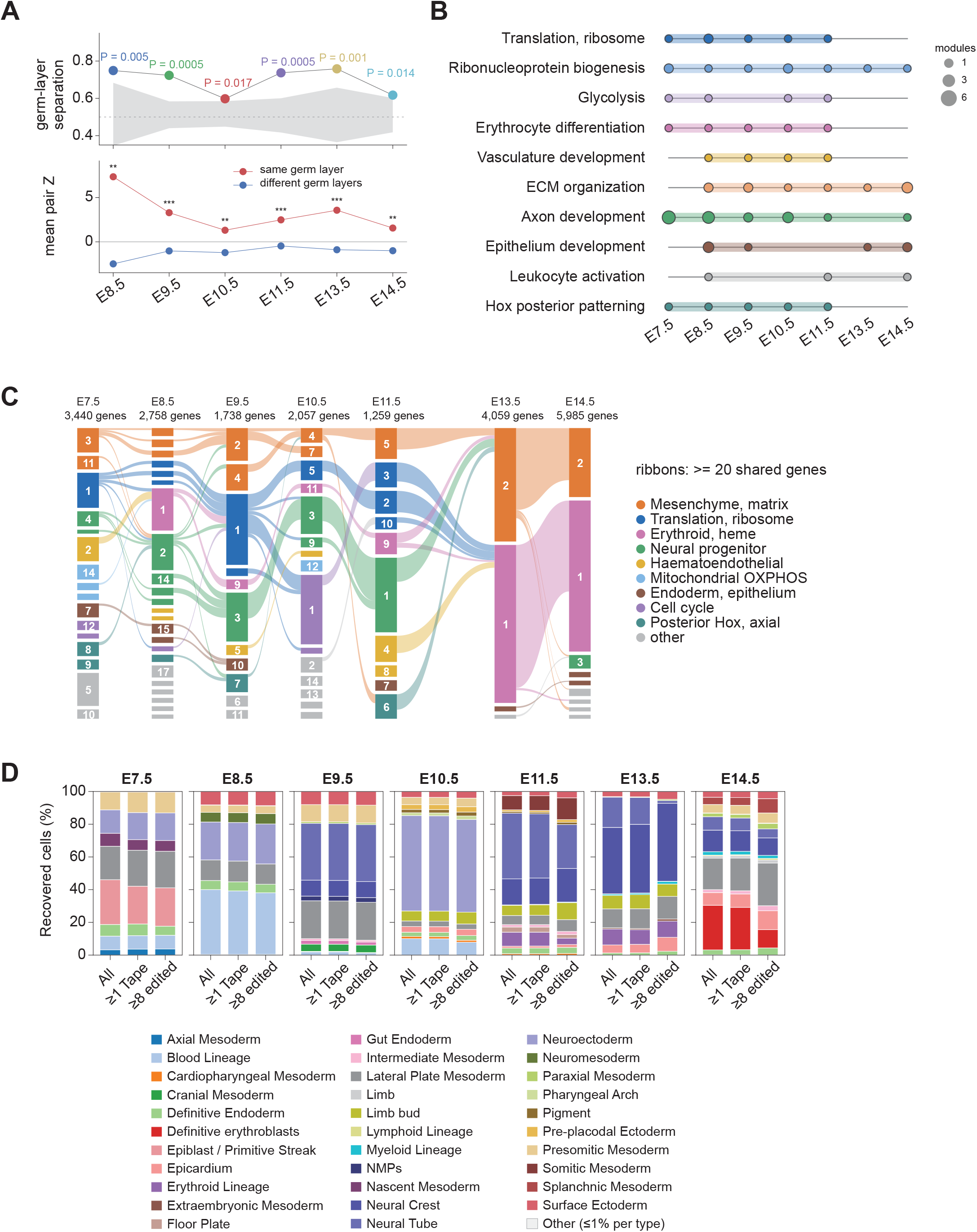
Germ-layer organization and shared transcriptional programs across development. **(A)** Separation of same- and different-germ-layer cell-type pairs by symmetrized PATH scores at E8.5–E14.5, retaining types with ≥20 cells. Upper, AUROC was compared with 2,000 label permutations; gray bands show 95% null intervals. Lower, mean pairwise Z scores; stars denote two-sided Mann–Whitney tests. **(B)** Functional themes represented by lineage-associated Hotspot modules from E7.5 to E14.5. Rows indicate ten themes assigned by Gene Ontology enrichment (**Methods**); columns indicate stages. Dot area denotes the number of modules assigned to each theme at each stage. Colored horizontal segments span the first and last stages with an assigned module. **(C)** Gene overlap between Hotspot modules identified independently at each stage. Blocks represent modules, labeled with stage-specific IDs and colored by functional themes assigned from predefined gene signatures. Ribbons connect modules at adjacent sampled stages sharing ≥20 genes and take the color of the earlier-stage module. Block heights and ribbon widths scale with gene counts; each stage is normalized separately. Numbers above columns give the total genes assigned to modules. **(D)** Germ-layer composition with different filter conditions at each stage: all transcriptome-annotated embryonic cells (All), embryonic cells with at least 1 Tape detected (≥1 Tape), and embryonic cells with at least 8 edited Tapes that are subsequently used for lineage reconstruction (≥8 Tape). Extraembyronic cells are not included.

**Fig. S17.**
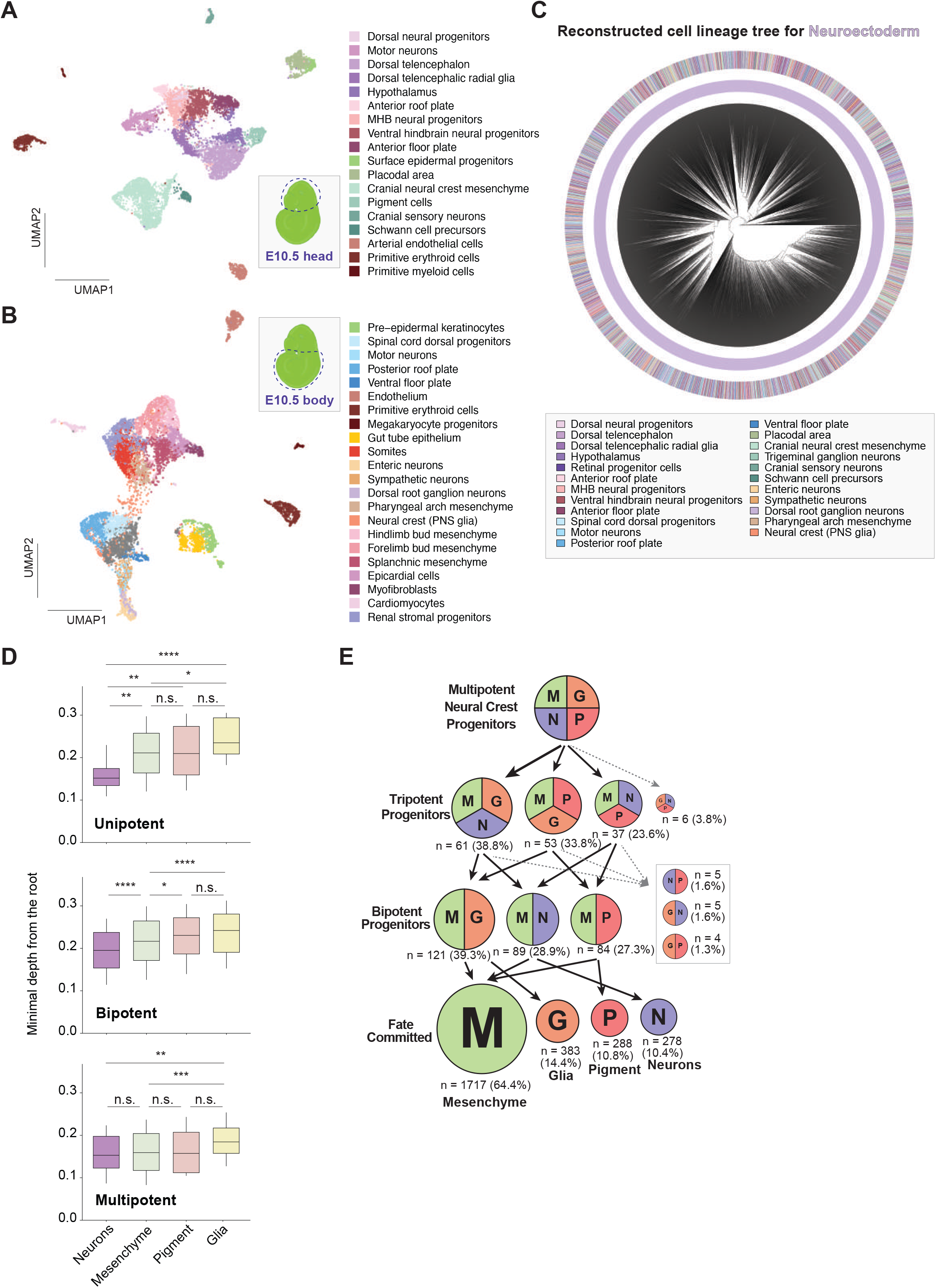
Cell type annotation and observed ancestral fate breadth during neural crest diversification. **(A, B)** UMAP of single cell transcriptomic data for E10.5 embryos, annotated with cell types of the head region in **A**, and the body region in **B**. **(C)** Reconstruction of the lineage tree for the displayed neuroectodermal and neural crest-associated populations in E10.5 mouse embryos. **(D)** Violin plots show the distribution of the minimal depth from the root for shared internal nodes associated with each neural crest derivative class (Mesenchyme, Glia, Neurons, Pigment), stratified by observed descendant fate breadth: Unipotent; nodes with descendants of one derivative class, Bipotent; nodes with descendants of two derivative classes, and Multipotent; nodes with descendants of three or more derivative classes. Colors denote derivative class. Box plots show median and interquartile range, whiskers 10–90%. **(E)** Observed fate sets among inferred ancestral nodes and terminal cells in the E10.5 neural crest-associated tree. Tripotent and bipotent categories contain 157 and 308 nodes, respectively; the terminal layer contains 2,666 cells. M, mesenchyme; G, glia; N, neurons; P, pigment. Counts and percentages show the frequency of each fate set within its displayed category. Arrows schematically connect nested fate sets.

**Fig. S18.**
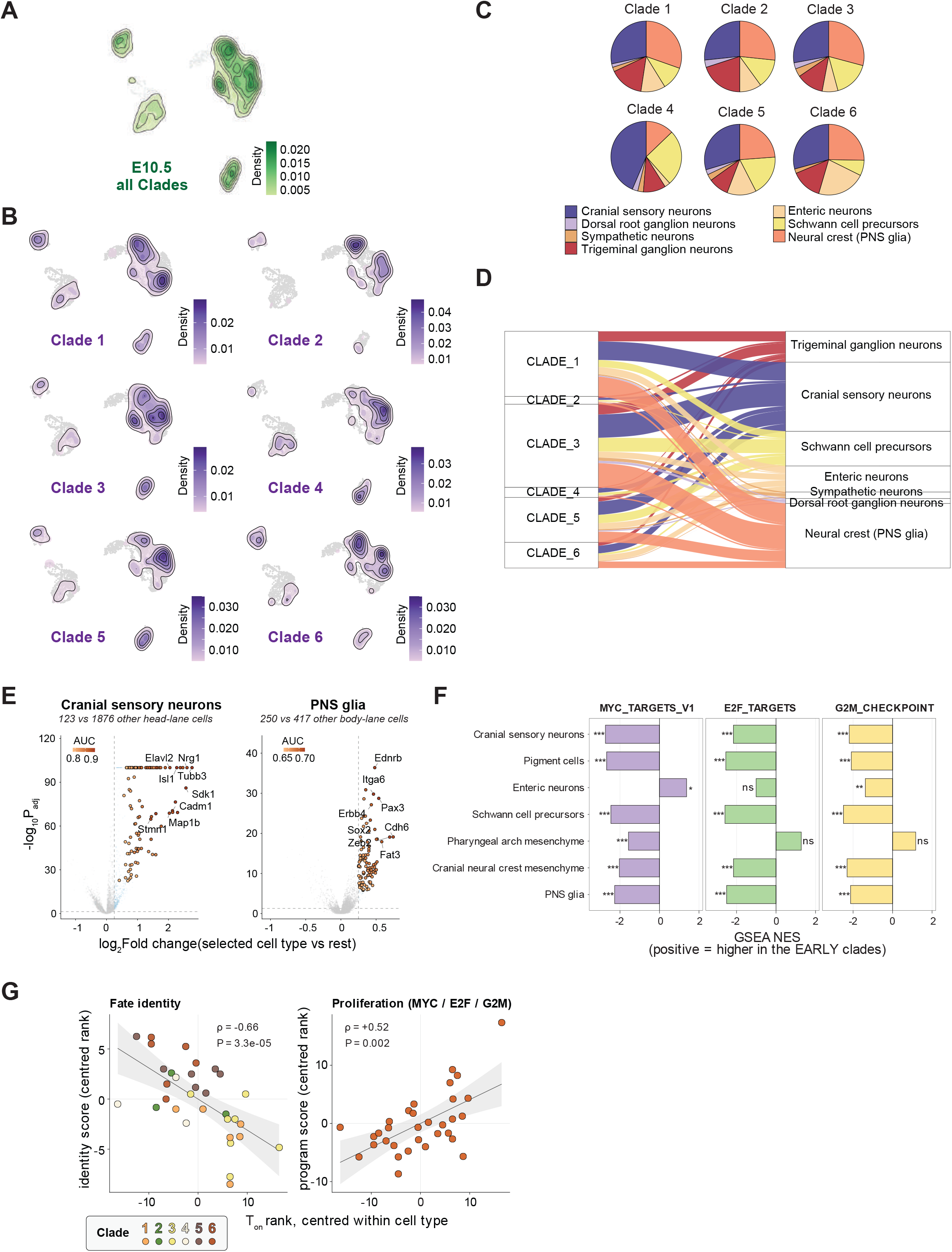
Clade-based stratification links lineage-inferred fate-bias timing to transcriptional programs within neural crest derivatives. **(A)** Density plot of the displayed neuronal and glial neural crest-associated populations at E10.5 in all six clades. The density of the contours reveals cell density in transcriptomic space. **(B)** Density plots of the same displayed neuronal and glial populations across the six clades. Color scales indicate cell density. **(C)** Pie charts showing the distribution of the displayed neuronal and glial populations within each clade. Colors identify the seven cell types in the key. **(D)** Alluvial plots showing the contribution of the displayed neuronal and glial cell types from the six clades. Ribbon widths indicate cell counts; colors denote cell types. **(E)** Identity-marker contrasts for cranial sensory neurons versus other head cells (123 versus 1,876 cells; left) and PNS glia versus other body cells (250 versus 417 cells; right). Volcano plots show log₂ fold change and adjusted significance; colors indicate marker-specificity AUC. **(F)** GSEA of MYC-target, E2F-target, and G2/M-checkpoint programs comparing early- and late-onset clades within each indicated cell type. Positive normalized enrichment scores indicate enrichment in the early-onset clades. **(G)** Within-cell-type centered *T_on_* ranks versus centered ranks of fate-identity scores (left) and proliferation-program scores (right). Each point represents a clade–cell-type combination; colors denote clades. Spearman ρ = −0.66 and 0.52, with P = 3.3 × 10⁻⁵ and 0.002, respectively.

**Fig. S19.**
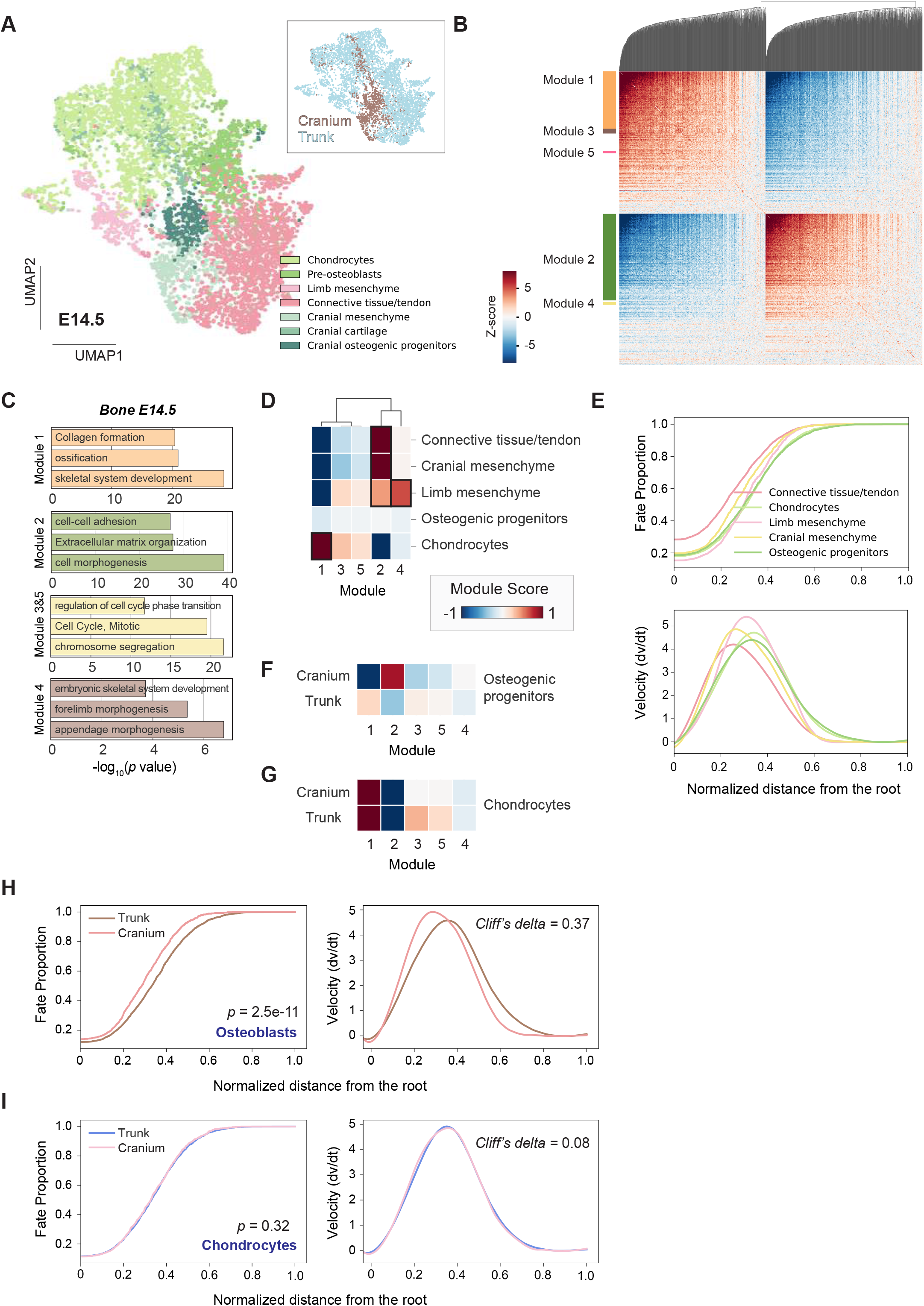
Transcriptional modules capture differentiation programs and regional kinetics in E14.5 skeletal lineages. **(A)** UMAP of seven E14.5 skeletal mesenchymal populations colored by annotated cell types; inset shows anatomical origin assignment (Cranium and Trunk). **(B)** Heatmap of Hotspot local-correlation Z scores, highlighting five transcriptional modules (Modules 1–5). **(C)** Functional enrichment for genes in each module (representative top terms shown). **(D)** Average module scores across major skeletal derivatives (scaled module score). **(E)** Lineage-inferred dynamics for each derivative: mean FateVec trajectories along normalized root-to-tip depth (top) and corresponding derivatives (bottom). **(F)** Regional comparison (Cranium vs Trunk) of module scores within osteogenic populations. **(G)** Regional comparison (Cranium vs Trunk) of module scores within chondrogenic populations. **(H, I)** Developmental kinetics of osteogenic cells **(H)** and chondrocytes **(I)** in Trunk versus Cranium, shown as FateVec trajectories and corresponding derivatives. Two-sided Mann–Whitney tests compare per-tip proportion-curve areas (P = 2.48 × 10⁻¹¹ and 0.323, respectively). Cliff’s delta values for derivative-curve areas are 0.37 and 0.08, respectively.

**Fig. S20.**
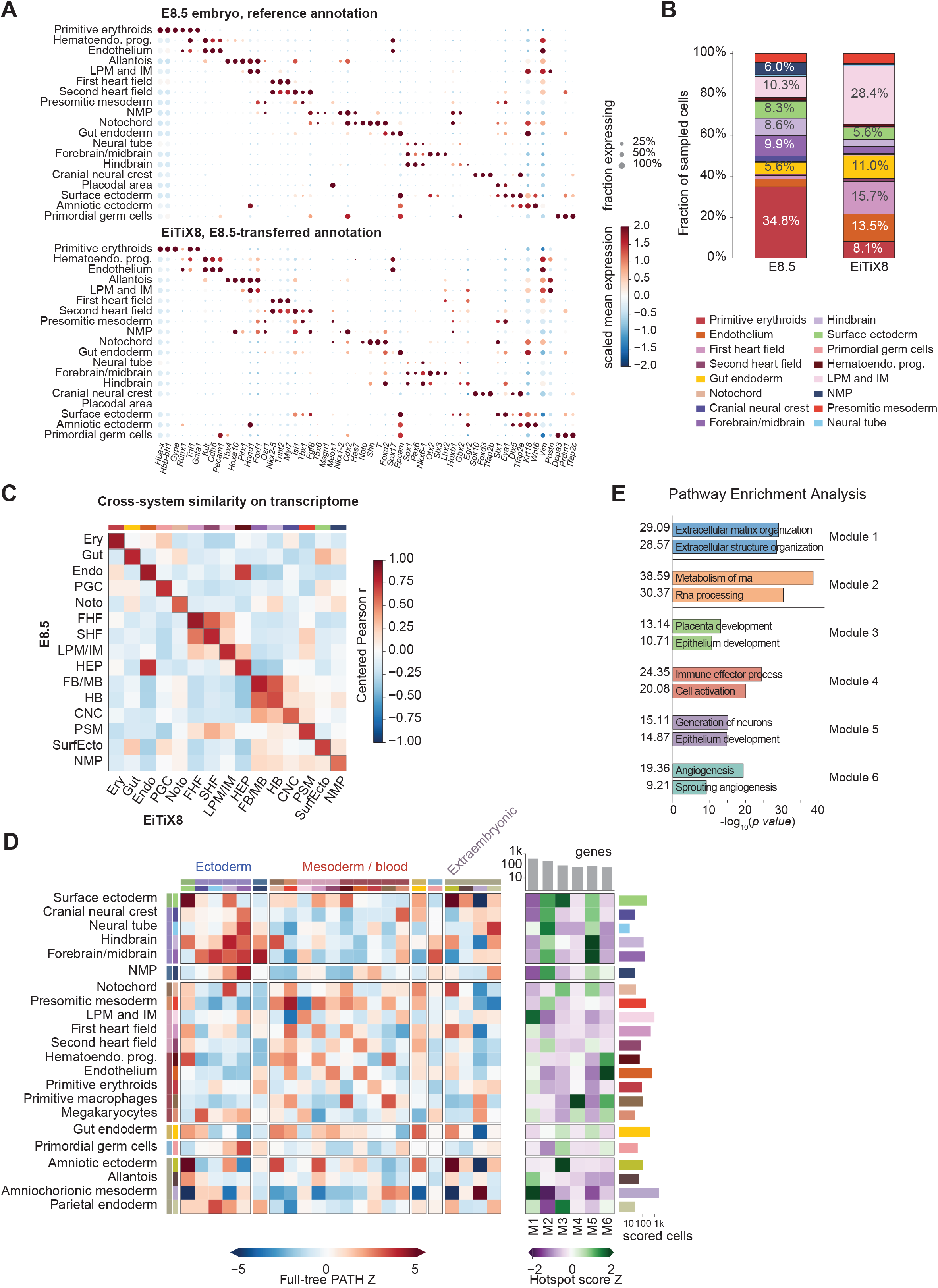
Cell identities and lineage-associated transcriptional programs in EiTiX embryoids. **(A)** Canonical marker expression in E8.5 embryos (Upper) and Day 8 EiTiX-embryoids (Lower). **(B)** Relative abundance of 16 shared cell types in E8.5 (four pooled embryos; 10,860 cells) and EiTiX8 (one profiled sample; 7,694 cells). **(C)** Cross-system transcriptomic similarity for 15 shared cell types. Colors show Pearson correlations between gene-centered cell-type pseudobulk profiles (E8.5, rows; EiTiX8, columns) across 2,740 highly variable genes. **(D)** Lineage associations and gene-module profiles across 22 EiTiX8 cell types. Left, PATH Z scores for within-type (diagonal) and between-type (off-diagonal) associations. Right, mean scores for six Hotspot modules (FDR threshold, 0.25), standardized across modules within each cell type. Bars show cell counts and module gene counts. **(E)** Selected functional enrichments for genes in the six modules in **(D)**, spanning extracellular matrix and structure organization (M1), RNA metabolism and processing (M2), placenta and epithelium development (M3), immune effector processes and cell activation (M4), neuron generation and epithelium development (M5), and angiogenesis (M6).

**Fig. S21.**
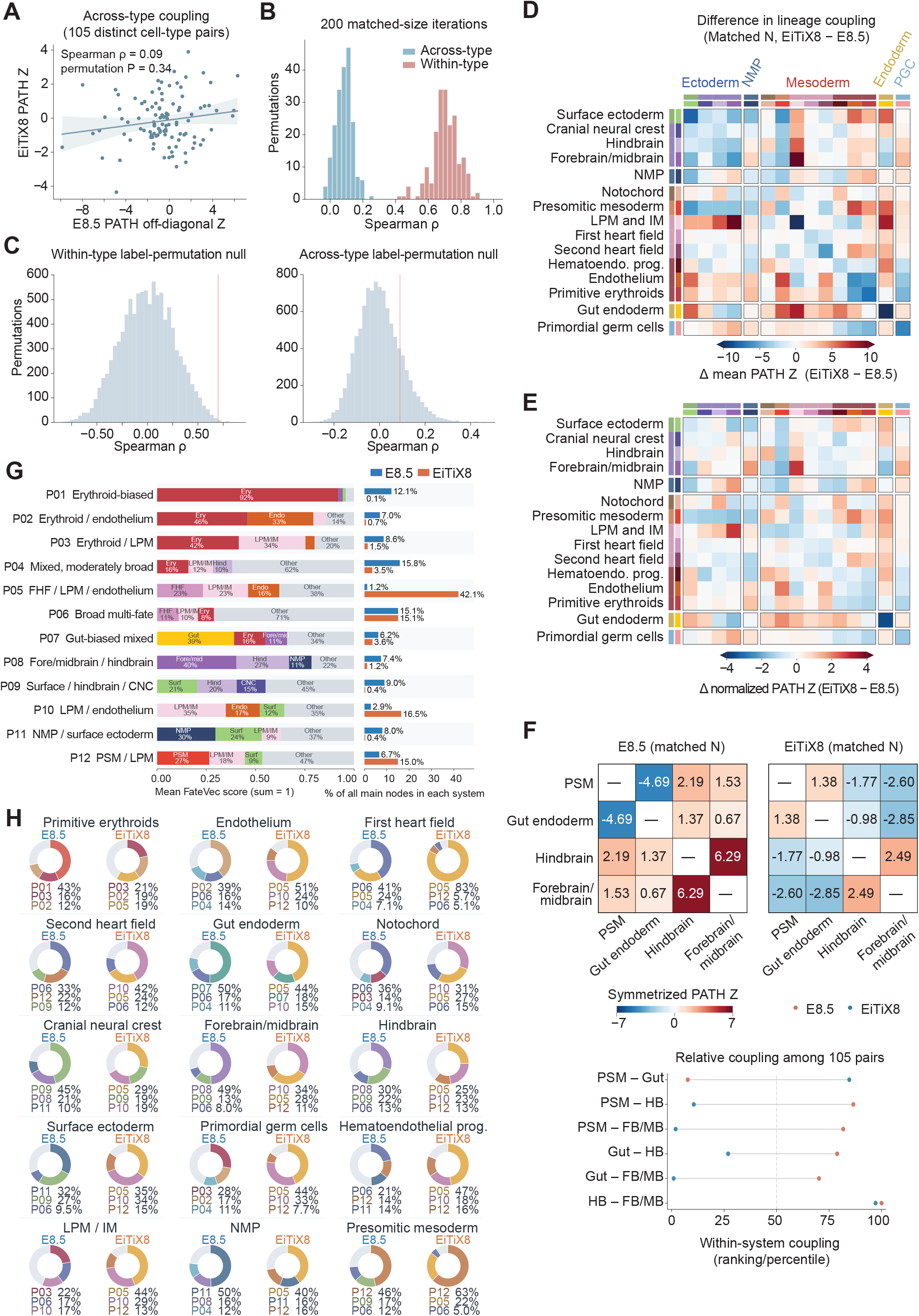
Distinct lineage relationships among shared cell types in embryos and embryoids. Matched cell types permit comparisons of within-type lineage coherence, between-type relationships, and inferred progenitor contributions in embryos and EiTiX embryoids. **(A)** Cross-system agreement of between-type PATH associations across 105 distinct pairs of the 15 shared cell types (Spearman ρ = 0.09; permutation P = 0.34). Each point represents one cell-type pair. **(B,C)** Sampling sensitivity of within-type and across-type agreement over 200 matched subsamples **(B)** and null distributions from 10,000 joint row-and-column label permutations **(C)**. **(D,E)** Differences in lineage organization, expressed as EiTiX-minus-E8.5 PATH scores before **(D)** and after **(E)** standardizing all 225 matrix entries within each system. **(F)** Selected pairwise lineage associations in the matched-cell-count data. Upper, symmetrized PATH Z scores among presomitic mesoderm, gut endoderm, hindbrain, and forebrain/midbrain. Lower, the corresponding percentile ranks among all 105 distinct pairs within each system. **(G)** Mean FateVec profiles and relative frequencies of ancestry-profile groups P01–P12. Fate bars show the three largest components and the remainder; node frequencies use E8.5 and EiTiX group-assigned internal nodes. **(H)** Dominant ancestral node-group associations for each terminal state, assigned by the nearest group-labeled ancestor. The top three groups are shown; gray combines remaining associations, with percentages calculated from all cells in each state.

**Fig. S22.**
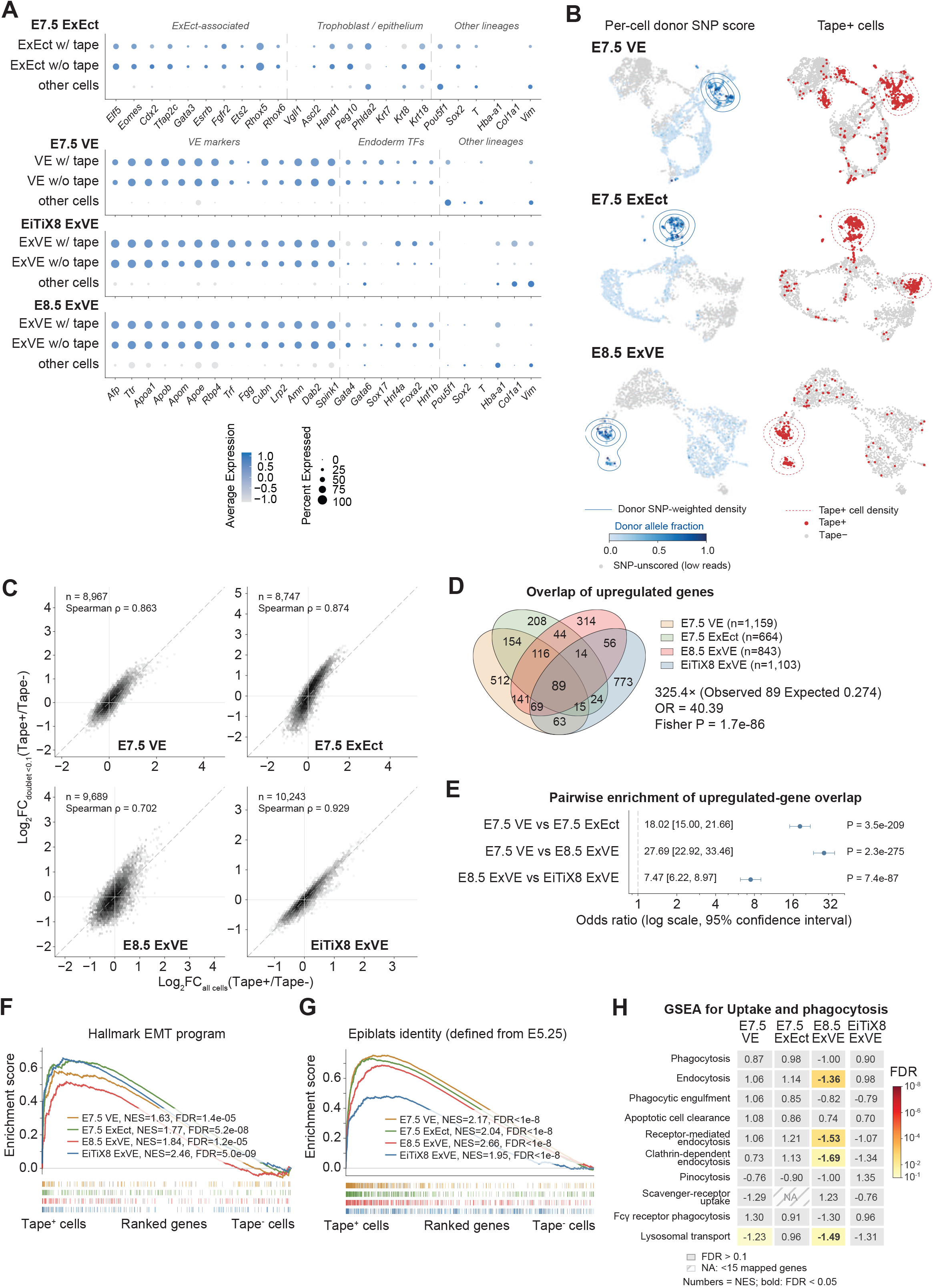
Tape-positive extraembryonic cells share epiblast-associated transcriptional programs. **(A)** Compartment-specific marker expression; dot size indicates expressing fraction and color indicates standardized mean expression. **(B)** Relationship of RNA-SNP donor-allele fractions to Tape-positive-cell distributions in compartment UMAPs. Fractions require ≥5 informative UMI observations across ≥2 of five chromosome-11 genes; gray marks insufficient coverage. Donor-weighted and Tape-positive contours indicate 25, 50, and 75% of each density maximum. **(C)** Sensitivity of Tape-associated expression differences to doublet-score filtering (<0.10), assessed by gene-wise log₂-fold-change concordance. Spearman ρ = 0.863, 0.874, 0.702, and 0.929 for E7.5 VE, E7.5 ExEct, E8.5 ExVE, and EiTiX8 ExVE, respectively. **(D)** Overlap of upregulated genes across the four comparisons, including 89 genes shared by all four sets. The four-set observed/expected ratio is 325.4; OR = 40.39 compares E7.5 VE∩ExEct with E8.5 ExVE∩EiTiX ExVE. **(E)** Pairwise overlap enrichment, shown as odds ratios with 95% log-Wald intervals and one-sided Fisher exact tests. **(F,G)** GSEA of epithelial–mesenchymal transition **(F)** and the independent 500-gene E5.25 epiblast signature **(G)**, plotted against each group’s gene-rank percentile. **(H)** GSEA of uptake- and phagocytosis-associated gene sets. Numbers indicate normalized enrichment scores; positive values indicate enrichment in Tape-positive cells. Colors indicate FDR; bold values have FDR <0.05, and gray indicates FDR >0.10. NA indicates fewer than 15 mapped genes. Tape-positive thresholds are defined in **Fig. 6A**.

**Fig. S23.**
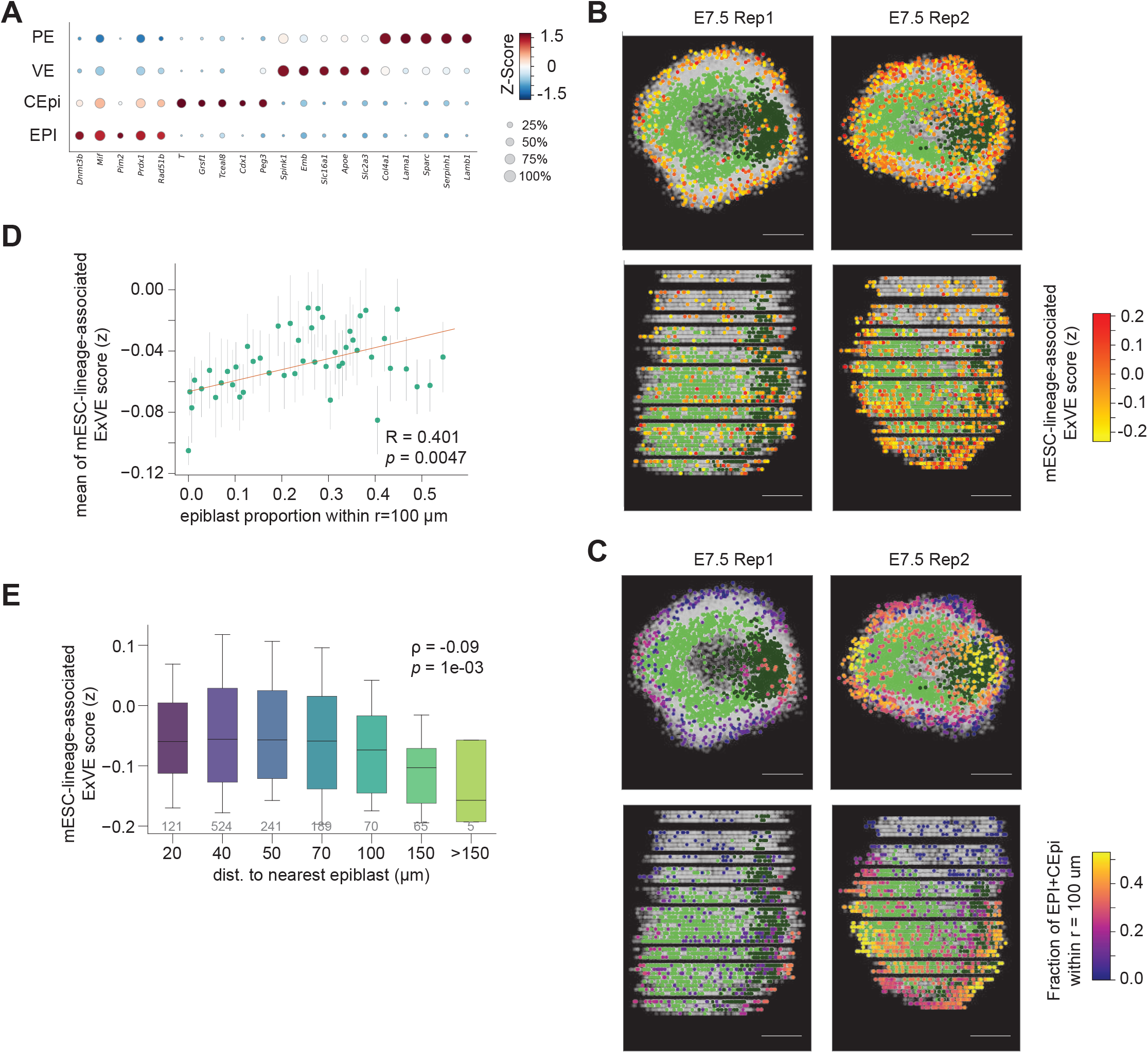
Epiblast neighborhoods and transcriptional variation in extraembryonic endoderm. Spatial transcriptomes test whether the mESC-lineage-associated ExVE signature in VE/PE cells varies with the abundance and proximity of neighboring epiblast cells. **(A)** Marker expression supporting EPI, CEpi, VE, and PE annotations. Dot size indicates the fraction of cells expressing each gene; color indicates standardized mean expression. **(B)** Two spatial projections of each E7.5 replicate. VE/PE cells are colored by the mESC-lineage-associated ExVE score; EPI and CEpi cells are shown in green and dark green, respectively. The score summarizes similarity to the Tape-associated expression state and does not assign lineage origin. **(C)** Local epiblast neighborhood composition in the two E7.5 spatial replicates. VE/PE cells are colored by the fraction of EPI plus CEpi cells within a 100 μm radius. **(D)** Association between local epiblast abundance and the mESC-lineage-associated ExVE score in 143 VE and 1,072 PE cells (350 and 865 cells in the two replicates). Points show mean scores for cells binned by the fraction of EPI plus CEpi neighbors within 100 μm. Pearson r = 0.401, P = 0.0047. **(E)** mESC-lineage-associated ExVE scores in VE/PE cells grouped by distance to the nearest epiblast cell. Numbers below the boxes give cell counts. The cell-level association is weak (Spearman ρ = −0.09, P = 0.001).

## Supplementary table legends

**Table S1.** Summary of scRNA-seq samples, retained cell numbers, and quality-control metrics across developmental stages.

**Table S2.** Lineage-associated gene modules across mouse embryogenesis. Genes assigned to Hotspot modules identified independently at seven developmental stages from E7.5 to E14.5, and EiTiX8. Each row reports the stage, module identifier, gene symbol, gene-level lineage-autocorrelation Z score, and corresponding false discovery rate (FDR). The table contains 22,151 gene–dataset assignments, including 22,320 across mouse developmental stages and 1,024 in EiTiX8; all listed genes have FDR < 0.05 for mouse developmental stages or FDR < 0.25 for EiTiX8. Module identifiers are stage specific and do not imply correspondence between stages.

**Table S3.** Cell-type-specific restriction-onset estimates for 189 cell type–stage combinations across seven stages from E7.5 to E14.5.

**Table S4.** Method comparison, developmental reference times, and cell annotations. Separate worksheets compare FateVec with TarCA, QFM, and Carta; reference germ-layer programs used to calibrate FateVec timing and the mapping from Fig. 2D cell types to Fig. 2E calibration categories in E8.5 cell type annotations. Annotation worksheets cover E7.5, E8.5, E9.5, E10.5, E11.5, E13.5, and E14.5 embryos and day-8 EiTiX embryoids. Cell denotes the recorded cell identifier, germlayer the broad lineage annotation, and subcluster the cell-type annotation.

**Table S5.** Gene signature for spatial analysis of the Tape-associated ExVE program. The 75 genes used to calculate the mESC-lineage-associated ExVE score in whole-mouse-gastrula Stereo-seq datasets.

**Table S6.** Marker genes used to annotate cell types across developmental stages and dissected regions from E7.5 to E14.5.

**Table S7.** Sequences of the 50 TapeBCs for tetraploid complementation and 8 TapeBCs for EiTiX-embryoids identified by bulk genomic DNA amplicon sequencing of DNA Tape arrays.

**Table S8.** Spatially binned signature scores for visceral endoderm (VE) and parietal endoderm (PE) cells in two E7.5 samples, using the genes and scoring procedure in Table S5. Each sample is partitioned into 3, 5, 10, 15, or 20 bins along the section-stacking z axis and directions parallel (AP_axis_along) or perpendicular (AP_axis_across) to an anterior–posterior reference line. The line approximately bisects both epiblast and caudal epiblast distributions; AP_axis_definition records its geometry and orientation. Each bin reports its boundaries, VE and PE counts, landmark-cell counts, and the mean and population SD of scores across VE and PE cells combined.

